# Molecular and cellular dynamics of the developing human neocortex at single-cell resolution

**DOI:** 10.1101/2024.01.16.575956

**Authors:** Li Wang, Cheng Wang, Juan A. Moriano, Songcang Chen, Guolong Zuo, Arantxa Cebrián-Silla, Shaobo Zhang, Tanzila Mukhtar, Shaohui Wang, Mengyi Song, Lilian Gomes de Oliveira, Qiuli Bi, Jonathan J. Augustin, Xinxin Ge, Mercedes F. Paredes, Eric J. Huang, Arturo Alvarez-Buylla, Xin Duan, Jingjing Li, Arnold R. Kriegstein

**Affiliations:** The Eli and Edythe Broad Center of Regeneration Medicine and Stem Cell Research, University of California San Francisco; San Francisco, CA 94143, USA; Department of Neurology, University of California San Francisco; San Francisco, CA 94143, USA; University of Barcelona Institute of Complex Systems; Barcelona, 08007, Spain; Department of Neurological Surgery, University of California San Francisco; San Francisco, CA 94143, USA; Department of Ophthalmology, University of California San Francisco; San Francisco, CA 94143, USA; Neuro-immune Interactions Laboratory, Institute of Biomedical Sciences, Department of Immunology, University of São Paulo; São Paulo, SP 05508-220, Brazil; Department of Physiology, University of California San Francisco, San Francisco, CA 94143, USA; Department of Pathology, University of California San Francisco; San Francisco, CA 94143, USA

## Abstract

The development of the human neocortex is a highly dynamic process and involves complex cellular trajectories controlled by cell-type-specific gene regulation^1^. Here, we collected paired single-nucleus chromatin accessibility and transcriptome data from 38 human neocortical samples encompassing both the prefrontal cortex and primary visual cortex. These samples span five main developmental stages, ranging from the first trimester to adolescence. In parallel, we performed spatial transcriptomic analysis on a subset of the samples to illustrate spatial organization and intercellular communication. This atlas enables us to catalog cell type-, age-, and area-specific gene regulatory networks underlying neural differentiation. Moreover, combining single-cell profiling, progenitor purification, and lineage-tracing experiments, we have untangled the complex lineage relationships among progenitor subtypes during the transition from neurogenesis to gliogenesis in the human neocortex. We identified a tripotential intermediate progenitor subtype, termed Tri-IPC, responsible for the local production of GABAergic neurons, oligodendrocyte precursor cells, and astrocytes. Remarkably, most glioblastoma cells resemble Tri-IPCs at the transcriptomic level, suggesting that cancer cells hijack developmental processes to enhance growth and heterogeneity. Furthermore, by integrating our atlas data with large-scale GWAS data, we created a disease-risk map highlighting enriched ASD risk in second-trimester intratelencephalic projection neurons. Our study sheds light on the gene regulatory landscape and cellular dynamics of the developing human neocortex.

## Main Text

Human neocortex development is a complex and coordinated process crucial for establishing the brain’s intricate structure and functionality. In the developing neocortex, radial glia (RGs) generate glutamatergic excitatory neurons (ENs) in a characteristic inside-out pattern, with deep-layer neurons produced first, followed by upper-layer intratelencephalic (IT) projection neurons^1^. Subsequently, ENs migrate along the radial glial scaffold to the cortical plate, where they differentiate and form distinct cortical layers with coordinated synaptic connections. Meanwhile, GABAergic inhibitory neurons (INs) originating in the ganglionic eminence migrate to the cortex through the marginal and germinal zones, eventually becoming cortical interneurons of the adult cortex. During the late second trimester, RGs transition from neurogenesis to gliogenesis, producing astrocytes and oligodendrocyte lineage cells that populate the cortex. Cell-type-specific gene regulatory mechanisms that underlie cell proliferation and differentiation govern these highly regulated processes. However, our understanding of these mechanisms remains incomplete.

Gene regulation involves epigenetic reprogramming and subsequent gene expression changes^2^. Over the past decade, single-cell transcriptome^3–14^ and chromatin accessibility^11,15–17^ analyses have expanded our knowledge of cellular diversity and the molecular changes that occur during human neocortical development. However, in many instances, measurements of the transcriptome and epigenome were conducted independently, limiting our understanding of how these two modalities coordinate with each other to form regulatory networks in the same cell. Recent studies explored gene-regulatory mechanisms in the developing human cortex by profiling chromatin accessibility and gene expression within the same nuclei^18,19^. However, these analyses were confined either to a restricted number of samples and cell types or to the first trimester, warranting further exploration to obtain a more comprehensive understanding.

In this study, we conducted paired RNA sequencing (RNA-seq) and assay for transposase-accessible chromatin with sequencing (ATAC-seq) on single nuclei from multiple regions and age groups of the developing human neocortex. In addition, spatial transcriptomic analysis was utilized to reveal cellular niches and cell-cell communication. These datasets have enabled the construction of a multi-omic atlas of the human neocortex across different developmental stages at single-cell resolution. Leveraging this atlas, we investigated molecular and cellular dynamics of the developing human neocortex, including cellular composition, spatial organization, intercellular signaling, gene regulatory networks, lineage potential, and disease susceptibility. Our results highlight novel multipotential intermediate progenitor cells (IPCs) and cellular trajectories and shed light on the mechanisms underlying brain cancer and neuropsychiatric disorders.

## Results

### A single-cell multi-omic survey of the developing human neocortex

To characterize transcriptomic and epigenomic changes during human neocortex development, we obtained 27 brain specimens and 38 unique biological samples across five major developmental stages ranging from the first trimester to adolescence, covering key events such as neurogenesis, neuronal migration, gliogenesis, synaptogenesis, and myelination (Fig. 1a, Supplementary Table 1). In addition, we included samples from both the prefrontal cortex (PFC) and primary visual cortex (V1), two poles of the rostral-caudal axis of the neocortex, to understand regional diversity. Applying the single-nucleus multiome (snMultiome) technique from 10x Genomics, we obtained paired single-nucleus ATAC-seq and RNA-seq data from 243,535 nuclei after quality control (see Methods). Some early-stage samples included brain regions other than the neocortex, such as the diencephalon and striatum (Extended Data Fig. 1a–d). We removed non-neocortical nuclei to focus our analysis on the neocortex, resulting in 232,328 nuclei in the final dataset (Supplementary Table 2). We detected similar numbers of genes, transcripts, and ATAC peak region fragments across different samples, with a median of 2289 genes, 4840 transcripts, and 4121 ATAC peak region fragments per nucleus (Extended Data Fig. 2a).

**Fig. 1.**
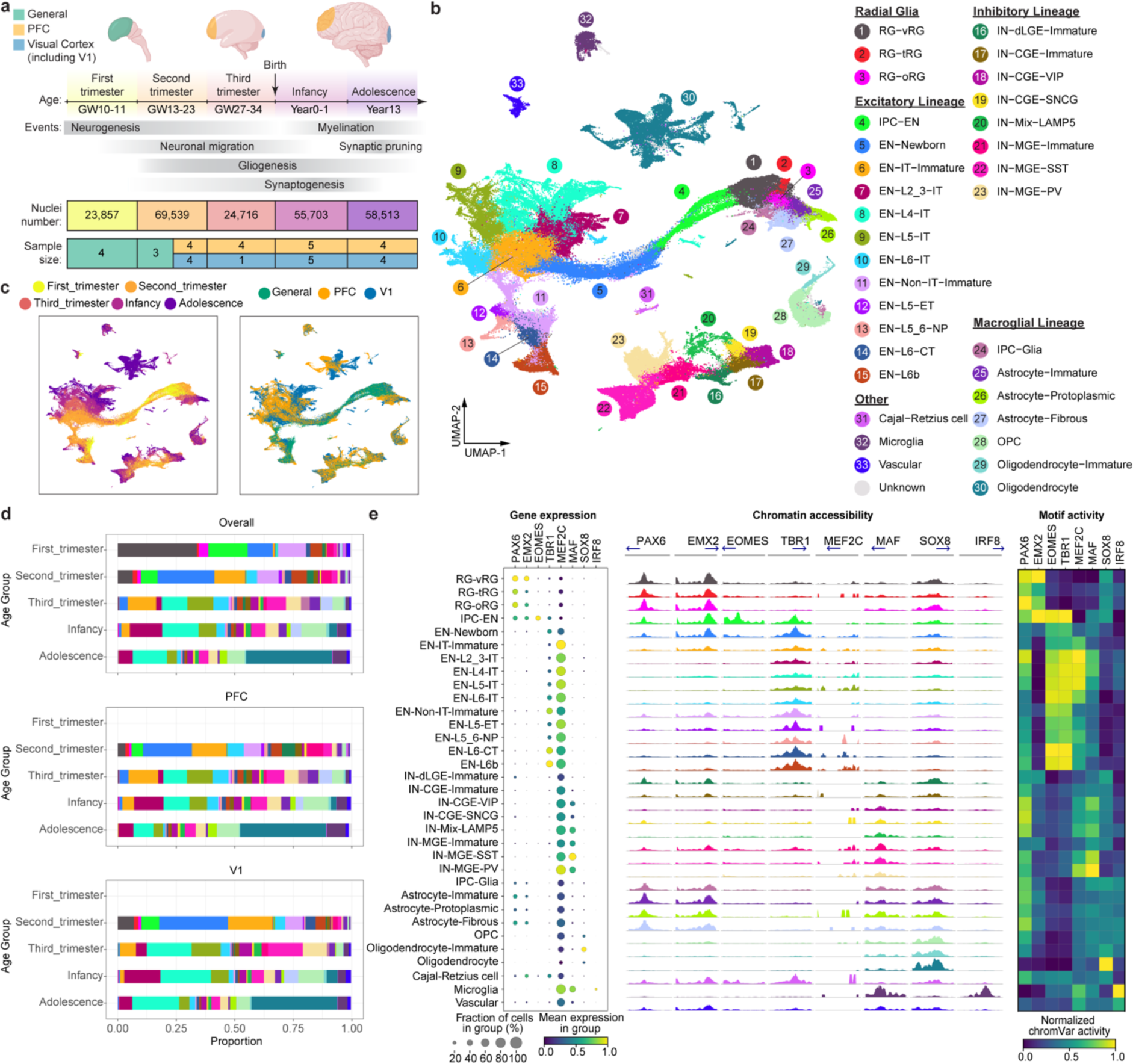
A multi-omic survey of the developing human neocortex. **a**, Description of samples used in this study. **b**, UMAP plots of the snMultiome data showing the distribution of 33 cell types. **c**, UMAP plots showing the distribution of age groups (left) and regions (right). **d**, Proportion of individual cell types across developmental stages and cortical regions. Bars are color-coded by cell types, the legend of which can be found in panel a. **e**, Left, a dotplot of the signature transcriptional factors (TFs) in individual cell types. Middle, aggregated chromatin accessibility profiles on the promoter of signature TFs across cell types. The blue arrow represents each TF’s transcriptional starting site and gene body. Right, heatmap of normalized chromVar motif activity of signature TFs across cell types.

We performed weighted nearest neighbor analysis^20^ to integrate information from the paired ATAC and RNA modalities. The resulting nearest neighbor graph was used for uniform manifold approximation and projection (UMAP) embedding and clustering. We used previously established hierarchical cortical cell-type architecture in the developing and adult human neocortex^14,21^ as references for cluster annotation. Meanwhile, we took into consideration that cell identities can be ambiguous and transient during development. Therefore, we carefully evaluated the expression of marker genes (Extended Data Fig. 3, Supplementary Table 3) and determined 5 classes, 11 subclasses, and 33 high-fidelity cell types (Fig. 1b, Extended Data Fig. 1e, Supplementary Table 2). As expected, cells primarily clustered according to their lineages and, within individual lineages, further clustered by types, age groups, and regions (Fig. 1b,c, Extended Data Fig. 2b). ENs, oligodendrocytes, and astrocytes showed strong regional differences (Fig. 1b,c). By contrast, INs, oligodendrocyte precursor cells (OPCs), microglia, and vascular cells lacked strong region specificity (Fig. 1b,c). Compared with UMAP embeddings based on either ATAC or RNA, embeddings based on both modalities had a more precise separation between cell types, age groups, and regions, suggesting that the combination of both modalities better delineates spatiotemporal cell identities (Extended Data Fig. 2c).

Cell type proportions were comparable between samples of the same age group and region (Extended Data Fig. 2a). However, cell type proportions became substantially different when samples across age groups or regions were compared (Fig. 1d, Supplementary Table 3). Specifically, progenitors (e.g., RG-vRGs [moderated t-test, *P_adj._* = 1.61E−06] and IPC-ENs [*P_adj._* = 9.03E−06]) and immature neurons (e.g., EN-Newborns [*P_adj._* = 9.42E−08] and EN-IT-Immatures [*P_adj._* = 2.48E−09]) were more abundant in the first and second trimester but became depleted at later stages. Conversely, proportions of upper-layer intratelencephalic (IT) neurons (e.g., EN-L2_3-ITs [*P_adj._* = 1.17E−03] and EN-L4-ITs [*P_adj._* = 1.14E−03]) and macroglia (e.g., Astrocyte-Protoplasmic [*P_adj._* = 6.27E−06] and Oligodendrocytes [*P_adj._* = 3.14E−11]) became more abundant after birth. Moreover, EN-L4-ITs were more abundant in V1 than in PFC after the third trimester (*P_adj._* = 1.10E−02), consistent with the expansion of the thalamorecipient layer 4 in V1.

To further evaluate data quality, we compared gene expression, chromatin accessibility, and transcriptional regulatory activities of lineage-specific transcription factors (TFs) across cell types (Supplementary Table 4). We found that the three attributes were concordant with each other at most genomic loci (Fig. 1e). For example, *PAX6* and *EMX2*, two TFs critical for cortical neural progenitor specification^22^, were selectively expressed, had high promoter accessibility, and exhibited enriched motif activities in RGs (Fig. 1e). Similar results were obtained with other lineage-specific TFs. Thus, dynamic changes in epigenome and transcriptome are highly coordinated during human neocortex development.

### Molecularly defined cytoarchitecture of the developing human neocortex

To localize the observed cell types from our snMultiome data, we performed spatial transcriptomic analysis of the developing human neocortex using multiplexed error-robust fluorescence in situ hybridization (MERFISH)^23^. First, guided by the snMultiome data, we designed a 300-gene panel composed of gene markers for the main cell types in the developing cortex (Supplementary Table 5). We then analyzed their expression patterns in PFC and V1 at three age groups from the second trimester to infancy (Supplementary Table 5). From six samples, we retained 404,030 high-quality cells, resulting in 29 cell types that had one-to-one correspondence to those at similar developmental stages in the snMultiome data (Fig. 2a, Extended Data Fig. 4a, Supplementary Table 6). The cell type proportions are comparable between MERFISH and snMultiome data within the same age group, indicating limited sampling bias for both assays (Extended Data Fig. 4b). To determine the cytoarchitecture of the developing neocortex, we defined a cell’s neighborhood as each cell’s 50 closest neighbors. We then unbiasedly divided cells into 10 niches based on the cell type composition of their neighborhoods. The 10 identified niches coincided well with histologically established cortical domains and were thus named after their closest counterpart (Fig. 2a).

**Fig. 2.**
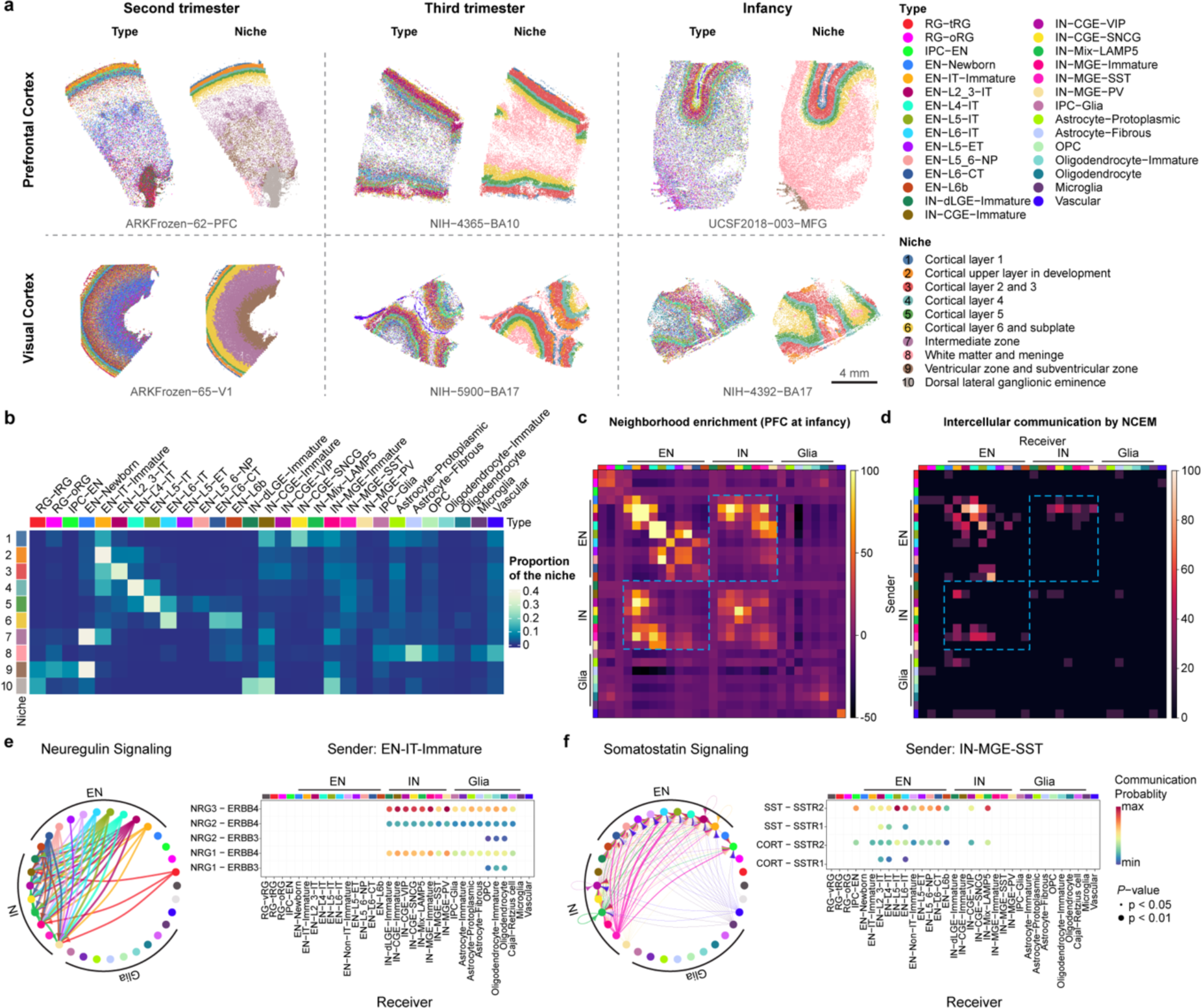
Cell-cell communication in the developing human neocortex. **a**, Spatial transcriptomic analysis of six neocortical samples. Cells are color-coded by types or the niches to which they belong. **b**, Proportion of different cell types in individual niches. Niche numbers correspond to the legend in panel a. **c**, Heatmap showing neighborhood enrichment scores of the PFC sample at infancy. The row and column annotations are color-coded by cell types, the legend of which can be found in panel a. **d**, Heatmap showing the percentage of significant intercellular communication determined by NCEM identified across all datasets. The row and column annotations are color-coded by cell types, the legend of which can be found in panel a. **e**, Left, a circular plot showing the direction of cellular interactions mediated by neuregulin signaling. Right, a dotplot showing communication probability of example ligand-receptor pairs in the neuregulin signaling pathway from EN-IT-Immature to other cell types. Empty space means the communication probability is zero. P-values were calculated by one-sided permutation test. **f**, Left, a circular plot showing the direction of cellular interactions mediated by somatostatin signaling. Right, a dotplot showing communication probability of example ligand-receptor pairs in the somatostatin signaling pathway from IN-MGE-SST to other cell types. Empty space means the communication probability is zero. P-values were calculated by one-sided permutation test.

Different cell types exhibited distinct patterns of niche distribution. Neural progenitors were primarily localized in the ventricular/subventricular zone (VZ/SVZ), whereas mature ENs were confined to their specific cortical layers throughout development (Fig. 2b, Extended Data Fig. 5a–f). Immature interneurons in the second trimester were enriched in both the marginal zone and VZ/SVZ, two routes they use to migrate into the cortex^24^. In the second trimester, the overall ratio of migrating interneurons in the marginal zone to VZ/SVZ was 1:4.1. Interestingly, this ratio was 1:3.3 for caudal ganglionic eminence (CGE)-derived interneurons and 1:5.2 for medial ganglionic eminence (MGE)-derived interneurons (odds ratio = 1.58, *P* < 2.2E−16, two-sided Fisher’s exact test), demonstrating lineage-specific preference in migration routes. Immunostaining using independent samples further validated this observation (Extended Data Fig. 6a,b, weighted average odds ratio = 1.56). This bias may contribute to the laminar distribution of interneuron subtypes at later stages, with CGE-derived interneurons enriched in upper layers and IN-MGE-PVs enriched in layers 4–6 (Fig. 2a,b, Extended Data Fig. 5a–f). Notably, in the developing mouse cortex, biases in tangential migratory route choices based on interneuron identities have been observed^25^. However, unlike our observations in humans, there were no significant differences between the overall MGE- and CGE-derived IN populations in mice. The dorsal lateral ganglionic eminence (dLGE) primarily gives rise to olfactory bulb interneurons^26^. Interestingly, we observed immature INs expressing *MEIS2*, *SP8*, *TSHZ1*, and *PBX3*, presumably originating from dLGE (IN-dLGE-Immatures), in the white matter across all three age groups (Extended Data Fig. 5a–f). These neurons will likely constitute a subset of the white matter interstitial GABAergic interneurons in adulthood. Regarding glial cells, OPCs were evenly distributed between gray and white matter from the second trimester to infancy. However, oligodendrocytes were predominantly present in the white matter for all three age groups (Fig. 2b, Extended Data Fig. 5a–f). This difference supports a non-progenitor role of OPCs in cortical gray matter^27^. Microglia were highly enriched in the white matter (Fig. 2b, Extended Data Fig. 5a–f), consistent with their spatial distribution in the adult brain^28^.

In early neonatal and adult mammalian brains, neurogenesis continues in the VZ/SVZ of the lateral ventricles, and the interneurons produced migrate to the olfactory bulb^29^. Most of these olfactory bulb interneurons are GABAergic, but some could be glutamatergic^30^. We examined our perinatal PFC sample, which contained VZ/SVZ. We found a surprisingly large number of glutamatergic EN-Newborns, along with a small number of IPC-ENs, specifically within the SVZ (Extended Data Fig. 5c). Remarkably, within the VZ/SVZ of this sample, the count of EN-Newborns was 10.3-fold higher than that of IN-dLGE-Immatures, which are considered putative newborn GABAergic olfactory bulb interneurons. Whether these late-born EN-Newborns will migrate to the cortical gray matter, the subcortical white matter, or the olfactory bulb remains to be determined.

### Cell-cell communication in the developing human neocortex

To identify cell-cell communication in the developing human neocortex, we first evaluated the spatial proximity of cell types in each MERFISH sample through neighborhood enrichment analysis. We found that different types of ENs were enriched in their own neighborhoods, consistent with their strong layer specificity. Interestingly, we also observed robust neighborhood enrichment between specific types of ENs and INs, such as EN-IT-Immatures and IN-CGE-VIPs, as well as EN-L4-ITs and IN-MGE-SSTs (Fig. 2c, Extended Data Fig. 7a). To determine if the gene expression of a cell type was influenced by its proximity to a neighboring cell type, we performed node-centric expression modeling (NCEM)^31^. Cell communication inference via NCEM revealed strong interactions among various types of ENs and between ENs and INs across multiple datasets (Fig. 2d, Extended Data Fig. 7b, Supplementary Table 7). Notably, EN-IT-Immatures (sender) affected gene expression in various IN types (receivers). In contrast, IN-MGE-SSTs (sender) influenced gene expression in multiple EN types (receivers).

Since most of the MERFISH samples were collected from stages preceding the peak of synaptogenesis in humans, we resorted to ligand-receptor analysis using CellChat^32^ to identify potential mechanisms underlying the communication between ENs and INs (Extended Data Fig. 7c). Focusing on EN-IT-Immatures and IN-MGE-SSTs as ligand producing cells, we found that neuregulin and somatostatin were potential mediators for their communication with INs and ENs, respectively (Fig. 2e,f, Supplementary Table 8). To explore the role of somatostatin signaling in EN differentiation, we treated midgestational human cortical organotypic slice cultures with two different somatostatin receptor agonists. We then performed single-cell RNA sequencing (scRNA-seq) to analyze gene expression changes in individual EN subtypes (Extended Data Fig. 8a,b, Supplementary Tables 9 and 10). Both agonists inhibited neuron projection development and synaptogenesis while activating multiple metabolic processes, effects observed consistently across multiple EN subtypes (Extended Data Fig. 8c, Supplementary Tables 11 and 12). These results suggest that somatostatin produced by IN-MGE-SST regulates EN maturation. Together, our findings highlight the reciprocal communications between the two major neuronal subclasses during human cortical development.

### Gene regulatory networks in the developing human neocortex

To establish the gene regulatory networks (GRNs) governing human neocortical development, we employed SCENIC+^33^, a computational framework that combines single-cell ATAC and gene expression data with motif discovery to infer enhancer-driven regulons (eRegulons), linking individual TFs to their respective target cis-regulatory regions and genes. Our analysis identified 582 eRegulons, comprising 385 transcriptional activators and 197 repressors (Supplementary Table 13). These eRegulons collectively targeted 8134 regions and 8048 genes. To validate the predicted eRegulons, we evaluated the overlap between eRegulon-predicted target regions and ChIP-seq data of the corresponding TFs from human neocortex^34^. We found that 79% of the tested TFs exhibited higher-than-expected overlap, with 58% showing significant enrichment (Extended Data Fig. 9a, Supplementary Table 14). Additionally, the predicted enhancer-to-gene connections were significantly enriched in enhancer-promoter loops identified through 3D genome profiling of the developing human neocortex^35^ (Extended Data Fig. 9b, Supplementary Table 14, odds ratio = 2.47, P value = 1.1E-7). These findings support the validity of the identified eRegulons.

We quantified the activity of each eRegulon in each nucleus using the AUCell algorithm^36^, assessing region-based and gene-based AUC scores according to the overall accessibility of target enhancers and expression levels of target genes, respectively. Consistent with expectations, expression levels of transcriptional activators exhibited a positive correlation with the AUC scores of their target regions and genes, whereas transcriptional repressors negatively correlated with their targets (Extended Data Fig. 9c). Focusing on activators, we not only recovered established master regulators of cortical progenitors (e.g., *EMX1* and *SALL1*), ENs (e.g., *FOXP1* and *TBR1*), INs (e.g., *ARX* and *LHX6*) but also uncovered novel cell-type- and age-specific eRegulons that potentially serve as lineage-determining factors (Fig. 3a, Extended Data Fig. 9d, Supplementary Table 15).

**Fig. 3.**
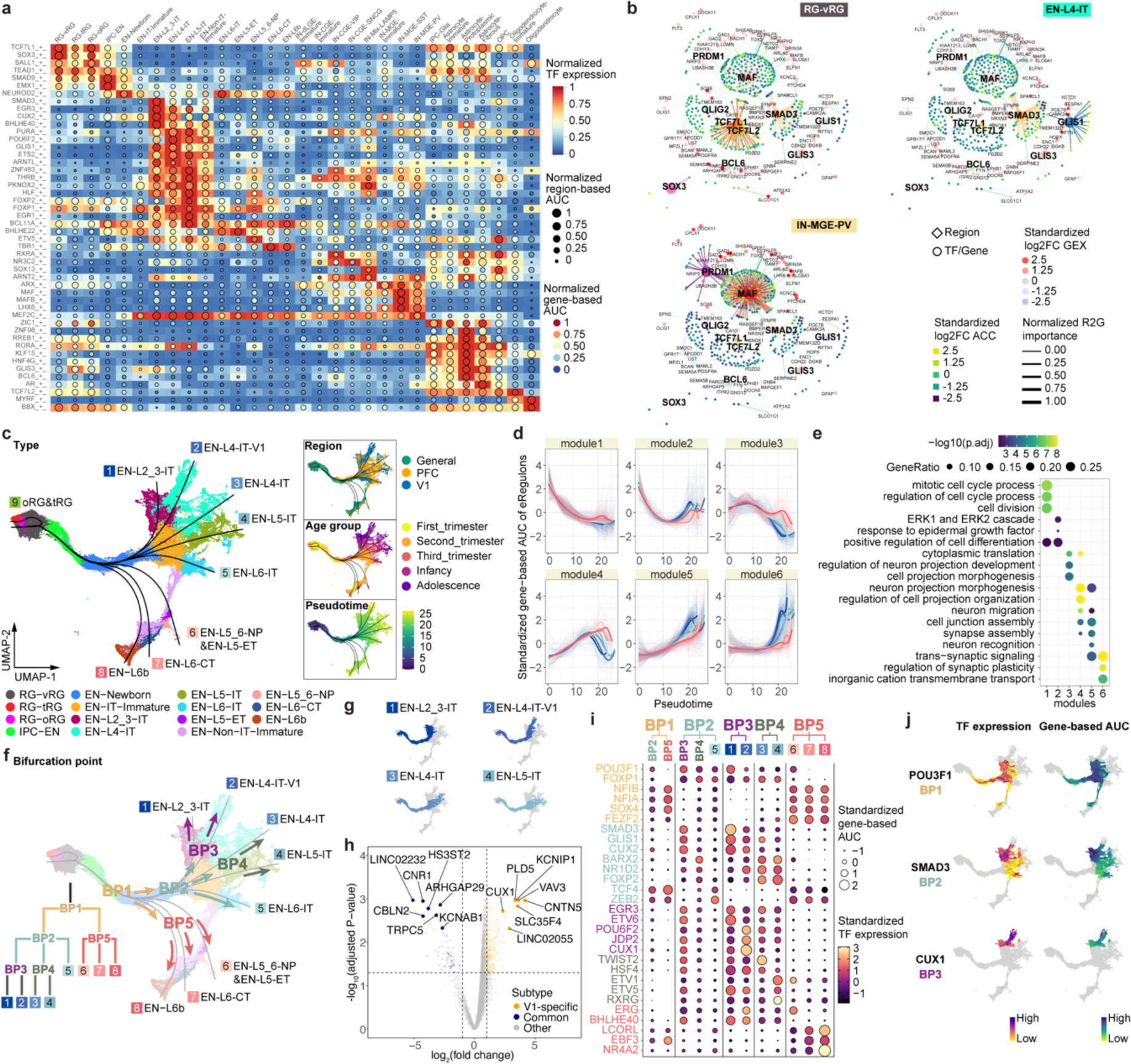
Gene regulatory networks that establish cell identities. **a**, A heatmap-dotplot showing the min-max normalized TF expression levels, region-based AUC scores, and gene-based AUC scores of selective eRegulons across cell types. **b**, Gene regulatory networks of selective eRegulons in three distinct cell types (RG-vRG, EN-L4-IT, and IN-MGE-PV). TF nodes and their links to enhancers are individually colored. The size and the transparency of the TF nodes represent their gene expression levels in each cell type. **c**, UMAP plots of cells belonging to excitatory neuron lineages showing the nine trajectories. Cells are color-coded by types, regions, age groups, or pseudotime. **d**, Standardized gene-based AUC scores of six eRegulon modules along the trajectories of excitatory neuron lineages. eRegulons are color-coded by neuronal trajectories. Thick, non-transparent lines represent the average AUC scores of each module in each trajectory. **e**, Gene ontology enrichment analysis for target genes of individual eRegulon modules. Empty space means adjusted P values > 0.05. One-sided hypergeometric test; nominal P values were adjusted by the Benjamini and Hochberg method. **f**, Bifurcation points during excitatory neuron differentiation. **g**, Trajectories of four intratelencephalic neuron lineages. **h**, Volcano plots highlighting differentially expressed genes between V1-specific and common EN-L4-IT neurons. Likelihood ratio test; nominal P values were adjusted by the Benjamini and Hochberg method. **i**, A dotplot highlighting representative eRegulons (activators) involved in trajectory determination at bifurcation points. **j**, UMAP plots highlighting representative eRegulons involved in trajectory determination at bifurcation points.

In addition, we observed that many cell-type-specific eRegulons shared target regions and target genes (Extended Data Fig 10a). Notable instances included *TCF7L1* and *TCF7L2* in RG-vRGs, *GLIS1* and *SMAD3* in EN-L4-ITs, *MAF* and *PRDM1* in IN-MGE-PVs, *PAX6* and *SOX9* in Astrocyte-Protoplasmics, as well as *OLIG2* and *VSX1* in OPCs (Fig. 3b, Extended Data Fig. 10b). These cooperative TFs exhibit three modes of action: they share the same motif and binding sites (Extended Data Fig. 10c), they bind in tandem at the same enhancer (Extended Data Fig. 10d), or they target different enhancers but converge on the same target gene (Extended Data Fig. 10e,f). The cooperative sharing of regulatory targets likely serves to increase the specificity and robustness of GRNs during cortical development^37,38^.

### Genetic programs that determine excitatory neuron identities

Having established the GRNs, we sought to understand how the activation of cell-type-specific eRegulons controls cortical neuron differentiation. To this end, we selected nuclei from EN lineages, inferred nine differentiation trajectories originating from RG-vRG, and calculated pseudotime values for each nucleus (Fig. 3c, Extended Data Fig. 11a–f, Supplementary Table 16)^39^. Except for one trajectory leading to late-stage radial glia (oRG and tRG), the remaining eight trajectories ended with terminally differentiated ENs. Utilizing a generalized additive model^40^, we analyzed eRegulon activity along each trajectory, categorizing all eRegulons into six modules based on their temporal patterns of activity (Fig. 3d, Supplementary Table 17). Overall, all six modules exhibited distinct activity patterns along the pseudotime but comparable patterns across trajectories (Fig. 3d). Modules specifically active in the early, intermediate, and late stages respectively promoted cell division, cell projection morphogenesis, and synaptic plasticity (Fig. 3e, Supplementary Table 17). These findings highlight that most eRegulons demonstrate conserved activity across various types of ENs, governing shared cellular processes during neuronal differentiation.

Our subsequent objective was to explore gene regulatory mechanisms that determine EN identities. To achieve this, we pinpointed five bifurcation points (BPs) along the eight differentiation trajectories (Fig. 3f). An intriguing finding emerged regarding EN-L4-ITs, which delineated into two distinct trajectories based on their region of origin (Fig. 3c,f). Specifically, the divergence occurred at BP2, where V1-specific EN-L4-ITs continued their trajectory alongside EN-L2_3-IT, while the EN-L4-ITs shared between PFC and V1 followed a trajectory partially overlapping with EN-L5-IT (Fig. 3f,g). To further discriminate between the two EN-L4-IT subtypes, we performed differential gene expression analysis, identifying 1,908 differentially expressed genes between V1-specific and common EN-L4-ITs (Fig. 3h, Extended Data Fig. 12a,b, Supplementary Table 18). We then examined the expression patterns of top differentially expressed genes using in situ hybridization (ISH) data from Allen Brain Atlas. Notably, *CUX1* and *KCNIP1*, two genes preferentially expressed in V1-specific EN-L4-IT, exhibited stronger ISH signals in layer 4 of V1 compared to the adjacent secondary visual cortex (V2) (Extended Data Fig. 12c). In contrast, the common EN-L4-IT biased gene *KCNAB1* showed robust and specific signals in layer 4 of V2 but only displayed scattered signals in V1 (Extended Data Fig. 12c). Moreover, both V1-specific and common EN-L4-ITs expressed markers of their counterparts recently reported in the adult human cortex^41^ (Extended Data Fig. 12d). These findings confirm the presence of V1-specific EN-L4-ITs in the developing neocortex and underscore their distinct developmental trajectory compared to EN-L4-ITs found in other cortical regions.

To identify eRegulons associated with lineage bifurcation, we segmented the differentiation trajectories into five parts and conducted trajectory-based differential eRegulon activity analysis within specific segments encompassing each BP (Extended Data Fig. 11g, Methods). Among the top-ranked differentially active eRegulons at BPs were those featuring well-established TFs crucial for cell identity acquisition, including *CUX2* for upper-layer IT neurons, *FEZF2* for non-IT neurons, and *NR4A2* for EN-L6bs (Fig. 3i, Supplementary Table 19). Furthermore, our analysis revealed novel candidate regulators at multiple levels of lineage bifurcation, such as *POU3F1* for IT neurons, *SMAD3* for upper-layer IT neurons, and *CUX1* for V1-specific EN-L4-ITs, among many others (Fig. 3i,j, Extended Data Fig. 11h). Collectively, these results reveal genetic programs that control the divergence of EN identities.

### Lineage potential of glial progenitors in the late second trimester

Between gestational week (GW) 18 and 26, RGs in the human neocortex gradually transition from neurogenesis to gliogenesis^42^. However, our understanding of gliogenesis in the human neocortex is still limited compared to neurogenesis. In the snMultiome dataset, we identified a total of 10 different cell types within the macroglia lineage, including three RGs types, IPC-Glia, and other cell types associated with either the astrocyte or oligodendrocyte lineages (Extended Data Fig. 13a,b). Among these cell types, *EGFR*^high^*OLIG2*^+^ IPC-Glia has been previously reported by us and others as “pre-OPC”^43^, “pri-OPC”^44^, “mGPC”^16^, “bMIPC”^45^, “gIPC”^10^, or “GPC”^46^ in humans. A similar cell type has been noted in mice as “pri-OPC”^47^, “tri-IPC”^48^, or “MIPC”^49^. Studies using human tissue have demonstrated IPC-Glia’s capacity to generate OPCs ^43^ and astrocytes^46^. Moreover, genetic labeling experiments in mice suggested their additional potential to produce olfactory bulb interneurons^48^. Despite these advancements, ongoing debates and uncertainties persist regarding the origin and lineage potential of human glial progenitors, especially in the late second trimester, when various glial progenitor types emerge.

To address this uncertainty, we leveraged our snMultiome data collected between GW20 and GW24 and explored the expression patterns of surface protein markers (Extended Data Fig. 13c.d). We identified five proteins whose combinatorial expression effectively distinguishes between different glial cell types in the late second trimester (Fig. 4a, Extended Data Fig. 13e). Employing tissue dissection, surface protein staining, and fluorescence-activated cell sorting, we isolated four different glial progenitors–RG-tRGs, RG-oRGs, IPC-Glia, and OPCs (Fig. 4b, Extended Data Fig. 13f) from the late second-trimester human cortex. We first assessed the isolated cells morphologically after culturing for five days in basal culture medium without growth factor supplement (Fig. 4b). RG-tRGs and RG-oRGs were mostly unipolar, featuring a large soma and a thick, long primary process akin to the radial fiber. IPC-Glia appeared mostly bipolar or oligopolar, with shorter processes compared to RGs. OPCs exhibited a “bushy” morphology, suggesting they had started differentiating into pre-myelinating oligodendrocytes. Most cells in the OPC culture died within eight days, consistent with their dependence on specific growth factors for survival^46^. Thus, our subsequent analysis focused on the remaining three progenitor types. We immunostained the sorted cells on day one in vitro (DIV1) to validate their identities (Extended Data Fig. 14a–e). Isolated RG-tRGs and RG-oRGs were positive for the progenitor marker, TFAP2C, whereas the tRG marker, CRYAB, was specifically expressed in RG-tRGs. In contrast, IPC-Glia were positive for both OLIG2 and EGFR. Few cells across all three cultures displayed positivity for the EN marker, NeuN, the astrocyte marker, SPARCL1, or the IN lineage marker, DLX5. In addition, few cells were OLIG2^+^ only, suggesting minimum contamination from OPCs or oligodendrocytes.

**Fig. 4.**
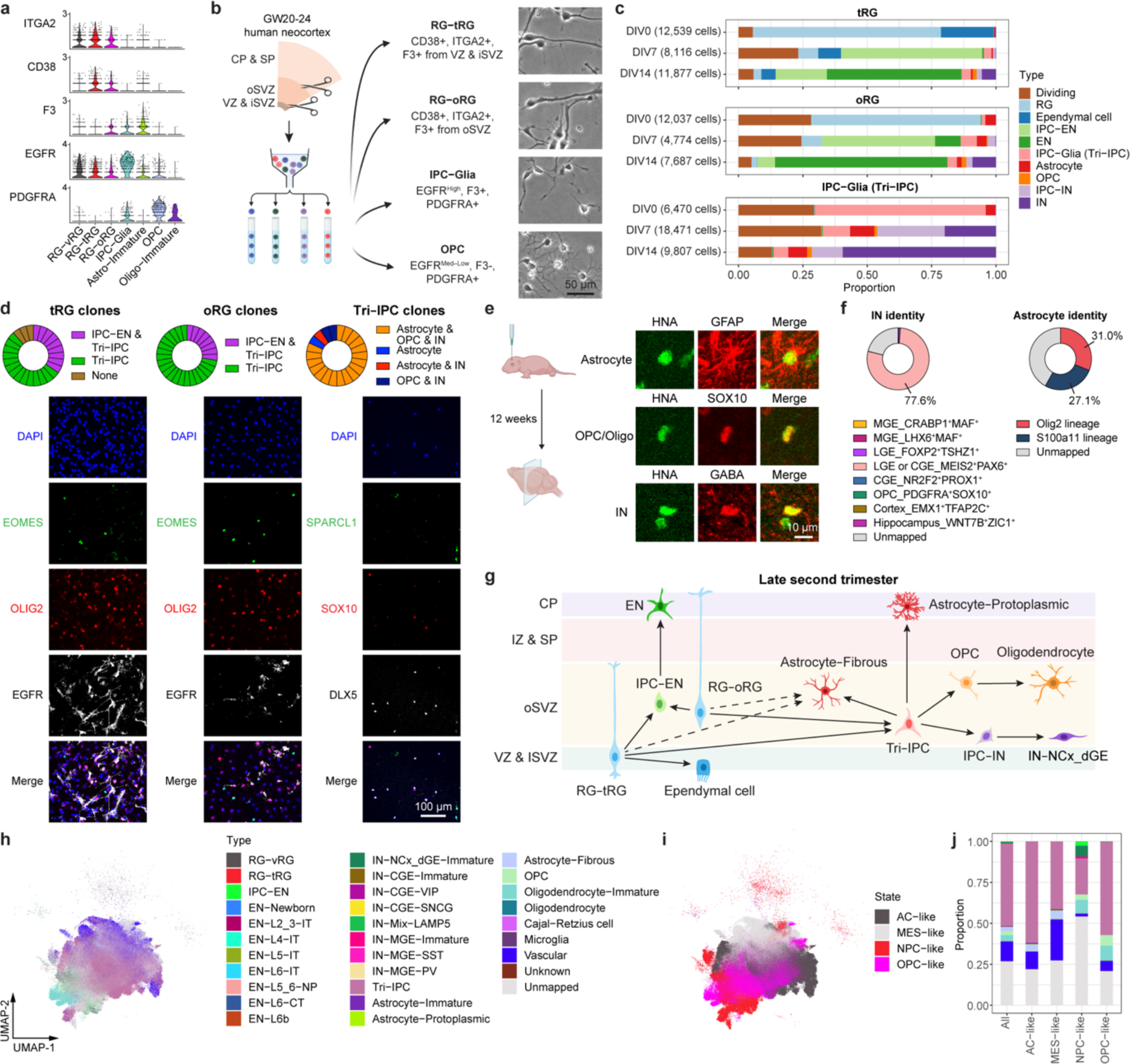
Multipotent progenitors during transition from neurogenesis to gliogenesis. **a**, Violin plots showing the expression patterns of surface proteins used for progenitor isolation. **b**, Left, schematic diagram showing the sorting strategy for isolation of progenitor subtypes. Right, phase-contrast images of progenitor subtypes after five days in culture. VZ & iSVZ, ventricular zone and inner subventricular zone; oSVZ, outer subventricular zone; CP & SP, cortical plate and subplate. **c**, Proportion of individual cell types across progenitor subtypes and differentiation stages during progenitor differentiation in vitro. **d**, Clonal analysis demonstrating multipotency of individual progenitor cells (n = 26, 29, 22 clones across three independent experiments). **e**, Immunostaining of progenies of Tri-IPCs 12 weeks after transplantation into mouse cortex, demonstrating presence of astrocytes (GFAP^+^), OPCs or oligodendrocytes (SOX10^+^), and IN (GABA^+^) (n = 2 injections). HNA, human nuclear antigen. **f**, SingleCellNet predicted identities of interneurons (INs) and astrocytes derived from Tri-IPCs. **g**, Graphical summary of cell lineage relationships in late second-trimester human neocortex. **h**, UMAP plots of malignant GBM cells color-coded by their main cellular states. **i**, UMAP plots of malignant GBM cells color-coded by SingleCellNet-predicted cell types. **j**, Proportion of predicted cell types across different cellular states in malignant GBM cells. The legend can be found in panel i.

Having validated our isolation strategy, we allowed cells to spontaneously differentiate without growth factor supplement for 14 days and performed scRNA-seq at DIV0, 7, and 14 to track their differentiation (Extended Data Fig. 15a, Supplementary Table 20). In the UMAP space, cells clustered according to the stage of differentiation, the seeding cell type, and their identity (Extended Data Fig. 15b–d). The scRNA-seq data revealed ten distinct cell types (Extended Data Fig. 15d,e, Methods). These cell types closely matched the in vivo populations observed in the snMultiome data (Extended Data Fig. 15f, Supplementary Table 21), confirming that the in vitro differentiation faithfully recapitulates the cell types found in vivo. Data from DIV0 reaffirmed the identities of the sorted cells (Fig. 4c, Extended Data Fig. 15g). On DIV7, three different types of descendants emerged in the IPC-Glia culture—astrocytes (9.4%), OPCs (1.1%), and a notable population of IN lineage cells, namely *DLX5*^+^*BEST3*^+^ IPC-INs (26.2%) and *DLX5*^+^*BEST3*^−^ INs (19.9%) (Fig. 4c, Extended Data Fig. 15e,g). Hence, we renamed IPC-Glia Tri-IPC to highlight their tripotency. The relatively low proportion of OPCs observed (1.1% on DIV7 and 1.8% on DIV14) could be attributed to the absence of specific growth factors required for their survival.

In contrast, both RG-tRGs and RG-oRGs differentiated into IPC-ENs at DIV7 and further into ENs by DIV14, indicating their continued production of ENs into the late second trimester (Fig. 4c, Extended Data Fig. 15g). Interestingly, Tri-IPCs emerged in both the RG-tRG and RG-oRG cultures by DIV7 (3.0% and 6.3%), along with a small proportion of IPC-INs (1.0% and 3.0%) but not INs (0.1% and 0.2%). By DIV14, astrocytes (0.7% and 1.8%), OPCs (1.5% and 1.8%), and INs (5.4% and 9.1%) were all present (Fig. 4c, Extended Data Fig. 15g). The delayed appearance of INs from RG cultures was consistent with our recent report that oRGs can produce INs^50^, but provided additional evidence that they do so indirectly through Tri-IPCs. Immunostaining further validated these results (Extended Data Fig. 14f–j).

The lineage tracing experiments described so far were conducted at the population level. To assess the lineage potential of glial progenitors at the single-cell level, we isolated individual RG-tRGs, RG-oRGs, and Tri-IPCs and cultured them for 14 days to produce clonal descendants. For both RG-tRGs and RG-oRGs, approximately 30% of all clones contained both IPC-ENs and Tri-IPCs, illustrating that individual RGs can generate both cell types (Fig. 4d, Supplementary Table 22). Moreover, about 80% Tri-IPC clones contained astrocytes, OPCs, and INs, confirming the tripotential nature of individual Tri-IPCs (Fig. 4d, Supplementary Table 22). Additionally, we transplanted isolated glial progenitors onto cultured human cortical slices ex vivo to provide a more physiologically relevant environment (Extended Data Fig. 16a). Consistent with our in vitro findings, RGs predominantly produced IPC-ENs within 8 days, whereas Tri-IPCs produced astrocytes, OPCs, and INs (Fig. 4f, Extended Data Fig. 16b–e). While Tri-IPCs maintained their tripotential nature in both in vitro and ex vivo conditions, we observed changes in the proportions of descendant cell types. Specifically, there was an increase in the production of OPCs in the ex vivo condition (Extended Data Fig. 16e), suggesting that cellular environment influences Tri-IPC fate decisions or descendant survival. To determine the lineage potential of Tri-IPCs in vivo, we performed xenograft experiments by transplanting Tri-IPCs into the cortex of early postnatal immunodeficient mice (Extended Data Fig. 16f). After 12 weeks of differentiation in vivo, the transplanted Tri-IPCs produced GFAP^+^ astrocytes, SOX10^+^ oligodendrocyte lineage cells, and GABA^+^ INs, predominantly in the deep layers of cortex, white matter, and subventricular zone (Fig. 4e, Extended Data Fig. 16g,h). Together, these results demonstrate that Tri-IPCs are tripotential neural progenitor cells.

To determine the specific subtype of INs produced by Tri-IPCs, we obtained scRNA-seq data from human ganglionic eminence as a reference^51^ and annotated interneuron subtypes based on marker genes reported in the literature^52^ (Extended Data Fig. 17a,b). We then trained a random-forest-based classifier using SingleCellNet^53^ based on this reference dataset, revealing that INs derived from Tri-IPCs closely resembled *MEIS2*^+^*PAX6*^+^ INs from dLGE and CGE (Fig. 4f). The finding is consistent with the fact that INs derived from Tri-IPCs map to IN-dLGE-Immatures in the snMultiome data (Extended Data Fig. 15f). These cells were also *SP8*^+^*SCGN*^+^ and were projected to develop into olfactory bulb interneurons and white matter interneurons^52^. This aligns with the presence of Tri-IPCs and IN-dLGE-Immatures in the white matter of both prenatal and postnatal human telencephalon observed in our MERFISH data (Extended Data Fig. 5a–f) and suggests that some of these IN-dLGE-Immatures may originate from Tri-IPCs. We, therefore, renamed IN-dLGE-Immatures as IN-NCx_dGE-Immatures to highlight their origin beyond dLGE. Similar results were obtained with a nearest-neighbor-based label transfer approach using Seurat (Fig. 4f, Extended Data Fig. 17c,d). Additionally, we aimed to categorize the types of astrocytes derived from Tri-IPCs. A recent study delineated two lineage origins of astrocytes in the mouse neocortex—an Olig2 lineage primarily producing gray matter or protoplasmic astrocytes and an S100a11 lineage primarily producing white matter or fibrous astrocytes^54^. We applied similar classification analysis using scRNA-seq data from the developing mouse neocortex^55^ and human snMultiome data from this study as references (Extended Data Fig. 17e,f,i,j). We found that Tri-IPC-derived astrocytes were mapped to both Olig2 and S100a11 lineages, indicating their potential to produce both protoplasmic and fibrous astrocytes (Fig. 4f, Extended Data Fig. 17g,h,k,l). Based on these results, we propose an updated model of the origin and lineage potential of human neural progenitors in the late second trimester (Fig. 4g).

### A majority of glioblastoma multiform cells resemble Tri-IPCs

Tri-IPCs produce neurons, oligodendrocyte lineage cells, and astrocytes, all considered important components of glioblastoma multiform (GBM)^56^. Previous studies also suggested the existence of glial progenitor cell-like populations in malignant GBM cells^10,57^. We aimed to leverage our comprehensive developmental atlas as a reference to map GBM cells to their closest counterparts in the developing human cortex. To this end, we trained a multi-class classifier using SingleCellNet based on our snMultiome data. We then used the trained model to assign cell types to malignant GBM cells from GBMap^58^. Our analysis revealed that more than half of malignant GBM cells transcriptionally resemble Tri-IPCs (Fig. 4h–j). Moreover, Tri-IPC was the most abundant mapped cell type across all four tumor cell states defined by Neftel et al. (Fig. 4j)^56^, and it was present in 87% of all GBM samples (Extended Data Fig. 17m). The second most abundant cell type is Vascular, which likely correspond to the glial-like wound response state (Fig. 4j)^59^. Other cell types substantially present in GBM include OPC, Oligodendrocytes-Immature, Astrocyte-Fibrous, and IN-NCx_dGE-Immature (Fig. 4j), all of which can be descendants from Tri-IPCs. These findings suggest that GBM cells hijack developmental processes, particularly the multipotency and proliferative capacity of Tri-IPCs, to achieve tumor heterogeneity and rapid growth.

### Cell type relevance to human cognition and brain disorders

Approximately 90% of variants identified in genome-wide association studies (GWASs) were found within non-protein-coding regions of the genome^60,61^. Leveraging the chromatin accessibility data we obtained from the developing human neocortex, we applied SCAVENGE^62^ to map GWAS variants to their relevant cellular context at single-cell resolution. Specifically, the algorithm quantifies the enrichment of GWAS variants within the open chromatin regions of a cell and overcomes the sparsity issue of single-cell profiles via network propagation. The enrichment strength was quantified by trait-relevance scores (TRSs) at the single-cell level and the proportion of significantly enriched cells at the cell-group level. Using this approach, we analyzed four cognitive traits and five neuropsychiatric disorders, revealing that they all had significant associations with specific cell types (Fig. 5a–c, Supplementary Table 23). Concerning cognitive traits, we found that fluid intelligence and processing speed were associated with IT neurons, aligning with previous results in the adult human brain (Fig. 5a,c) ^41^. In addition, we were surprised to observe an association between RGs and executive function and between microglia and working memory (Fig. 5a,c). The exact mechanisms underlying these associations remain to be elucidated. Regarding psychiatric disorders, all exhibited significant associations with various types of ENs (Fig. 5b,c). Bipolar disorder (BPD), schizophrenia (SCZ), and attention-deficit/hyperactivity disorder (ADHD), but not autism spectrum disorder (ASD) or major depressive disorder (MDD), were additionally linked to INs (Fig. 5b,c), highlighting differential disease association between the two major neuronal subclasses. Notably, some of the strongest associations were found between ASD and specific IT types (EN-IT-Immatures and EN-L6-ITs). As a control, we evaluated the association between neocortical cell types and Alzheimer’s disease, which is known to have a strong heritability component in microglia^63,64^. We not only observed the strongest enrichment of Alzheimer’s disease-associated variants in microglia but also identified significant enrichment in vascular cells and astrocytes (Extended Data Fig. 18a,b), consistent with their involvement in the disease^65,66^. It is important to note that our analysis was based on common variants and may not uncover contributions from other cell types due to the involvement of rare variants or environmental factors.

**Fig. 5.**
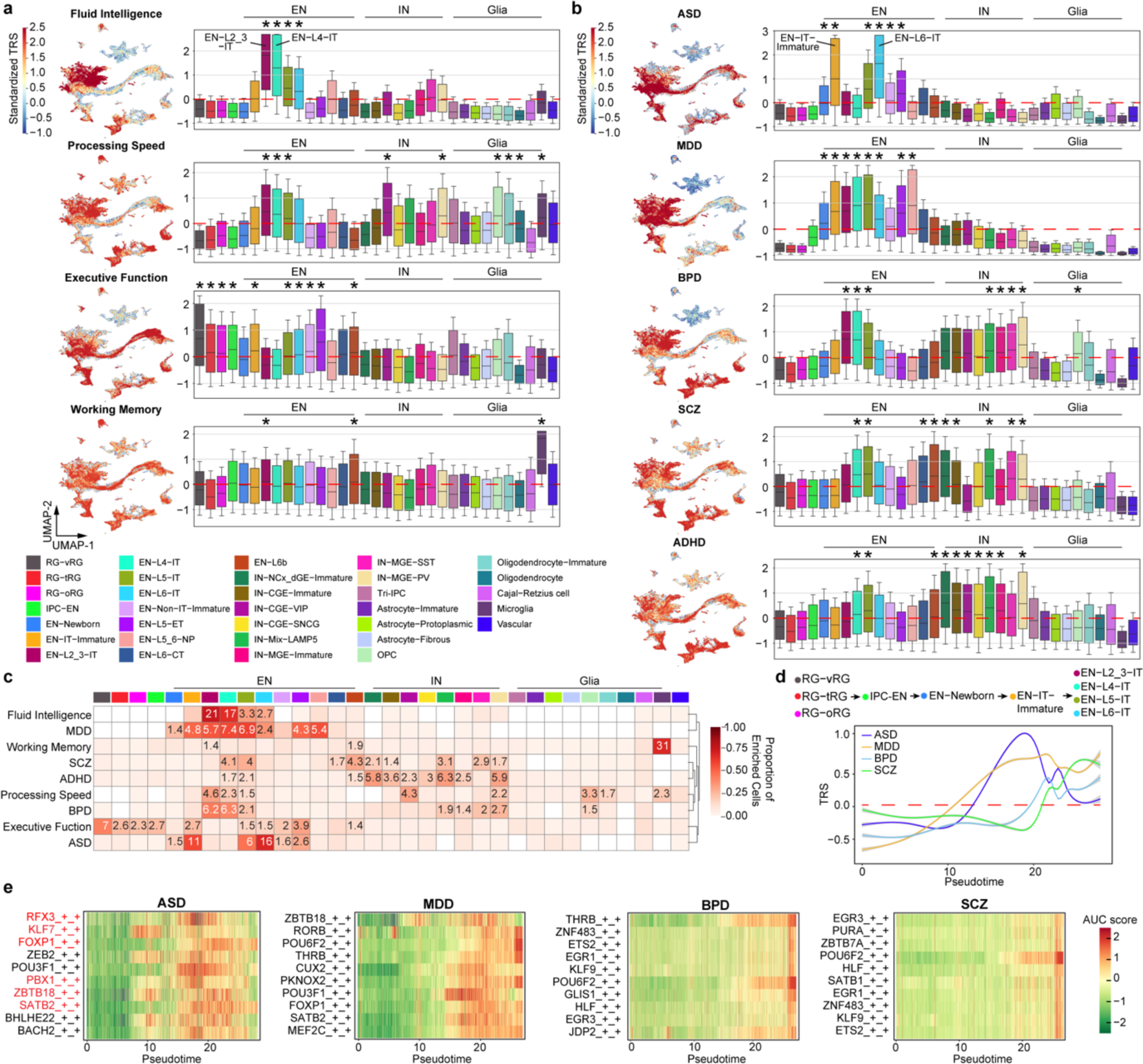
Cell type association with human cognition and brain disorders. **a**, Standardized per-cell SCAVENGE trait relevance score (TRS) for four cognitive functions. Boxplot center: median; hinges: the 25th and 75th percentiles; whiskers: standard error. **b**, Standardized per-cell SCAVENGE TRS for five brain disorders, including autism spectrum disorder (ASD), major depressive disorder (MDD), bipolar disorder (BPD), attention-deficit/hyperactivity disorder (ADHD), and schizophrenia (SCZ). Boxplot center: median; hinges: the 25th and 75th percentiles; whiskers: standard error. Two-sided hypergeometric test, *FDR < 0.01 & odds ratio > 1.4. **c**, Heatmap showing the proportion of the cells with enriched trait relevance across cell types. Tiles with significant TRS enrichment (two-sided hypergeometric test, *FDR < 0.01 & odds ratio > 1.4) are annotated by their odd ratios. **d**, Standardized SCAVENGE TRS of four brain disorders plotted along the intratelencephalic (IT) neuron lineage pseudotime. The best-fitted smoothed lines indicate the average TRS and the 95% confidence interval in each pseudo-time bin. **e**, Heatmaps of standardized gene-based AUC scores for top ten disease-relevant eRegulons ranked by Spearman’s ρ along the IT neuron lineage psuedotime. eRegulons with SFARI ASD-associated genes as core TFs are highlighted in red.

Besides cell types, we also compared trait associations among brain regions and age groups, revealing that differences between age groups were more pronounced than between regions (Extended Data Fig. 18c–f, Supplementary Table 24). For example, risk variants associated with neuropsychiatric disorders displayed distinct patterns of enrichment across age groups, with ASD risk enrichment peaking in the second trimester (Extended Data Fig. 18e–f). Given the predominant enrichment of these risk variants in ENs (Fig. 5b,c), we postulated that they target distinct stages of EN differentiation and maturation. To test this hypothesis, we selected EN lineage cells and examined the patterns of TRSs along their pseudotime (Fig. 5d). Indeed, ASD showed the earliest TRS peak, followed by MDD, BPD, and SCZ. This pattern is consistent with the earlier onset of ASD compared to other disorders and explains why previous heritability analyses of ASD in the adult brain found only a modest signal in ENs^41^. To pinpoint potential gene regulatory networks disrupted by disease risk variants during EN differentiation, we identified eRegulons whose activity positively correlated with the TRSs for each disorder (Fig. 5e, Supplementary Table 25). Among the core TFs of the top ten eRegulons correlated with ASD, six were recognized as ASD risk genes and listed in the SFARI gene database^67^. Together, our analysis not only pinpoints the most relevant cell types and developmental stages for cognitive traits and brain disorders but also elucidates potential disease mechanisms at cellular and molecular levels.

## Discussion

In this study, we extensively characterized the developing human neocortex at multiple stages, regions, and dimensions, including transcriptomic, epigenomic, spatial, and functional analyses. These data collectively establish an atlas of the developing human neocortex at single-cell resolution. The integration of multi-omic data provides insights into diverse aspects, including cellular composition, spatial organization, gene regulatory networks, lineage potential, and susceptibility to diseases during brain development. By combining spatial and snMultiome data, we further elucidate intricate cell-cell communication networks during development, emphasizing robust interactions between EN and IN subclasses mediated by specific signaling pathways.

V1 in humans, primates, and other binocular mammals exhibits a specialized cytoarchitecture characterized by an enlarged layer 4 that receives inputs from the thalamus^68^. Recent brain cell census studies in humans and non-human primates have identified a distinct population of EN-L4-ITs exclusively present in V1^21,69^. However, the mechanisms responsible for their emergence and the factors determining their identity have been unknown. Our results suggest that common and V1-specific EN-L4-ITs initially share a common developmental trajectory until the third trimester, after which they diverge. Common EN-L4-ITs follow a trajectory similar to that of EN-L5-IT, whereas V1-specific EN-L4-ITs partially share a trajectory with EN-L2_3-IT. Furthermore, we have identified TFs and eRegulons responsible for V1-specific EN-L4-ITs differentiation, including *SMAD3*, *GLIS3*, and *CUX2* at early stages, as well as *POU6F2*, *JDP2*, and *CUX1* at later stages. These results elucidate genetic programs governing sequential neuronal fate determination. They also offer crucial insights and serve as a benchmark for the future development of area-specific in vitro models of human cortical development.

Previous studies in rodents have demonstrated that following the peak neurogenesis of ENs, RGs within the dorsal telencephalon gradually transition to gliogenesis. Concurrently, they begin transitioning into a specific subtype of VZ/SVZ stem cells that produces olfactory bulb interneurons or becomes quiescent^70–73^. In humans and other non-human primates, however, a longstanding debate persists concerning two fundamental questions: firstly, whether cortical progenitors, particularly cortical RGs, can generate INs during embryonic development, and secondly, what subtype of neurons these INs eventually mature into^74–79^. Regarding the first question, most evidence supporting the “local production” hypothesis focuses primarily on identifying IN progenitors in the cortex, albeit not conclusively ruling out the possibility that these IN progenitors originate in the ventral telencephalon^80^. Recently, we and others have demonstrated that cortical RGs in the second trimester can produce LGE- and CGE-like INs that share a lineage with ENs^50,81^. However, whether these INs are generated directly from RGs or indirectly via IPCs remained uncertain. In this study, we observed that both oRGs and tRGs give rise to INs through Tri-IPCs, which are tripotential and capable of producing INs, oligodendrocyte lineage cells, and astrocytes locally in the neocortex. Based on the expression of EGFR and OLIG2, human Tri-IPCs likely correspond to “MIPCs” found in mice^49^ and proposed in humans^45^. The onset of Tri-IPC production occurs in the late second trimester (after GW18), potentially due to increased sonic hedgehog signaling during later stages of cortical development^82–85^. These findings provide an explanation for the limited production of INs observed in short-term cultures of human organotypic slices before GW18^76^. The involvement of Tri-IPCs in GBM is another interesting observation and helps explain how GBM cells maintain their stemness and achieve heterogeneity. Future research to understand Tri-IPC biology will offer insights into the cellular origins and potential vulnerabilities of GBM cells.

Concerning the identity of INs born in the cortex, our classification suggests that most Tri-IPC-derived INs are transcriptomically similar to *MEIS2*^+^*PAX6*^+^ INs presumed to originate from the dLGE^52^. Interestingly, these INs are also found in scRNA-seq data from the CGE^51^, consistent with the fact that *MEIS2*^+^ cells have been observed in the CGE^86^. Moreover, *MEIS2*^+^*PAX6*^+^ INs emerge in dorsally patterned human cerebral organoids, particularly at their later developmental stages^50^. Thus, instead of an IN type whose origin is confined to the LGE, we propose that *MEIS2*^+^*PAX6*^+^ INs represent the most dorsal type of IN generated within the germinal zone of the cortex and its neighboring ganglionic eminence. In mice, INs derived from MIPCs were reported to differentiate into olfactory bulb interneurons ^49^. However, our spatial transcriptomic data demonstrate the presence of *MEIS2*^+^*PAX6*^+^ INs in the white matter of both the prenatal and postnatal human brain, indicating their potential role as white matter interneurons. With recent reports of a shared origin between some cortical interneurons and ENs^87,88^, it remains to be determined whether INs derived from Tri-IPCs also differentiate into cortical interneurons.

Most genetic risk for ASD comes from common variants found in non-coding regions of the genome^89^. However, understanding the underlying cellular and molecular mechanisms has remained challenging due to a lack of comprehensive cell-type-resolved epigenomic data from the developing human brain. Our variant mapping at single-cell resolution reveals pronounced enrichment of ASD-linked common risk variants within chromatin-accessible regions specific to IT neurons in the second trimester, aligning with ASD as a neurodevelopmental disorder primarily originating at midgestation. The relevance of midgestational cortical development to ASD is further supported by data from gene expression analysis of both common and rare de novo ASD variants^90–92^. Moreover, our analyses indicate that disrupting intratelencephalic connectivity, particularly by impacting IT neurons in early development, may contribute to ASD pathophysiology. Notably, EN-IT-Immatures in the second trimester differentiate predominantly into EN-L2_3-ITs and EN-L4-ITs postnatally. Intriguingly, EN-L2_3-ITs and EN-L4-ITs are among the most affected cell types in post-mortem ASD brain^93^, highlighting how early-acting ASD risk variants cascade into postnatal deficits within IT neurons. Our analysis extends beyond ASD and reveals temporal- and cell-type-specific risk patterns associated with multiple brain disorders. For example, ASD exhibits the earliest risk, succeeded by MDD, and then followed by BPD and SCZ. Moreover, BPD, SCZ, and ADHD, but not ASD or MDD, were linked to inhibitory neurons, consistent with observations from the adult brain^41^. These findings underscore the significance of studying the typical trajectory of brain development in understanding the deviations leading to specific diseases.

## Supporting information

SI Guide

## Methods

### Brain tissue samples

Human brain tissue samples (Supplementary Table 1 and 5) were acquired from four sources.

Four de-identified first-trimester human tissue samples were collected from the Human Developmental Biology Resource (HDBR), staged using crown-rump length, dissected, and snap-frozen on dry ice.

Thirteen de-identified second-trimester human tissue samples were collected at the Zuckerberg San Francisco General Hospital (ZSFGH). Acquisition of second-trimester human tissue samples was approved by the UCSF Human Gamete, Embryo and Stem Cell Research Committee (study number 10-05113). All experiments were performed in accordance with protocol guidelines. Informed consent was obtained before sample collection and use for this study.

Two de-identified third-trimester and early postnatal tissue samples were obtained at the UCSF Pediatric Neuropathology Research Laboratory (PNRL) led by Dr. Eric Huang. These samples were acquired with patient consent in strict observance of the legal and institutional ethical regulations and in accordance with research protocols approved by the UCSF IRB committee. These samples were dissected and snap-frozen either on a cold plate placed on a slab of dry ice or in isopentane on dry ice.

Twenty-three de-identified third trimester, early postnatal, and adolescent tissue samples without known neurological disorders were obtained from the University of Maryland Brain and Tissue Bank through NIH NeuroBioBank.

Samples used for single-nucleus analysis were listed in Supplementary Table 1, and those for spatial transcriptomic analysis were listed in Supplementary Table 5.

### Nuclei isolation and generation of single-nucleus multiome (snMultiome) data

A detailed protocol can be found at ref^94^. All procedures were done on ice or at 4°C. Briefly, frozen tissue samples (20–50 mg) were homogenized using a pre-chilled 7 ml Dounce homogenizer containing 1 ml cold homogenization buffer (HB) (20 mM Tricine-KOH pH 7.8, 250 mM sucrose, 25 mM KCl, 5 mM MgCl_2_, 1 mM dithiothreitol, 0.5 mM Spermidine, 0.5 mM Spermine, 0.3% NP-40, 1× cOmplete protease inhibitor [Roche], and 0.6 U/mL RiboLock [Thermo Fisher]). The tissue samples were homogenized 10 times with the loose pestle and 15 times with the tight pestle. Nuclei were pelleted by spinning at 350 × g for 5 min, resuspended in 25% iodixanol solution, and loaded onto 30% and 40% iodixanol layers to make a gradient. The gradient was spun at 3,000 × g for 20 min. Clean nuclei were collected at the 30%–40% interface and diluted in wash buffer (10 mM Tris-HCl pH 7.4, 10 mM NaCl, 3 mM MgCl_2_, 1 mM dithiothreitol, 1% BSA, 0.1% Tween 20, and 0.6 U/mL RiboLock [Thermo Fisher]). Next, nuclei were pelleted by spinning at 500 × g for 5 min and resuspended in diluted nuclei buffer (10x Genomics). Nuclei were counted using a hemocytometer, diluted to 3220 nuclei/μL, and further processed following 10x Genomics Chromium Next GEM Single Cell Multiome ATAC + Gene Expression Reagent Kits user guide. We targeted 10,000 nuclei per sample per reaction. Libraries from individual samples were pooled and sequenced on the NovaSeq 6000 sequencing system, targeting 25,000 read pairs per nucleus for ATAC and 25,000 read pairs for RNA.

### snMultiome data pre-processing

The raw sequencing signals in the BCL format were demultiplexed into fastq format using the “mkfastq” function in the Cell Ranger ARC suite (v.2.0.0, 10x Genomics). Cell Ranger-ARC count pipeline was implemented for cell barcode calling, read alignment, and quality assessment using the human reference genome (GRCh38, GENCODE v32/Ensembl98) following the protocols described by 10x Genomics. The pipeline assessed the overall quality to retain all intact nuclei from the background and filtered out non-nucleus-associated reads. All gene expression libraries in this study showed a high fraction of reads in nuclei, indicating high RNA content in called nuclei and minimal levels of ambient RNA detected. The overall summary of data quality for each sample is listed in Supplementary Table 1. Next, we further assessed the data at the individual nuclei level and retained high-quality nuclei with the following criteria: (1) Gene expression count (nCount_RNA) is in the range of 1,000 to 25,000; (2) The number of detected genes (nFeature_RNA) is greater than 400; (3) The total ATAC fragment count in the peak regions (atac_peak_region_fragments) is in the range of 100 to 100,000; (4) The transcription start site (TSS) enrichment score for ATAC-seq is greater than 1; (5) The strength of nucleosome signal (the ratio of mononucleosome to nucleosome-free fragments) is below 2. To ensure only single nuclei were analyzed, we measured the doublet probability by Scrublet^95^ and excluded all potential doublets receiving a score greater than 0.3 for downstream analyses. In total, 243,535 nuclei that passed all QC criteria were included for further analysis.

### snMultiome data integration, dimensionality reduction, clustering, and cell type identification

For ATAC data of snMultiome analysis, open chromatin region peaks were called on individual samples using MACS2 (v2.2.7)^96^. Peaks from all samples were unified into genomic intervals, and the intervals falling in the ENCODE blacklisted regions were excluded^97^. Among all 398,512 processed ATAC peaks, the top 20% of consensus peaks (n = 82,505) across all nuclei were selected as variable features for downstream fragment counting and data integration. The peak-by-nuclei counts for each sample were integrated by reciprocal LSI projection functions using the R package Signac (v1.10.0)^98^. For RNA-seq data, normalization and data scaling were performed using SCTransform v2^99^ in Seurat v4^20^. The cell cycle difference between the G2M and S phase for each nucleus was scored and regressed out before data integration. The transformed gene-by-nuclei data matrices for all nuclei passing quality control were integrated by reciprocal PCA projections between different samples using Seurat v4 following the best practice described in Stuart et al.^98^ and Butler et al.^100^.

Weighted nearest neighbor analysis was done using Seurat v4 with 1–50 PCA components and 2–40 LSI components. The resulting nearest neighbor graph was used to perform UMAP embedding and clustering using the SLM algorithm^101^. Clusters with known markers expressed in the striatum (*ISL1* and *SIX3*) and diencephalon (*OTX2* and *GBX2*) were discarded. In addition, clusters with both transcripts present in neurites (*NRGN*) and oligodendrocyte processes (*MBP*), likely due to debris contamination, were discarded. These filtering steps resulted in 232,328 nuclei in the final dataset (Extended Data Fig. 1, Supplementary Table 2). Weighted nearest neighbor, dimension reduction, and clustering were re-calculated using the filtered data. Cell identities were determined based on the expression of known marker genes, as is shown in Extended Data Fig. 3 and Supplementary Table 3. The 5 identified classes were progenitor, neuron, glia, immune cell, and vascular cell. The 11 identified subclasses were radial glia, intermediate progenitor cell for excitatory neurons (IPC-EN), glutamatergic neuron, GABAergic neuron, intermediate progenitor cell for glia (IPC-Glia), astrocyte, oligodendrocyte precursor cell (OPC), oligodendrocyte, Cajal-Retzius cell, microglia, and vascular cell. The 33 identified cell types were ventricular radial glia (RG-vRG), truncated radial glia (RG-tRG), outer radial glia (RG-oRG), intermediate progenitor cell for excitatory neurons (IPC-EN), newborn excitatory neuron (EN-Newborn), immature intratelencephalic neuron (EN-IT-Immature), layer 2–3 intratelencephalic neuron (EN-L2_3-IT), layer 4 intratelencephalic neuron (EN-L4-IT), layer 5 intratelencephalic neuron (EN-L5-IT), layer 6 intratelencephalic neuron (EN-L6-IT), immature non-intratelencephalic neuron (EN-Non-IT-Immature), layer 5 extratelencephalic neuron (EN-L5-ET), layer 5–6 near-projecting neuron (EN-L5_6-NP), layer 6 corticothalamic neuron (EN-L6-CT), layer 6b neuron (EN-L6b), immature dorsal lateral ganglionic eminence inhibitory neuron (IN-dLGE-Immature), immature caudal ganglionic eminence inhibitory neuron (IN-CGE-Immature), VIP inhibitory neuron (IN-CGE-VIP), SNCG inhibitory neuron (IN-CGE-SNCG), LAMP5 inhibitory neuron (IN-Mix-LAMP5), immature medial ganglionic eminence inhibitory neuron (IN-MGE-Immature), SST inhibitory neuron (IN-MGE-SST), PVALB inhibitory neuron (IN-MGE-PV), intermediate progenitor cell for glia (IPC-Glia), immature astrocyte (Astrocyte-Immature), protoplasmic astrocyte (Astrocyte-Protoplasmic), fibrous astrocyte (Astrocyte-Fibrous), oligodendrocyte precursor cell (OPC), immature oligodendrocyte (Oligodendrocyte-Immature), oligodendrocyte (Oligodendrocyte), Cajal-Retzius cell, microglia (Microglia), and vascular cell (Vascular).

### Cell type proportion analysis

The investigation of variations in cell type proportions across different age groups and brain regions was conducted using a linear model approach implemented in the R packages speckle (v1.2.0)^102^ and limma (v3.58.1)^103^. To determine changes in cell type proportions over time, we logit-transformed the proportions within each sample and fitted a linear model (∼ log2_age + region) using limma. Moreover, to address the potential correlation among samples from the same individual, the duplicateCorrelation function in limma was applied. Once the model was fit, a moderated t-test with empirical Bayes shrinkage was used to test the statistical significance of the log2_age coefficient for each cell type. To determine cell type proportion differences between PFC and V1, a similar analysis was done, but only samples in the third trimester and older were used. Cell types with Benjamini–Hochberg adjusted P-values < 0.05 were determined significant (Supplementary Table 3).

### Transcription factor motif enrichment analysis

The per-cell regulatory activities of transcription factors (TFs) were quantified by chromVAR (v1.16.0)^104^. In brief, peaks were combined by removing any peaks overlapping with a peak with a greater signal, and only peaks with a width greater than 75bp were retained for motif enrichment analysis. We computed the per-cell enrichment of curated motifs from the JASPAR2020 database^105^. In total, 633 unique human transcriptional factors were assigned to their most representative motifs. The per-cell-type transcriptional activity of each TF was represented by averaging the per-cell chromVAR scores within the cell type, and the cell-type-specific TFs were chosen for further analysis and visualization (Supplementary Table 4).

### Spatial transcriptomic analysis using Multiplexed Error-Robust Fluorescence in situ Hybridization (MERFISH)

Spatial transcriptomic analysis using MERFISH was done using the Vizgen MERSCOPE platform. We designed a customized 300-gene panel composed of cell-type markers (Supplementary Table 5b) using online tools at https://portal.vizgen.com/. Fresh frozen human brain tissue samples were sectioned at a thickness of 10 µm using a cryostat and mounted onto MERSCOPE slides (Vizgen). Sections were fixed with 4% formaldehyde, washed three times with PBS, photo-bleached for 3 h, and stored in 70% ethanol for up to one week. Hybridizations with gene probes were performed at 37°C for 36–48 h. Next, sections were fixed using formaldehyde and embedded in a polyacrylamide gel. After gel embedding, tissue samples were cleared using a clearing mix solution supplemented with proteinase K for 1–7 days at 37°C until no visible tissue was evident in the gel. Next, sections were stained for DAPI and PolyT and fixed with formaldehyde before imaging. The imaging process was done on the MERSCOPE platform according to the manufacturer’s instructions. Cell segmentation was done using the Watershed algorithm based on Seed Stain (DAPI) and Watershed Stain (PolyT).

### MERFISH data integration, dimensionality reduction, clustering, cell type assignment, and niche analysis

Standard MERSCOPE output data were imported into Seurat v5^106^. We retained high-quality cells with the following criteria: (1) Cell volume is greater than 10 µm^3^; (2) Gene expression count (nCount_Vizgen) is in the range of 25 to 2,000; (3) The number of detected genes (nFeature_ Vizgen) is greater than 10. Normalization, data scaling, and variable feature detection were performed using SCTransform v2^99^. The transformed gene-by-cell data matrices for all cells passing quality control were integrated by reciprocal PCA projections between samples using 1–30 PCA components. After integration, nearest neighbor analysis was done with 1–30 PCA components. The resulting nearest neighbor graph was used to perform UMAP embedding and clustering using the Louvain algorithm^107^. Clusters with markers known to be mutually exclusive were deemed doublets and discarded. These filtering steps resulted in 404,030 cells in the final dataset (Supplementary Table 6). The identity of specific cell types was determined based on the expression of known marker genes, as is shown in Extended Data Fig. 4b. Niches were identified by k-means clustering cells based on the identities of their 50 nearest spatial neighbors.

### Frozen section staining to quantify the distribution of inhibitory neurons

GW23–24 human cortical samples were fixed in 4% paraformaldehyde in PBS at 4 °C overnight. The samples were cryoprotected in 15% and 30% sucrose in PBS and frozen in OCT. Samples were sectioned at a thickness of 16 µm, air-dried, and rehydrated in PBS. Antigen retrieval was carried out using citrate-based Antigen Unmasking Solution (Vector Laboratory) at 95 °C for 15 min. The slides were then washed in PBS and blocked in PBS-based blocking buffer containing 10% donkey serum, 0.2% gelatin, and 0.1% Triton X-100 at room temperature for 1 h. After blocking, slides were incubated with primary antibodies in the blocking buffer at 4°C overnight. The slides were washed in PBS and 0.1% Triton X-100 (PBST) three times and incubated with secondary antibodies in the blocking buffer at room temperature for 2 h. The slides were then washed in PBST three times as described above, counterstained with DAPI, and washed in PBS once more. Slides were mounted with coverslips using ProLong Gold (Invitrogen). Confocal tiled images were acquired with a Zeiss LSM900 microscope using a 20× air objective. Acquired images were processed using Imaris v9.7 (Oxford Instruments) and Fiji/ImageJ v1.54^108^. The following antibodies were used: NR2F2 (Abcam, ab211777, 1:250) and LHX6 (Santa Crux, sc-271433, 1:250).

### Neighborhood enrichment and intercellular communication modeling

To evaluate the spatial proximity of cell types in each sample, we obtained a neighborhood enrichment z-score using the nhood_enrichment function from Squidpy (v1.2.3)^109^. The graph neural network-based node-centric expression modeling (NCEM v0.1.4) method^31^ was used for intercellular communication modeling (Supplementary Table 7). A node-centric linear expression analysis was implemented to predict gene expression states from both cell type annotations and the surrounding neighborhood of each cell, where dependencies between sender and receiver cell types were constrained by the connectivity graph with a mean number of neighbors around 10 for each cell within each sample. One exception is that sample ARKFrozen-65-V1 was randomly downsampled to 60,000 cells to ensure it has a similar neighborhood size to other samples. Significant interactions were called if the magnitude of interactions (the Euclidean norm of coefficients in the node-centric linear expression interaction model) was above 0.5 and at least 25 differentially expressed genes (q value < 0.05 for specific sender-receiver interaction terms) were detected. For visualization purposes, only significant interactions were plotted in circular plots.

### Quantification of ligand-receptor (LR) communication using CellChat

We implemented CellChat (v1.6.1)^32^ to quantify the strength of interactions among cell types using default parameter settings (Supplementary Table 8). After normalization, the batch-corrected gene expression data from all 232,328 nuclei were taken as the CellChat input. We considered all curated ligand-receptor pairs from CellChatDB, where higher expression of ligands or receptors in each cell type was identified to compute the probability of cell-type-specific communication at the LR pair level (refer to the original publication for details). We filtered out the cell-cell communication if less than ten cells in the outgoing or incoming cell types expressing the ligand or receptor, respectively. The computed communication network was then summarized at a signaling pathway level and was aggregated into a weighted-directed graph by summarizing the communication probability. The calculated weights represent the total interaction strength between any two cell types. The statistically significant LR communications between the two groups were determined by one-sided permutation tests, where P value < 0.05 is considered significant.

### Organotypic slice culture and treatment with somatostatin receptor agonists

Primary cortical tissue from GW 16–24 was maintained in artificial cerebrospinal fluid (ACSF) containing 110 mM Choline chloride, 2.5 mM KCl, 7 mM MgCl_2_, 0.5 mM CaCl_2_, 1.3 mM NaH_2_PO_4_, 25 mM NaHCO_3_, 10 mM D-(+)-glucose, and 1 × Penicillin-Streptomycin. Before use, ACSF was bubbled with 95% O_2_/5% CO_2_. Cortical tissue was embedded in a 3.5% or 4% low-melting agarose gel. Embedded tissue was acutely sectioned at 300 μm thickness using a Leica VT1200 vibratome before being plated on Millicell inserts (Millipore, PICM03050) in 6-well tissue culture plates. Tissue slices were cultured at the air-liquid interface in media containing 32% HBSS, 60% Basal Medium Eagle, 5% FBS, 1% glucose, 1% N2 and 1 × Penicillin-Streptomycin-Glutamine. Slices were maintained for 12 h in culture at 37°C for recovery. After recovery, slices were grown in the presence of 1 μM Octreotide (SelleckChem, P1017), 4 μM (1R,1’S,3’R/1R,1’R,3’S)-L-054,264 (TOCRIS, 2444), or without any compound as a control. Slices were maintained for 72 hours in culture at 37°C, and the medium was changed every 24 hours.

### 10x fixed single-cell RNA Profiling of cultured slices treated with somatostatin receptor agonists

The cultured slices treated with somatostatin receptor agonists were fixed using the Chromium Next GEM Single Cell Fixed RNA Sample Preparation Kit (10x Genomics, 1000414) according to the manufacturer’s instructions. In brief, slices were finely minced on the pre-chilled glass petri dish, transferred into 1 mL fixation buffer, incubated at 4°C for 18 hours, and stored at - 80°C with 10% enhancer and 10% glycerol. After collecting all samples from six experimental batches, the stored samples were manually dissociated using Liberase TL (Sigma, 5401020001). Dissociated cells were counted using a hemocytometer and then proceeded to fixed single-cell RNA sequencing following the 10X Chromium Fixed RNA Profiling Reagent Kits (for Multiplexed Samples) user guide. Briefly, fixed single-cell suspensions were mixed with Human WTA Probes BC001–BC016, hybridized overnight (18 hours) at 42°C, washed individually, and pooled after the washing. Gene expression libraries were pooled and sequenced on the NovaSeq X sequencing platform, targeting 20,000 read pairs per cell.

The Cell Ranger multi pipeline was implemented for cell barcode calling, read alignment, and quality assessment using the human probe set reference (Chromium_Human_Transcriptome_Probe_Set_v1.0.1_GRCh38-2020-A) following the protocols described by 10x Genomics. The overall summary of data quality for each sample is listed in Supplementary Table 9. Next, we further assessed the data at the individual cell level and retained high-quality cells with the number of detected genes (nFeature_RNA) greater than 500. Doublets were removed using the R package scDblFinder (v1.18.0)^110^ with default settings. Normalization and data scaling were performed using SCTransform v2^99^. The transformed gene-by-cell data matrices for all cells passing quality control were integrated by reciprocal PCA projections between samples using 1–30 PCA components. After integration, nearest neighbor analysis was done with 1–30 PCA components. The resulting nearest neighbor graph was used to perform UMAP embedding and clustering using the Louvain algorithm^107^. Clusters with fewer UMI counts and markers known to be mutually exclusive were deemed low quality and discarded. These filtering steps resulted in 132,856 cells in the final dataset (Supplementary Table 10). The identity of specific cell types was determined based on the expression of known marker genes, as is shown in Extended Data Fig. 8b.

### Differential gene expression analysis to determine the effects of somatostatin receptor agonists

Pseudobulk differential gene expression analysis was done using the pseudoBulkDGE function from the R package scran (1.32.0). UMI counts were aggregated across cell types, individual patients, and treatment conditions. Pseudobulk samples with less than 10 cells were discarded. Next, we fitted the pseudobulked count data to a fixed-effect limma-voom model (∼ Patient_ID + Treatment). Once the model was fit, moderated *t*-tests were used to determine statistical significance through limma’s standard pipeline (Supplementary Table 11). The resulting moderated *t-*statistics of each gene were ranked and used as input for gene set enrichment analysis (GSEA) using the R package clusterProfiler^111^. GSEA was carried out against gene sets defined by the terms of biological processes in gene ontology (Supplementary Table 12). Only pathway sets with gene numbers between 10 and 500 were used for the analysis.

### Gene regulatory network analysis

We implemented the SCENIC+ (v0.1.dev448+g2c0bafd) workflow^33^ to build gene regulatory networks of the developing human neocortex based on the snMultiome data. As running the workflow on all nuclei is memory intensive, we subsampled 10,000 representative nuclei by geometric sketching^112^ to accelerate the analyses while preserving rare cell states and the overall data structure. First, MACS2 was used for consensus peak calling in each cell type^96^. Each peak was extended for 250bp in both directions from the summit. Next, weak peaks were removed, and the remaining peaks were summarized into a peak-by-nuclei matrix. Topic modeling was performed on the matrix by pycisTopic^113^ using default parameters, and the optimal number of topics (48) was determined based on log-likelihood metrics. Three different methods were used in parallel to identify candidate enhancer regions: (1) Regions of interest were selected by binarizing the topics using the Otsu method; (2) Regions of interest were selected by taking the top 3,000 regions per topic; (3) Regions of interest were selected by calling differentially accessible peaks on the imputed matrix using a Wilcoxon rank sum test (logFC > 0.5 and Benjamini–Hochberg adjusted P values < 0.05). Pycistarget and discrete element method (DEM) based motif enrichment analysis were then implemented to determine if the candidate enhancers were linked to a given TF^114^. Next, eRegulons, defined as TF-region-gene triplets consisting of a specific TF, all regions that are enriched for the TF-annotated motif, and all genes linked to these regions, were determined by a wrapper function provided by SCENIC+ using the default settings. We applied a standard eRegulon filtering procedure: (1) Only eRegulons with more than ten target genes and positive region-gene relationships were retained; (2) Only genes with top TF-to-gene importance scores were selected as the target genes for each eRegulon; 3) eRegulons with an extended annotation was only kept if no direct annotation is available. After filtering, 582 eRegulons were retained (Supplementary Table 13). For each retained eRegulon, specificity scores were calculated using the RSS algorithm based on region- or gene-based eRegulon enrichment scores (AUC scores)^115^ (Supplementary Table 14). eRegulons with top specificity scores in each cell type were selected for visualization. Finally, we extended our eRegulon enrichment analysis from the 10,000 sketched nuclei to all 232,328 nuclei by computing the gene-based AUC scores for all 582 eRegulons using the R package AUCell (v1.20.2)^36^ with default settings.

### Validation of the predicted eRegulons by SCENIC+

The predicted open chromatin regions (OCRs) regulated by the selected TFs in SCENIC+ were validated using ChIP-seq data described in Loupe et al.^34^. The data were downloaded from https://www.synapse.org/Synapse:syn51942384.1/datasets. We focused on available data for core TFs of eRegulons with > 10,000 ChIP-seq peaks, resulting in 24 datasets for further analysis. For each TF, the enrichment of eRegulon-targeted OCRs in the identified ChIP-seq peaks against the genomic background was computed as the odds ratio. The p-values were derived from the two-sided Fisher’s exact test, with corrections for multiple comparisons. The association of OCRs with their target genes was validated using long-range H3K4me3-mediated chromatin interactions captured by PLAC-seq^35^, where pairs with overlaps of both interaction bins were considered. The overrepresentation of OCR-to-gene interactions was tested using the two-sided Fisher’s exact test.

### Trajectory inference and trajectory-based differential expression analysis

Cells belonging to excitatory neuronal lineages, including radial glial cells, IPC-EN, and glutamatergic neurons, were selected from the whole dataset for trajectory inference using Slingshot (v2.6.0)^39^. A weighted nearest neighbor graph was re-calculated on the subset using 1– 50 PCA components and 2–40 LSI components. Dimension reduction was performed based on the calculated nearest neighbor graph, generating an 8-dimensional UMAP embedding. We identified 23 clusters in this UMAP space after removing one outlier cluster using mclust^116^. Next, we identified the global lineage structure with a cluster-based minimum spanning tree (MST). The cluster containing RG-vRG was set as the starting cluster, and those containing terminally differentiated cells were set as ending clusters (Extended Data Fig. 11a). Subsequently, we fitted nine simultaneous principal curves to describe each of the nine lineages, obtaining each cell’s weight based on its projection distance to the curve representing that lineage. Pseudotimes were inferred based on the principal curves, and shrinkage was performed for each branch for better convergence (Supplementary Table 16). Finally, the principal curves in the 8-dimensional UMAP space were projected to a 2-dimensional UMAP space for visualization.

### Identification of eRegulon modules

To model the activity of eRegulons along inferred trajectories, we fitted gene-based eRegulon AUC scores against pseudotimes by a generalized additive model (GAM) using tradeSeq (v1.12.0)^40^. As AUC scores can be seen as proportions data on (0,1), instead of the default negative binomial GAM, we fitted a beta GAM with six knots in tradeSeq. Fitted values from the tradeSeq models were extracted using the predictSmooth function, with 100 data points along each trajectory. The oRG&tRG trajectory was removed because we focused on excitatory neuronal lineages for eRegulon analysis. Based on fitted AUC values, six eRegulon modules were identified by k-means clustering (Supplementary Table 17a).

### Gene ontology enrichment analysis for eRegulon modules

The one-sided hypergeometric test implemented in clusterProfiler v4.0.5^111^ was used to identify overrepresented gene ontology (biological pathway) in each eRegulon module (Supplementary Table 17b). Genes present in at least 8% of all eRegulons in a module were regarded as the core target genes of that module. Module-specific core target gene sets were used as input gene sets. The union of target genes of any eRegulon was used as the background.

### Differential gene expression analysis between common and V1-specific EN-L4-IT

To identify genes differentially expressed between common and V1-specific EN-L4-IT, we first selected all EN-L4-IT nuclei and determined their subtype identity (common or V1-specific) based on markers and tissue of origin (Extended Data Fig. 12a,b). We then aggregated counts across samples and subtypes to generate pseudobulk samples. Differential gene expression analysis was done by fitting the pseudobulked count data to a generalized linear mixed model (∼ subtype + log2_age + [1 | dataset]) using the R package glmmSeq (v0.5.5)^117^. Size factors and dispersion were estimated using the R package edgeR (v3.42.4)^118^. Once the model was fit, likelihood ratio tests were used to determine statistical significance using (∼log2_age + [1 | dataset]) as the reduced model. Genes with Benjamini–Hochberg adjusted P-values < 0.05 were determined significant (Supplementary Table 18).

### Identification of key eRegulons that regulate neuronal lineage divergence

Based on the principal curves, five bifurcation points (BPs) were identified along neuronal differentiation. To identify genes that are differentiating around a BP of the trajectory, we performed an earlyDETest using tradeSeq. Specifically, we first separated the pseudotimes into five consecutive segments (Extended Data Fig. 11g). We then compared the expression patterns of gene-based eRegulon AUCs along pseudotime between lineages by contrasting 12 equally spaced pseudotimes within segments that enclose the BP (Supplementary Table 19). We included segments 2–3 for BP1, segments 3–4 for BP2, and segments 4–5 for BP3, BP4, and BP5.

### Isolation and in vitro culture of glial progenitors from late second-trimester human cortex

Glial progenitor cells were isolated from GW20–24 human dorsal cortical tissue samples. The ventricular zone/inner subventricular zone (VZ/iSVZ) and outer subventricular zone (oSVZ) were dissected and dissociated using the Papain Dissociation System (Worthington Biochemical). Dissociated cells were layered onto undiluted papain inhibitor solution (Worthington Biochemical) and spun down at 70 × g for 6 min to eliminate debris. The cell pellet was resuspended in 10 mL complete culture medium (DMEM/F12, 2 mM GlutaMAX, 2% B27 without vitamin A, 1% N2, and 1 × Penicillin-Streptomycin) and incubated at 37°C for 3 h for surface antigen recovery. From this point on, cells were handled on ice or at 4°C. Cells were washed once with staining buffer (Hank’s Balanced Salt Solution [HBSS] without Ca^2+^ and Mg^2+^, 10 mM HEPES pH 7.4, 1% BSA, 1 mM EDTA, 2% B27 without vitamin A, 1% N2, and 1 × Penicillin-Streptomycin), spun down at 300 × g for 5 min, and resuspended in staining buffer to a density of 1 × 10^8^ cells/mL. Cells were blocked by FcR Blocking Reagent (Miltenyi Biotech, 1:20) for 10 min, followed by antibody incubation for 30 min. Antibodies used for fluorescence-activated cell sorting (FACS) include FITC anti-EGFR (Abcam, ab11400), PE anti-F3 (Biolegend, 365204), PerCP-Cy5.5 anti-CD38 (BD Biosciences, 551400), Alexa Fluor 647 anti-PDGFRA (BD Biosciences, 562798), and PE-Cy7 anti-ITGA2 (Biolegend, 359314). All antibodies were used at 1:20 dilution. After incubation, cells were washed twice in staining buffer, resuspending in staining buffer containing Sytox Blue (Invitrogen), and sorted using BD FACSAria II sorters. Cells were sorted into collection buffer (HBSS without Ca^2+^ and Mg^2+^, 10 mM HEPES pH 7.4, 5% BSA, 2% B27 without vitamin A, 1% N2, and 1 × Penicillin-Streptomycin). After sorting, cells were spun down at 300 × g for 5 min, resuspended in complete culture medium, and plated onto glass coverslips pre-coated with poly-D-lysine and laminin at a density of 2.5 × 10^4^ cells/cm^2^. Cells were cultured in a humidified incubator with 5% CO_2_ and 8% O_2_. Half of the medium was changed with fresh medium every 3–4 days until harvest at the indicated time.

### Immunostaining of cultured cells and confocal imaging

On DIV0 and DIV14, glial progenitors or their progenies were fixed with 4% formaldehyde/4% sucrose in PBS and permeabilized/blocked with PBS-based blocking buffer containing 10% donkey serum, 0.2% gelatin, and 0.1% Triton X-100 at room temperature for 1 h. Samples were then incubated with primary antibodies diluted in the blocking buffer at 4 °C overnight. The next day, samples were washed in PBS three times and incubated with secondary antibodies in the blocking buffer at room temperature for 1 h. Samples were then washed twice in PBS, counterstained with DAPI, and washed in PBS again. Z-stack images were acquired with a Leica TCS SP8 using a 25× water immersion objective. Acquired images were processed using Imaris v9.7 (Oxford Instruments) and Fiji/ImageJ v1.54 ^108^. The following antibodies were used: TFAP2C (R&D systems, AF5059, 1:50), CRYAB (Abcam, ab13496, 1:200), OLIG2 (Abcam, ab109186, 1:150), EGFR (Abcam, ab231, 1:200), SPARCL1 (R&D systems, AF2728, 1:50), DLX5 (Sigma, HPA005670, 1:100), and NeuN (EMD Millipore, ABN90, 1:250).

### Single-cell RNA-seq analysis of glial progenitor differentiation

Glial progenitors were either immediately subjected to single-cell RNA-seq or cultured in vitro for 7 and 14 days before single-cell RNA-seq. In the latter cases, cells were released using the Papain Dissociation System (Worthington Biochemical) without DNase for 20 min. Released cells were washed twice in HBSS without Ca^2+^ and Mg^2+^ supplemented with 0.04% BSA, spun down at 250 × g for 5 min, and resuspended in HBSS without Ca^2+^ and Mg^2+^ supplemented with 0.04% BSA. Cells were counted using a hemocytometer, diluted to ∼1000 nuclei/μL, and further processed following the 10x Genomics Chromium Single Cell 3’ Reagent Kits User Guide (v3.1 Chemistry). We targeted 10,000 cells per sample per reaction. Libraries from individual samples were pooled and sequenced on the NovaSeq 6000 sequencing system, targeting 22,500 read pairs per cell.

The raw sequencing signals in the BCL format were demultiplexed into fastq format using the “mkfastq” function in the Cell Ranger suite (v.7.1.0, 10x Genomics). Cell Ranger count pipeline was implemented for cell barcode calling, read alignment, and quality assessment using the human reference genome (GRCh38, GENCODE v32/Ensembl98) following the protocols described by 10x Genomics. The pipeline assessed the overall quality to retain all intact cells from the background and filtered out non-cell associated reads. All gene expression libraries in this study showed a high fraction of reads in cells, indicating high RNA content in called cells and minimal levels of ambient RNA detected. The overall summary of data quality for each sample is listed in Supplementary Table 15. Next, we further assessed the data at the individual cell level and retained high-quality cells with the following criteria: (1) The number of detected genes (nFeature_RNA) is greater than 1000 and less than 10,000; (2) less than 10% of all reads mapped to mitochondrial genes. Raw counts were log-normalized with a size factor of 10,000. The first 30 principal components were used to construct the nearest neighbor graph, and Louvain clustering was used to identify clusters. Clusters with significantly fewer UMI counts, likely consisting of low-quality, dying cells, were also excluded for further analysis. The identity of specific cell types was determined based on the expression of known marker genes (Extended Data Fig. 15e, Supplementary Table 21). The 10 identified cell types were dividing cell (Dividing), radial glia (RG), ependymal cell (Ependymal cell), intermediate progenitor cell for excitatory neurons (IPC-EN), tripotential intermediate progenitor cell (Tri-IPC), astrocyte (Astrocyte), oligodendrocyte precursor cell (OPC), intermediate progenitor cell for inhibitory neurons (IPC-IN), and inhibitory neurons (IN).

### Classification of glial progenitor-derived cells by SingleCellNet

To determine the similarity between glial progenitor-derived cells and our atlas data, we applied SingleCellNet (v0.1.0), a random-forest-based cell type classification method^53^. Specifically, we randomly selected 700 cells from each cell type as the training set. We found the top 60 most differentially expressed genes per cell type, and then ranked the top 150 gene pairs per cell type from those genes. The preprocessed training data were then transformed according to the selected gene pairs and were used to build a multi-class classifier of 1000 trees. Additionally, we created 400 randomized cell expression profiles to train up an “unknown” category in the classifier. After the classifier was built, we selected 165 cells from each cell type from the held-out data, along with another 165 randomized cells, and assessed the performance of the classifier on the held-out data using Precision-Recall curves, obtaining an average AUPRC of 0.827. To classify Tri-IPC-derived inhibitory neurons, we transformed the query data with top pairs selected from the optimized training data and classified it with the trained classifier. Here, we chose a classification score threshold of 0.2, and cells with scores below this threshold were assigned as unmapped.

### Clonal analysis of glial progenitors

For clonal analysis, samples for FACS were processed as above with the following changes: individual tRG, oRG, or Tri-IPC was sorted using a BigFoot Spectral Cell Sorter (Thermo Fisher) via single-cell precision mode into a single well of 96-well glass-bottom plates pre-coated with polyethylenimine and laminin containing 100 μL complete culture medium. For tRG and oRG, the complete culture medium was supplemented with 10 ng/mL FGF2 to promote initial cell survival and proliferation. The culture medium was changed weekly for a total of two weeks. After two weeks, cells were fixed and stained in the same way as mentioned above. The following antibodies were used: EOMES (Abcam, ab23345, 1:200), OLIG2 (EMD Millipore, MABN50, 1:200), EGFR (Abcam, ab231, 1:200), SPARCL1 (R&D systems, AF2728, 1:50), SOX10 (Santa Cruz, sc-365692, 1:50) and DLX5 (Sigma, HPA005670, 1:100).

### Glial progenitor slice transplantation assay

Glial progenitors were isolated from GW 20–24 primary cortical tissue by FACS, as mentioned above. About 200,000 Cells were spun down at 300 × g for 5 min and resuspended in 0.5 mL complete culture medium containing 1 × 10^7^ PFU CMV-GFP adenoviruses (Vector Biolabs). Next, cells were incubated in a low attachment plate for 1 hour under the normal culture condition. After infection, cells were washed twice with complete culture medium containing 0.3% BSA and resuspended in slice culture medium. About 25,000 cells were transplanted onto the oSVZ of freshly prepared slices through a pipette. Slices were maintained for 8 days in culture at 37°C, and the medium was changed every other day.

After 8 days in culture, slices were fixed with 4% formaldehyde in PBS at room temperature for 1 h, followed by permeabilization and blocking with PBS-based blocking buffer containing 10% donkey serum, 0.2% gelatin, and 1% Triton X-100 at room temperature for 1 h. Samples were then incubated with primary antibodies diluted in the blocking buffer at 4 °C for 48 h. Two days later, samples were washed in PBS plus 0.1% Triton X-100 four times and incubated with secondary antibodies in the blocking buffer at 4 °C for 24 h. After secondary antibody incubation, samples were washed twice in PBS plus 0.1% Triton X-100, counterstained with DAPI, and washed in PBS again. Z-stack images were acquired with a Leica TCS SP8 using a 25× water immersion objective. Acquired images were processed using Imaris v9.7 (Oxford Instruments) and Fiji/ImageJ v1.54 ^108^. The following antibodies were used: GFP (Aveslabs, GFP-1020, 1:1,000), EOMES (Abcam, ab23345, 1:200), NeuN (EMD Millipore, ABN90, 1:250), OLIG2 (EMD Millipore, MABN50, 1:200), EGFR (Abcam, ab32077, 1:200), DLX5 (Sigma, HPA005670, 1:100), and SPARCL1 (R&D systems, AF2728, 1:50).

### Glial progenitor xenograft assay

FACS-sorted Tri-IPCs (60,000 cells) were spun down and resuspended in Leibovitz’s L-15 medium with DNAse I (180 μg/ml). Immediately before transplantation, cells were further concentrated by centrifugation (4 min, 800 × g) and resuspended in 2 μL Leibovitz’s L-15 with DNAse I. Cell suspension was loaded into beveled glass micropipettes (about 70–90 μm in diameter, Wiretrol 5 μl, Drummond Scientific Company) prefilled with mineral oil and mounted on a microinjector. Recipient mice (NSG, JAX 005557, postnatal day 5) were anesthetized by hypothermia (about 4 minutes) and positioned in a clay head mold to stabilize the skull^119^. Micropipettes were positioned vertically in a stereotactic injection apparatus. Injections were performed in both the left and right hemispheres perpendicular to the skin surface. Eye coordinates were x: 1.5, y: 3.6. Fifty nanoliters of cell suspension were released at z: 0.2, 0.4, 0.8, and 1 from the surface of the skin. Mice were returned to their litters after injection.

### Immunostaining of xenografted human cells

Twelve weeks after injection, the recipient mice were perfused with 4% paraformaldehyde (PFA) and post-fixed in 4% PFA at 4 °C overnight. The samples were cryoprotected in 15% and 30% sucrose in PBS and frozen in OCT. Samples were sectioned at a thickness of 16 µm, air-dried, and rehydrated in PBS. Immunostaining was done in the same way as described above for human brain sections. Confocal images were acquired with a Leica TCS SP8 using a 20× oil immersion objective. Acquired images were processed using Fiji/ImageJ v1.54^108^. The following antibodies were used: Human Nuclear Antigen (Abcam, ab191181, 1:200), GABA (Sigma, A2052, 1:250), GFAP (Invitrogen, 13-0300, 1:300), and SOX10 (R&D Systems, AF2864, 1:50).

### Classification of Tri-IPC-derived inhibitory neurons

Human ganglionic eminence single-cell RNA-seq data from Shi et al.^51^ were downloaded from GEO (GSE135827) and used as the reference. We integrated all samples using the RPCA methods, subset the data to focus on cells from the ganglionic eminence, re-clustered the cells, and annotated interneuron subtypes based on marker genes reported in the literature^52^ (Extended Data Fig. 17a,b).

To determine the identity of Tri-IPC-derived inhibitory neurons based on the reference dataset, we applied SingleCellNet in a similar way as mentioned above with the following parameter modifications. We randomly selected 400 cells from each cell type as the training set. We found the top 200 most differentially expressed genes per cell type, and then ranked the top 200 gene pairs per cell type from those genes. The preprocessed training data were then transformed according to the selected gene pairs and were used to build a multi-class classifier of 1000 trees. Additionally, we created 400 randomized cell expression profiles to train up an “unknown” category in the classifier. After the classifier was built, we selected 100 cells from each cell type from the held-out data, along with another 100 randomized cells, and assessed the performance of the classifier on the held-out data using Precision-Recall curves, obtaining an average AUPRC of 0.901. To classify Tri-IPC-derived inhibitory neurons, we transformed the query data with top pairs selected from the optimized training data and classified it with the trained classifier. Here, we chose a classification score threshold of 0.35, and cells with scores below this threshold were assigned as unmapped.

As an alternative classification method to determine the identity of Tri-IPC-derived inhibitory neurons, we performed mutual nearest neighbors-based label transfer using the MapQuery() function in Seurat v4. The first 30 principal components were used to identify transfer anchors. Cell type labels from Shi et al. were transferred to Tri-IPC-derived inhibitory neurons when confidence was high (prediction score > 0.5). Cells with prediction scores equal to or lower than 0.5 were labeled as unmapped.

### Classification of Tri-IPC-derived astrocytes

Mouse single-cell RNA-seq data from Di Bella et al.^55^ were downloaded from the Single Cell Portal (SCP1290) and used as the reference. We subset the data and focused on astrocytes and cycling glial cells (defined by the original authors). These cells were re-clustered and annotated as Olig2 or S100a11 lineages based on marker genes reported in the literature^54^ (Extended Data Fig. 17e,f). We used Tri-IPC-derived astrocytes as the query data and applied SingleCellNet in the same way as for Tri-IPC-derived inhibitory neurons. We also applied Seurat label transfer in the same way, except that 20 principal components were used to identify transfer anchors.

We also used astrocytes at the infancy stage from our snMultiome data, when we were able to distinguish the two astrocyte lineages, as the reference. We selected the astrocytes at infancy from the whole dataset and redid nearest neighbor analysis with 1–50 PCA components (already computed after SCTransform and RPCA integration). These cells were re-clustered on the basis of the resulting nearest neighbor graph and annotated on the basis of marker genes reported in the literature^54^ (Extended Data Fig. 17,j). We used Tri-IPC-derived astrocytes as the query data, which was re-processed in the same way as snMultiome data, including SCTransform v2 modeling and cell cycle regression. SingleCellNet was applied in the same way as above. For Seurat label transfer, the first 50 principal components were used to identify transfer anchors.

### Classification of human glioblastoma multiforme cells

We obtained single-cell and single-nucleus RNA-seq data of human glioblastoma multiforme cells from the extended GBmap^58^, downloaded from cellxgene (https://datasets.cellxgene.cziscience.com/ead761be-309f-4b79-8208-41da14ca305f.h5ad). Using the snMultiome atlas data as a reference, we applied SingleCellNet to identify the corresponding cell types of malignant cells in the GBmap. SingleCellNet was executed using the same parameters that were previously applied for the classification of glial progenitor-derived cells. Our analysis yielded an average AUPRC of 0.832. For classification, we set a score threshold of 0.15; cells with scores below this threshold were designated as unmapped.

### Building single-cell risk map for cognitive traits and brain disorders by SCAVENGE

We implemented SCAVENGE (v1.0.2)^62^ to integrate the snATAC-seq part of the snMultiome data with GWAS data of four cognitive traits (fluid intelligence, processing speed, executive function, and working memory) and five neuropsychiatric disorders (autism spectrum disorder [ASD], major depressive disorder [MDD], bipolar disorder [BPD], attention-deficit/hyperactivity disorder [ADHD], and schizophrenia [SCZ]). Analysis of Alzheimer’s disorder was included as a positive control. For each trait or condition, we performed multi-SNP-based conditional and joint association analysis on all GWAS SNPs with default settings. A stepwise model selection procedure was implemented to select independently associated SNPs and compute the fine-mapped posterior probability (PP). The PP was imported for our subsequent gchromVAR analysis^120^, where we built a cell-by-peak count matrix using peak called from integrated snATAC-seq data. A gchromVAR score indicating potential GWAS signal enrichment over a set of background peaks was calculated for each cell after correcting GC bias. To minimize the batch effects, we used the batch-aligned LSI matrix for the nearest neighbor graph construction and subsequent network propagation. A trait relevant score (TRS) representing the potential GWAS risk association was assigned to each cell to construct the single-cell risk map for cognitive traits or neurological disorders. To determine the significant trait-cell association, we considered cells receiving the top 0.1% TRS score traits-relevant and permuted the network propagation 1000 times for statistical significance. Cells with a P value less than 0.05 were defined as trait-associated. To determine the trait relevance per cell type, we calculated the odds ratio of cells associated with each trait in each cell type over the background and determined statistical significance by two-sided hypergeometric test followed by Benjamini-Hochberg correction. Cell types with FDR < 0.05 and odds ratio > 1.4 were deemed significantly enriched for trait-associated variants. A similar analysis was done for regions and age groups. Finally, the TRS scores were standardized by z transformation for comparison and visualization (Supplementary Table 23, Supplementary Table 24). The GWAS data used in this study can be downloaded from the following links: fluid intelligence (phenocode 20016), processing speed (phenocode 20023), executive function (phenocode 399), and working memory (phenocode 4282): https://pan.ukbb.broadinstitute.org/downloads/; ASD: https://figshare.com/articles/dataset/asd2019/14671989; MDD: https://datashare.ed.ac.uk/handle/10283/3203; BPD: https://figshare.com/articles/dataset/bip2021_noUKBB/; ADHD: https://figshare.com/articles/dataset/adhd2022/22564390; SCZ: https://figshare.com/articles/dataset/cdg2018-bip-scz/14672019; ALZ: https://ctg.cncr.nl/software/summary_statistics.

## Acknowledgements

We thank the NIH NeuroBioBank and the University of Maryland School of Medicine Brain and Tissue Bank for providing post-mortem brain tissue samples. We thank the Human Developmental Biology Resource for providing first-trimester brain tissue samples. We thank members of the A. R. Kriegstein laboratory and T. Nowakowski laboratory for helpful discussions. This study was supported by Simons Foundation Autism Research Initiative grant 697827 to J.L. and A.R.K., National Institute of Mental Health (NIMH) grant U01MH114825 to A.R.K. and E.J.H., National Institute of Neurological Disorders and Stroke (NINDS) grant R35NS097305 to A.R.K., NINDS grant P01NS083513 to A.A.-B., E.J.H., and A.R.K., NINDS grant R01NS123912 to X.D., and NIMH grant K99MH131832 to L.W.. J.A.M. was supported by funding from the Government of Catalonia (FI-SDUR 20) and from The Company of Biologists – Development (Travelling Fellowship).

## Author contributions

Conceptualization: L.W., C.W., J.L., A.R.K.; data curation: L.W., C.W.; formal analysis: L.W., C.W., J.A.M.; funding acquisition: L.W., E.J.H., A.A.-B., X.D., J.L., A.R.K.; investigation: L.W., S.C., G.Z., A.C.-S., S.Z., T.M., S.W., M.S., L.G.O., Q.B.; methodology: L.W., A.C.-S., S.Z., X.G.; resources: S.W., M.F.P., E.J.H., A.R.K.; software: L.W., C.W., J.A.M., J.J.A.; supervision: A.A.-B., X.D., J.L., A.R.K.; visualization: L.W., C.W., J.A.M.; writing – original draft: L.W., C.W., J.A.M.; writing – review & editing: all authors.

## Data availability

All raw and aligned snMultiome sequencing data were deposited to NeMO archive (https://assets.nemoarchive.org/dat-oiif74w). MERFISH data were deposited to Brain Image Library (https://doi.org/10.35077/g.1156). Processed data are available at an interactive portal (https://cell.ucsf.edu/snMultiome), at CELLxGENE (https://cellxgene.cziscience.com/collections/ad2149fc-19c5-41de-8cfe-44710fbada73), and at DRYAD (https://doi.org/10.5061/dryad.2280gb612). For inquiries regarding the acquisition of these data, please reach out to the corresponding authors.

## Code availability

The code used for data analysis in this manuscript is available at GitHub (https://github.com/complexdisease/Human_Cortex_Dev_Multiome).

---

**Extended Data Fig. 1.**
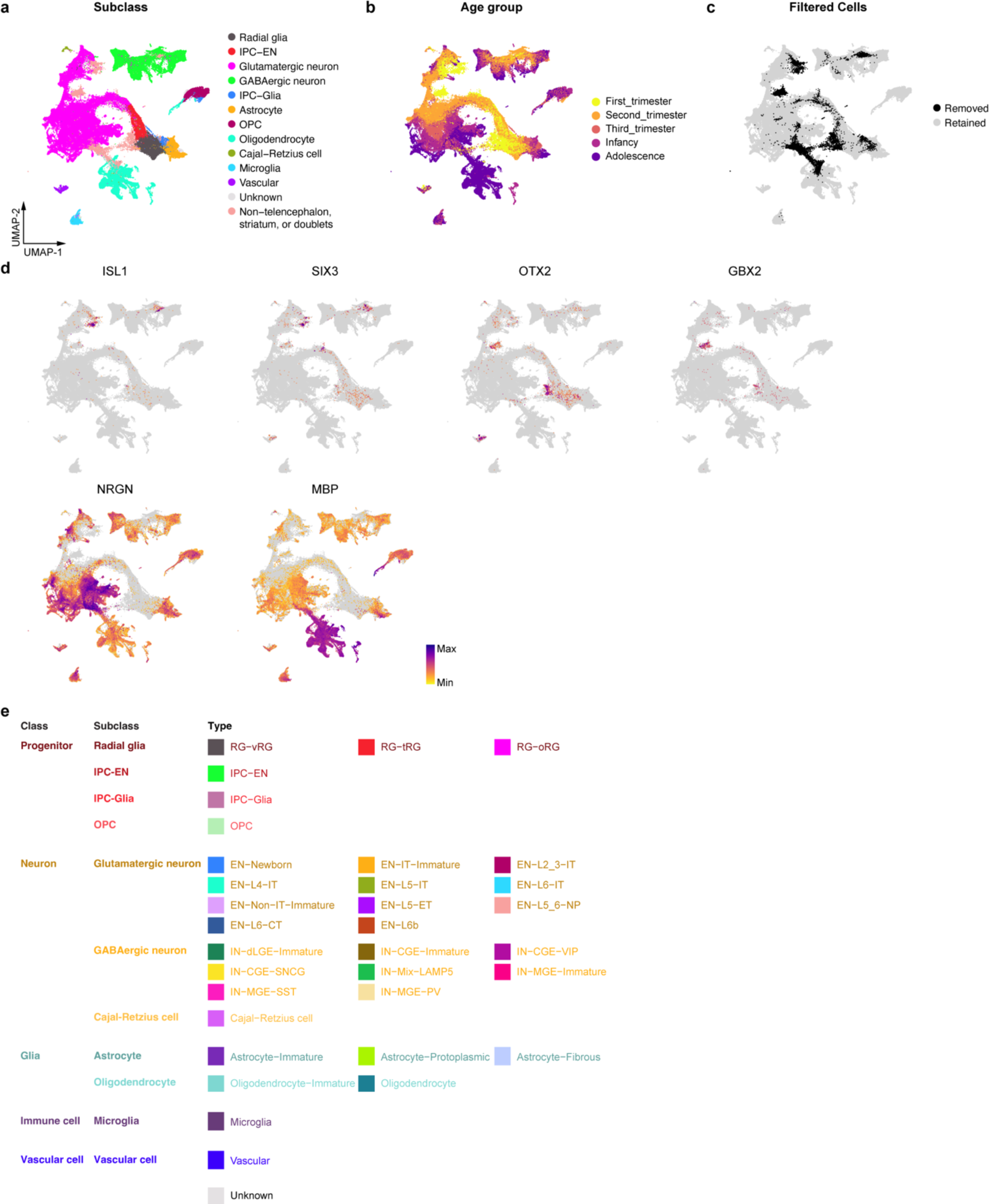
Filtering of the snMultiome data. **a**, UMAP plots showing the distribution of cell subclasses in the single-nucleus multiome data prior to data filtering. **b**, UMAP plots showing the distribution of age groups in the single-nucleus multiome data prior to data filtering. **c**, UMAP plots showing the distribution of cells removed during data filtering. **d**, UMAP plots showing the expression levels of genes identified in the striatum (*ISL1* and *SIX3*), diencephalon (*OTX2* and *GBX2*), neuronal dendrites (*NRGN*), and oligodendrocyte processes (*MBP*). **e**, Classes, subclasses, and types identified from the snMultiome data.

**Extended Data Fig. 2.**
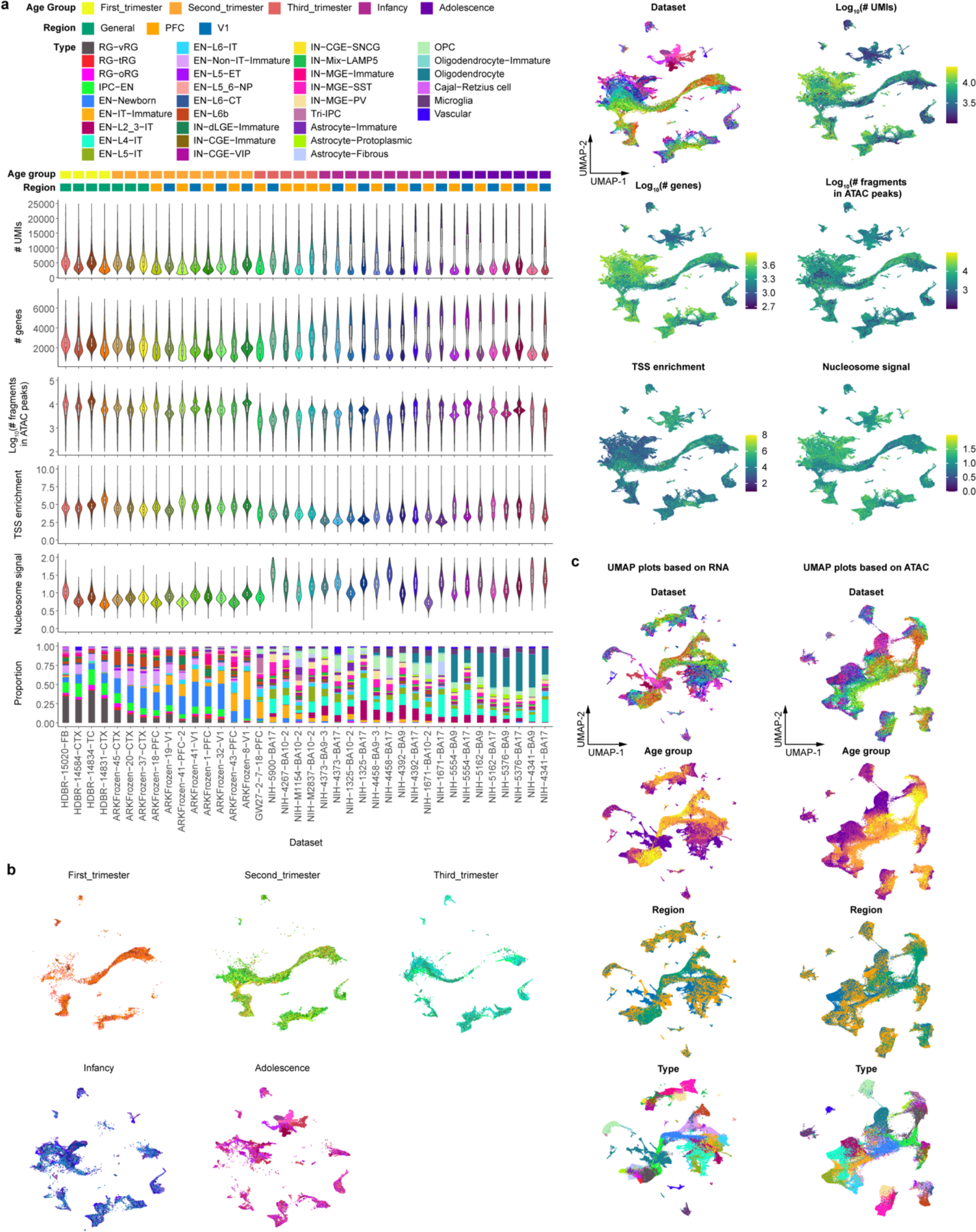
Quality control of the snMultiome data. **a**, Violin plots, box plots, barplots, and UMAP plots of several quality control metrics for evaluating the quality of individual samples in the snMultiome data, including numbers of unique molecular identifiers (# UMIs), numbers of identified genes (# genes), number of fragments in ATAC peaks, transcription start site (TSS) enrichment scores, and proportion of individual cell types in each sample. The legend for cell types can be found in panel b. **b** UMAP plots of cells from individual snMultiome datasets separated by age groups. **c**, UMAP plots generated based on RNA or ATAC data only. The legend can be found in panel a.

**Extended Data Fig. 3.**
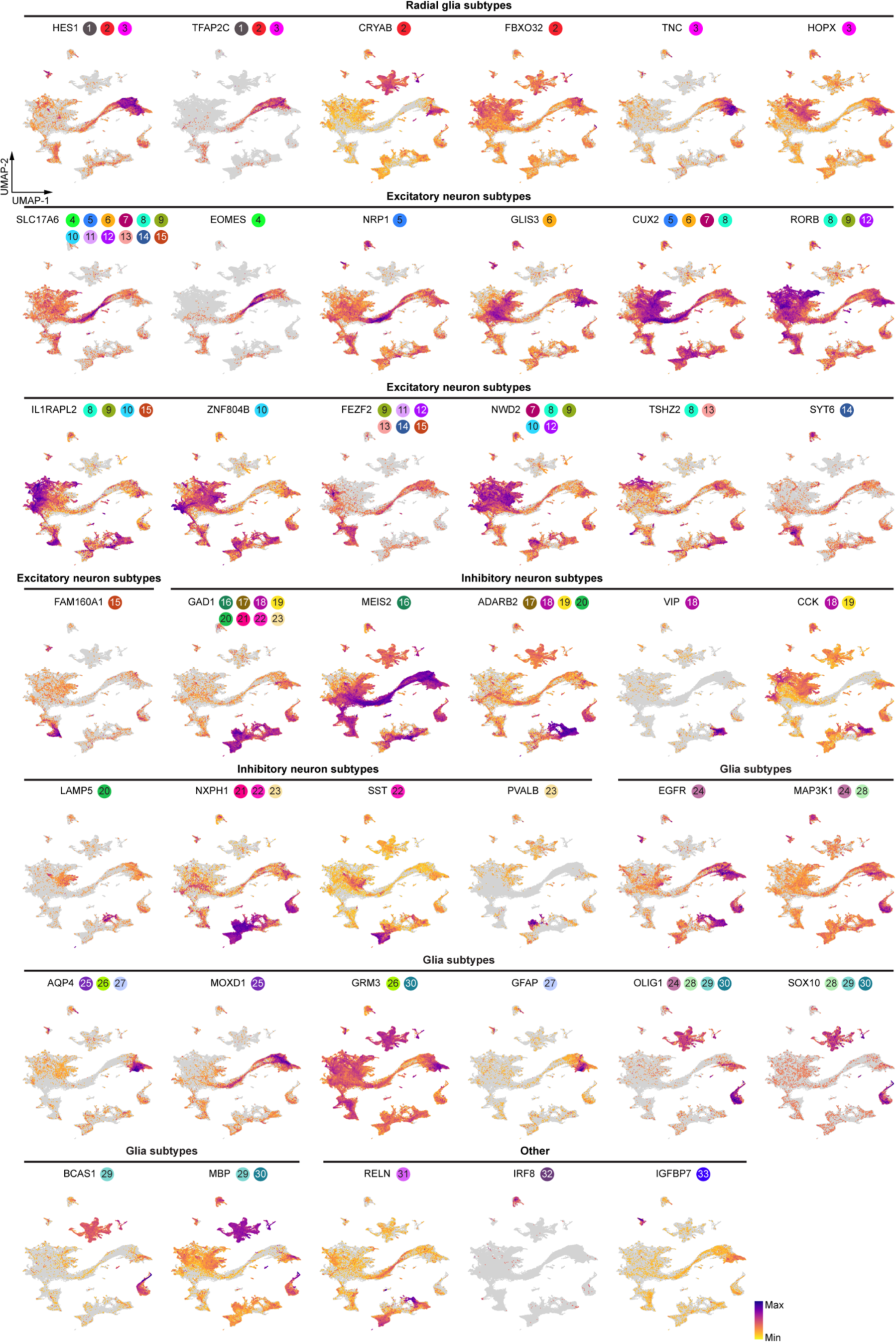
Expression patterns of marker genes in the single-nucleus multiome data. UMAP plots of all cells showing the expression levels of cell-type-specific marker genes. The colored circles and numbers pinpoint specific cell types where the gene is expressed. The legend for these numbers can be found in Fig. 1b.

**Extended Data Fig. 4.**
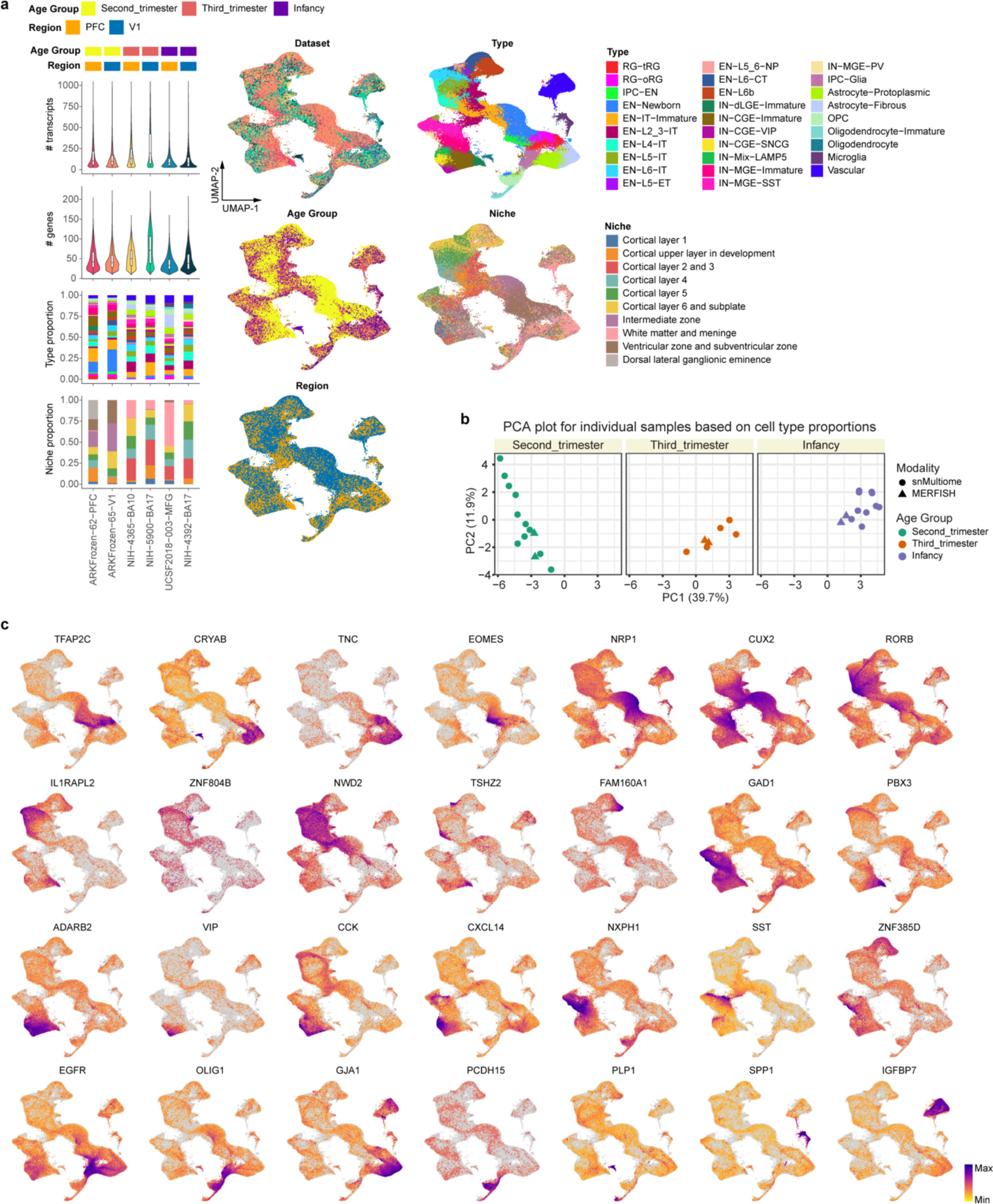
Quality control and annotation of MERFISH data. **a**, Violin plots, box plots, barplots, and UMAP plots of several metadata of MERFISH samples, including numbers of detected transcripts (# transcript), numbers of identified genes (# genes), age groups, regions, cell types, and niches. **b**, PCA plots based on cell type proportions for individual snMultiome and MERFISH samples in three different age groups. **c**, UMAP plots of all cells in the MERFISH dataset showing the expression levels of cell-type-specific marker genes.

**Extended Data Fig. 5.**
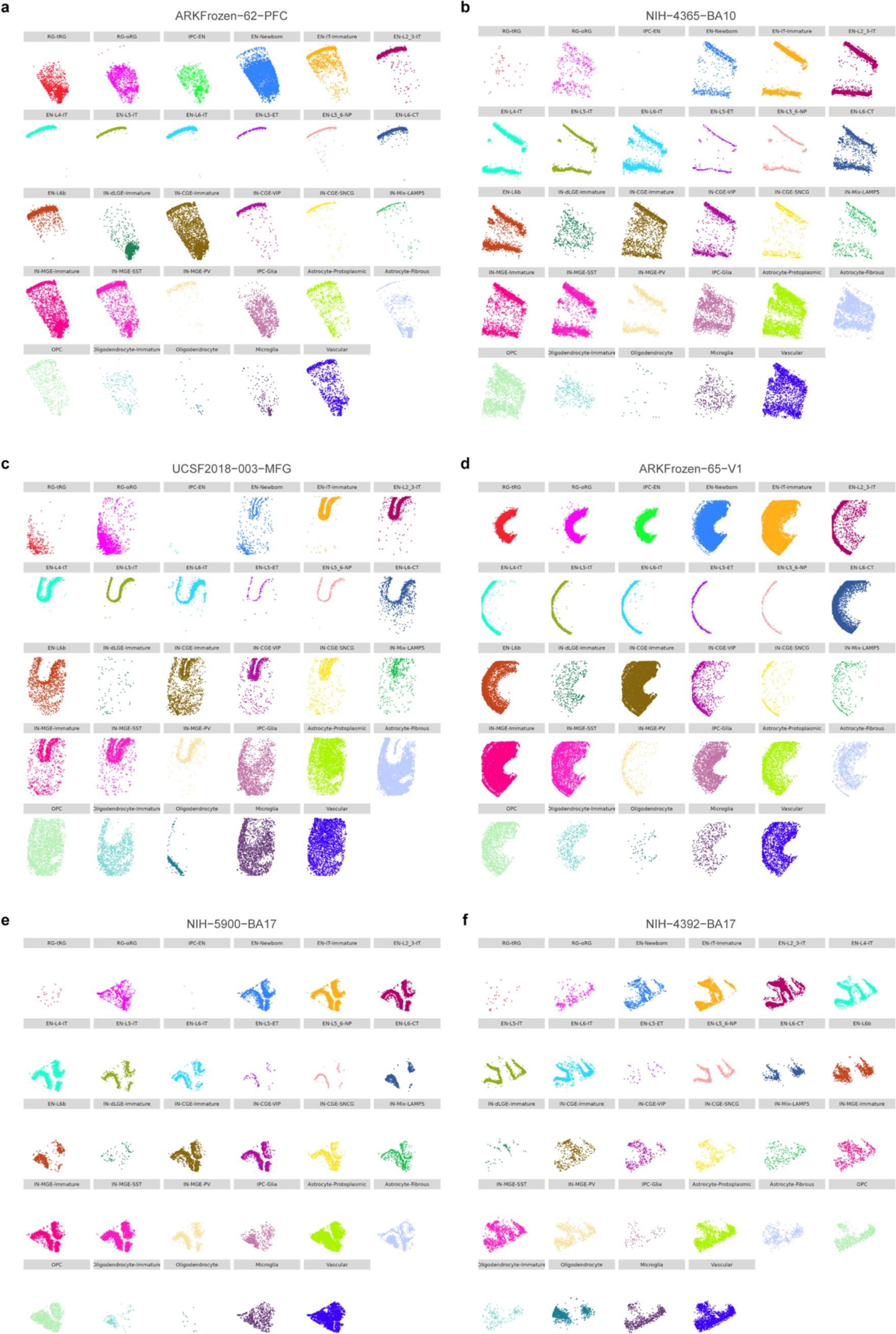
Spatial distribution of cell types in individual MERFISH samples.

**Extended Data Fig. 6.**
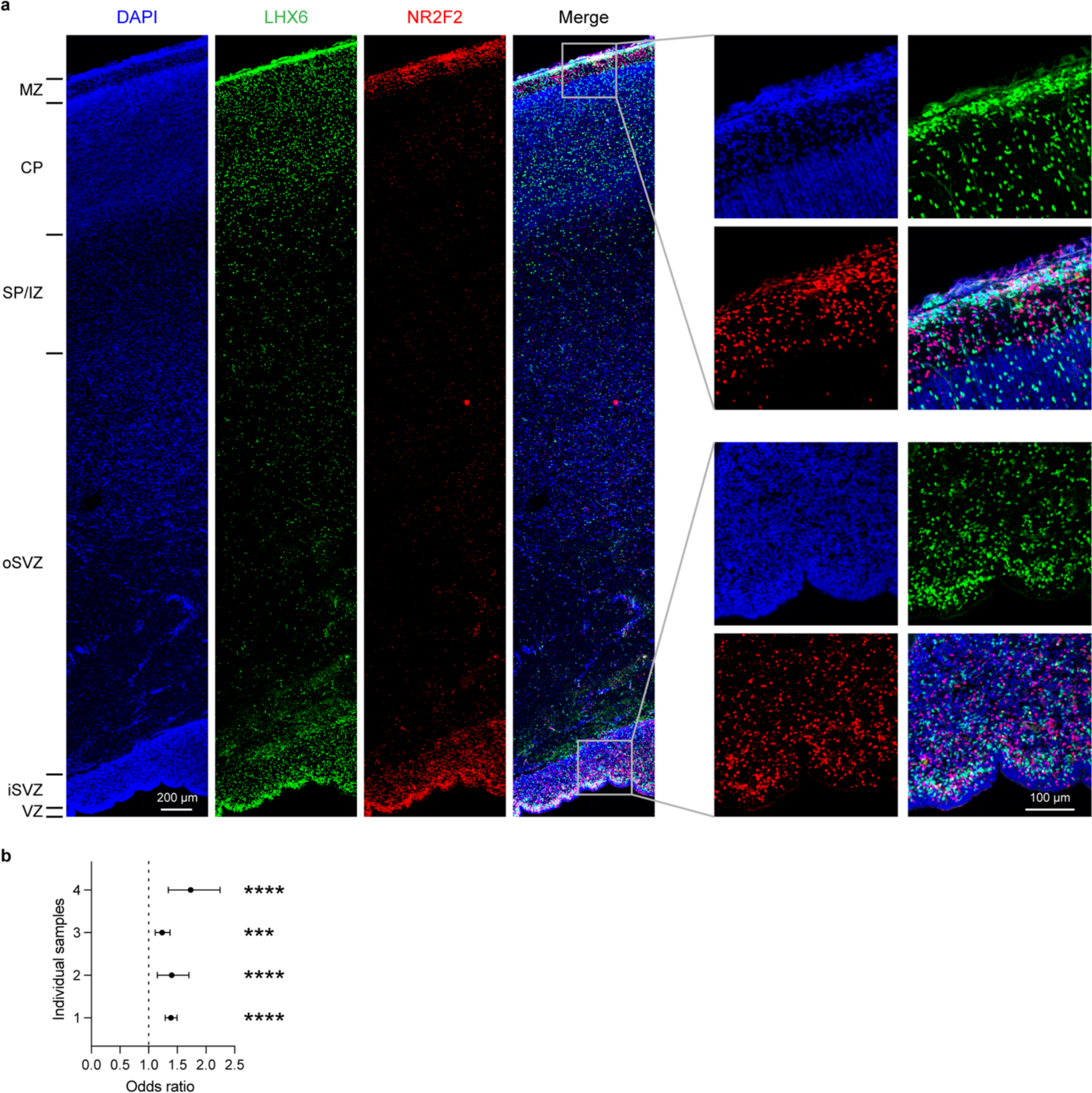
Difference in distribution of MGE- and CGE-derived interneurons in the second-trimester neocortex. **a**, Immunostaining of MGE-derived (LGX6^+^) and CGE-derived (NR2F2^+^) interneurons in the cortex of a gestational week (GW) 24 sample. MZ, marginal zone; CP, cortical plate; SP/IZ, subplate/intermediate zone; oSVZ, outer subventricular zone; iSVZ, inner subventricular zone; VZ, ventricular zone. **b**, Odds ratios of the number of CGE-derived interneurons in the MZ versus ventricular/subventricular zones relative to the number of MGE-derived interneurons. Data are presented as mean values with 95% confidence intervals. P values were obtained from two-sided Fisher’s exact test; ***P < 0.001, ****P < 0.0001.

**Extended Data Fig. 7.**
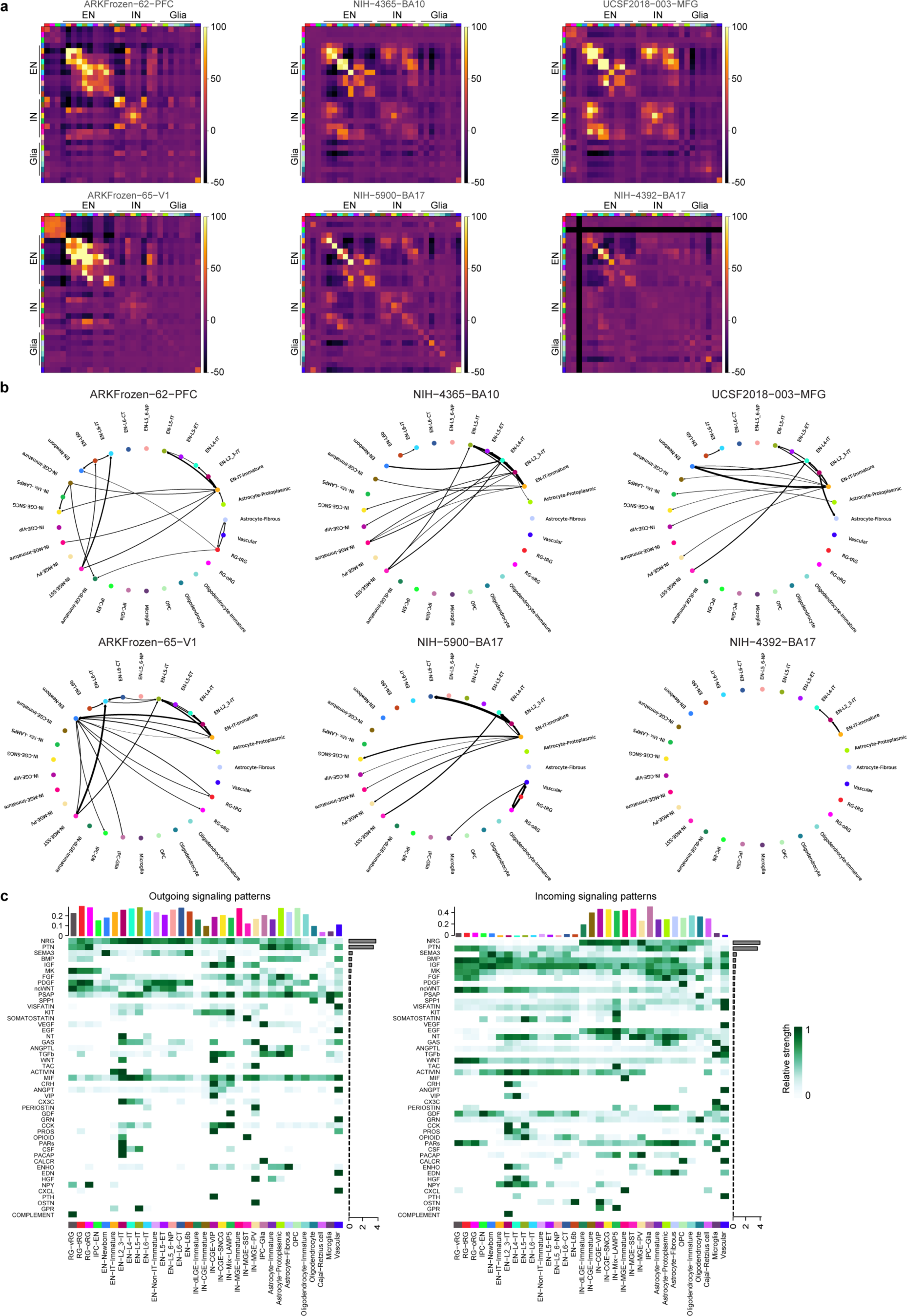
Intercellular communication between cell types in developing human cortex. **a**, Heatmaps showing neighborhood enrichment z scores of each MERFISH sample. The row and column annotations are color-coded by cell types, the legend of which can be found in Fig. 2a. When a particular cell type is not present in the dataset, the neighborhood enrichment z scores were arbitrarily set to −50. **b**, Circular maps showing significant intercellular communication determined by NCEM in each MERFISH sample. **c**, Heatmaps showing the relative strength of outgoing (left) and incoming (right) signaling pathways in individual cell types. The bar graphs on the top and right side of the heatmaps are the sum of communication probability (interaction strength) for each cell type and signaling pathway, respectively.

**Extended Data Fig. 8.**
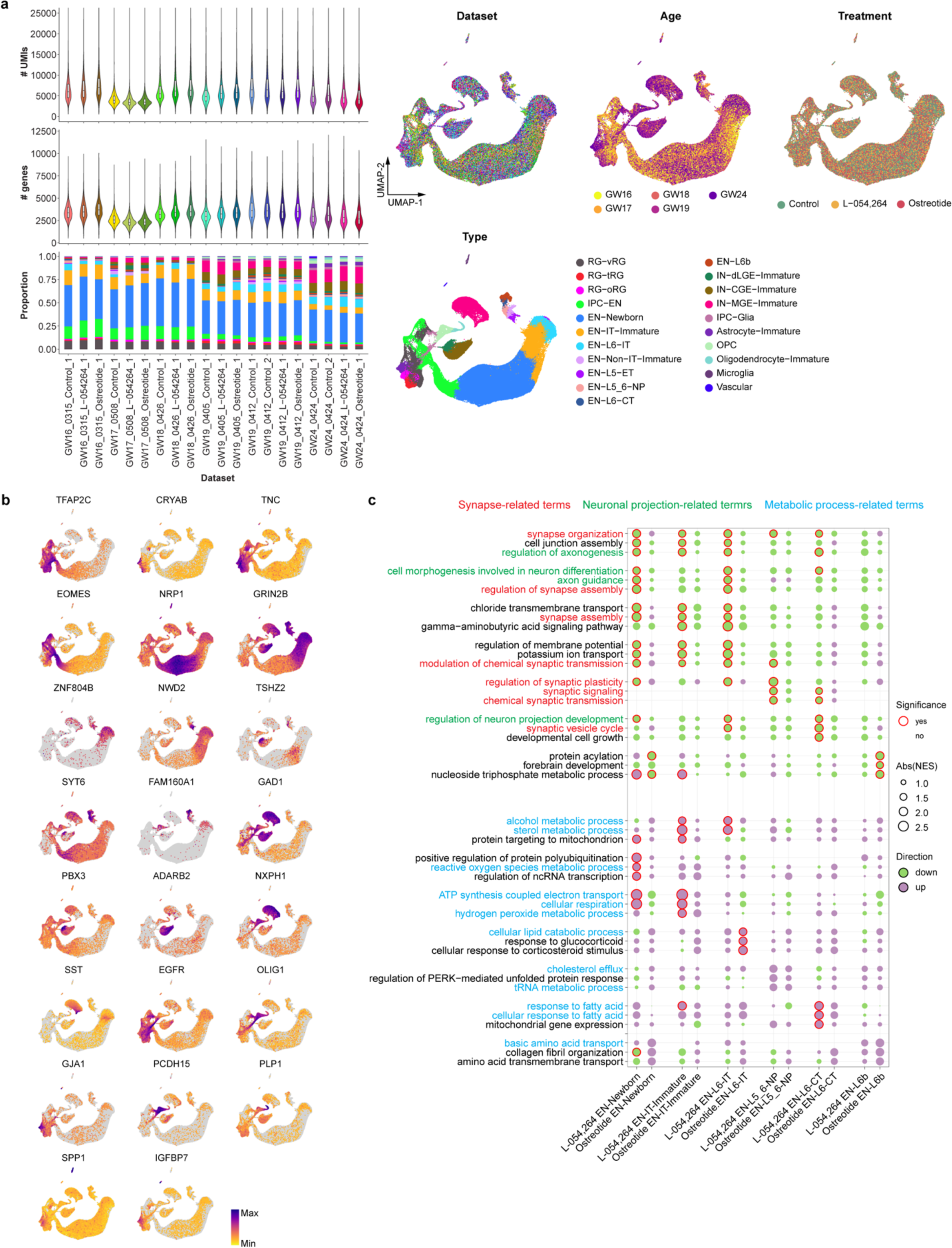
Effects of somatostatin on the transcriptome of excitatory neurons in the second-trimester human cortex. **a**, Violin plots, box plots, barplots, and UMAP plots of several metadata of scRNA-seq datasets from organotypic human brain slice cultures treated with and without somatostatin receptor agonists, including numbers of unique molecular identifiers (# UMIs), numbers of identified genes (# genes), ages, treatments, and cell types. **b**, UMAP plots of cells in the scRNA-seq dataset showing the expression levels of cell-type-specific marker genes. **c**, Gene set enrichment analysis (GSEA) highlighting the effects of L-054,264 and Ostreotide on different types of excitatory neurons. Significant terms, defined by Benjamini–Hochberg adjusted P values < 0.05, were outlined by a red circle. Abs(NES), absolute values of normalized enrichment scores.

**Extended Data Fig. 9.**
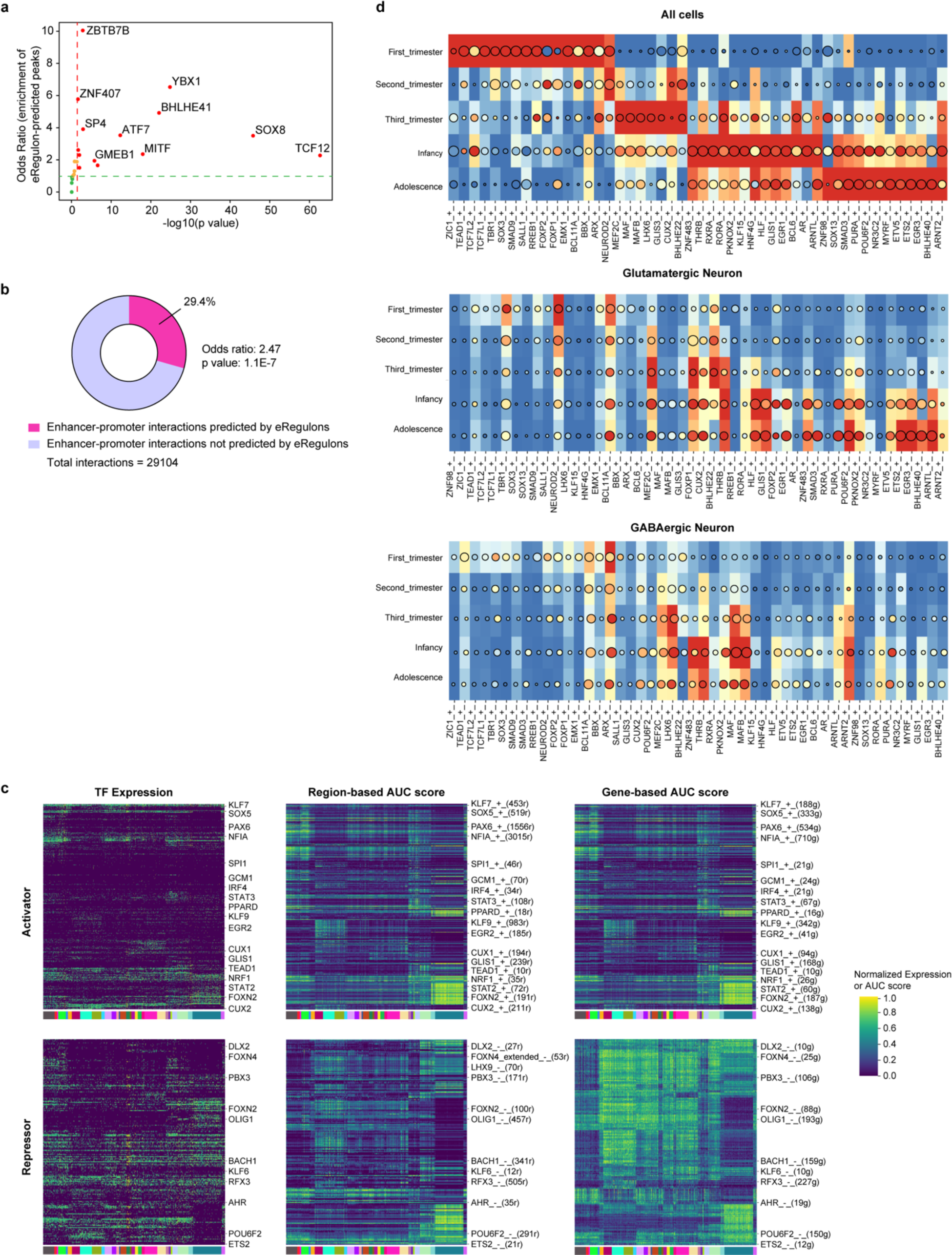
SCENIC+ identifies cell-type-specific eRegulons. **a**, Enrichment of eRegulon-predicted TF binding sites in ChIP-seq peaks from the human dorsolateral prefrontal cortex. P values were obtained from the two-sided Fisher’s exact test and adjusted using the Benjamini and Hochberg method. **b**, Overlap between eRegulon-predicted enhancer-promoter interactions and PLAC-seq loops from the developing human cortex. The P value was obtained from the two-sided Fisher’s exact test. **c**, Heatmaps showing the min-max normalized TF expression levels, region-based AUC scores, and gene-based AUC scores of activator eRegulons across cell types. **d**, Heatmap-dotplots showing the min-max normalized TF expression levels, region-based AUC scores, and gene-based AUC scores of selective eRegulons across age groups in all cells, Glutamatergic neurons, and GABAergic neurons.

**Extended Data Fig. 10.**
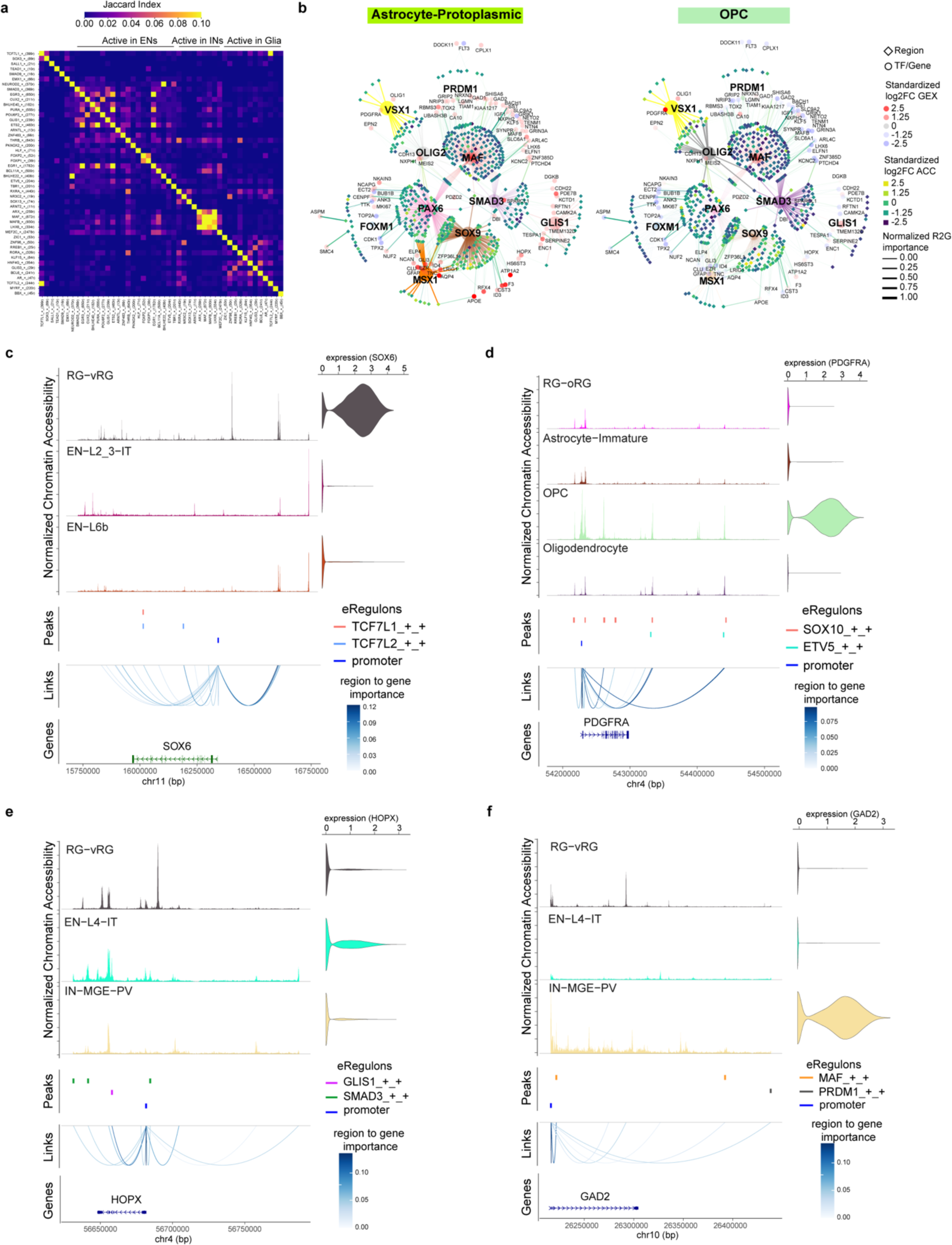
Cell-type-specific gene regulatory networks in the developing cortex. **a**, A heatmap showing Jaccard similarity matrix of target regions of cell-type-specific eRegulons listed in Fig. 3a. **b**, Gene regulatory networks of selective eRegulons in Astrocyte-Protoplasmics and OPCs. TF nodes and their links to enhancers are individually colored. The size and the transparency of the TF nodes represent their gene expression levels in each cell type. **c**, Coverage plots showing aggregated ATAC profiles across cell types on four genomic loci—*SOX6*, *PDGFRA*, *HOPX*, and *GAD2*. Identified candidate cis-regulatory elements (cCREs) are colored by their corresponding eRegulons. Region-to-gene links are shown as arcs and color-scaled based on region–gene importance scores obtained from SCENIC+ analysis.

**Extended Data Fig. 11.**
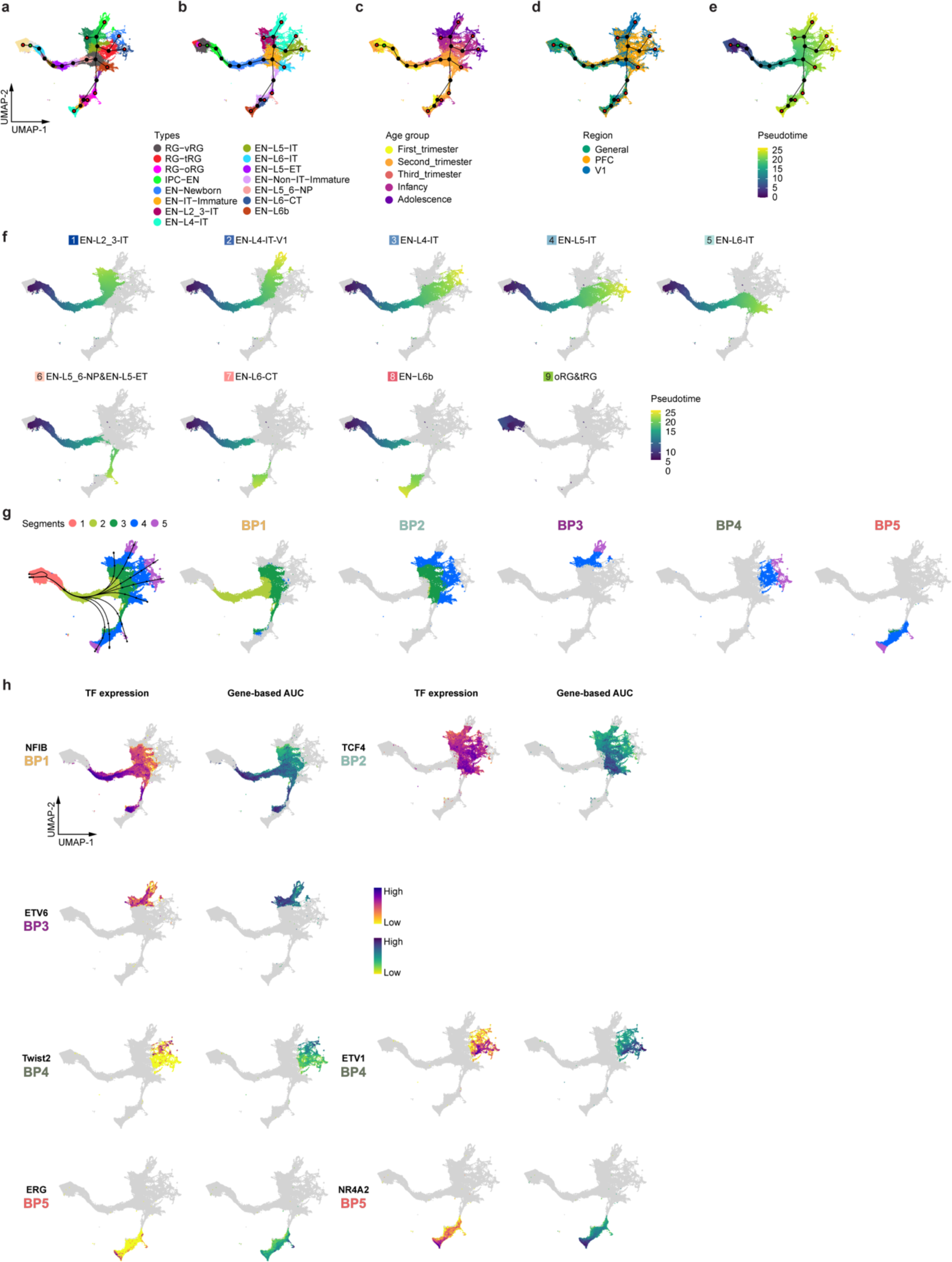
Differentiation trajectories of excitatory neuron lineages. **a**–**e**, UMAP plots of cells belonging to excitatory neuron lineages with clusters connected by a minimum spanning tree showing. The green node indicates the root node, and the red nodes indicate the ending nodes. Cells are color-coded by clusters (**a**), types (**b**), age groups (**c**), regions (**d**), or pseudotime (**e**). **f**, UMAP plots of each of the nine excitatory neuron lineages colored by pseudotime. **g**, UMAP plots of excitatory neuron lineages colored by the five pseudotime segments used for eRegulon activity analysis at bifurcation points. **h**, UMAP plots highlighting representative eRegulons involved in trajectory determination at bifurcation points.

**Extended Data Fig. 12.**
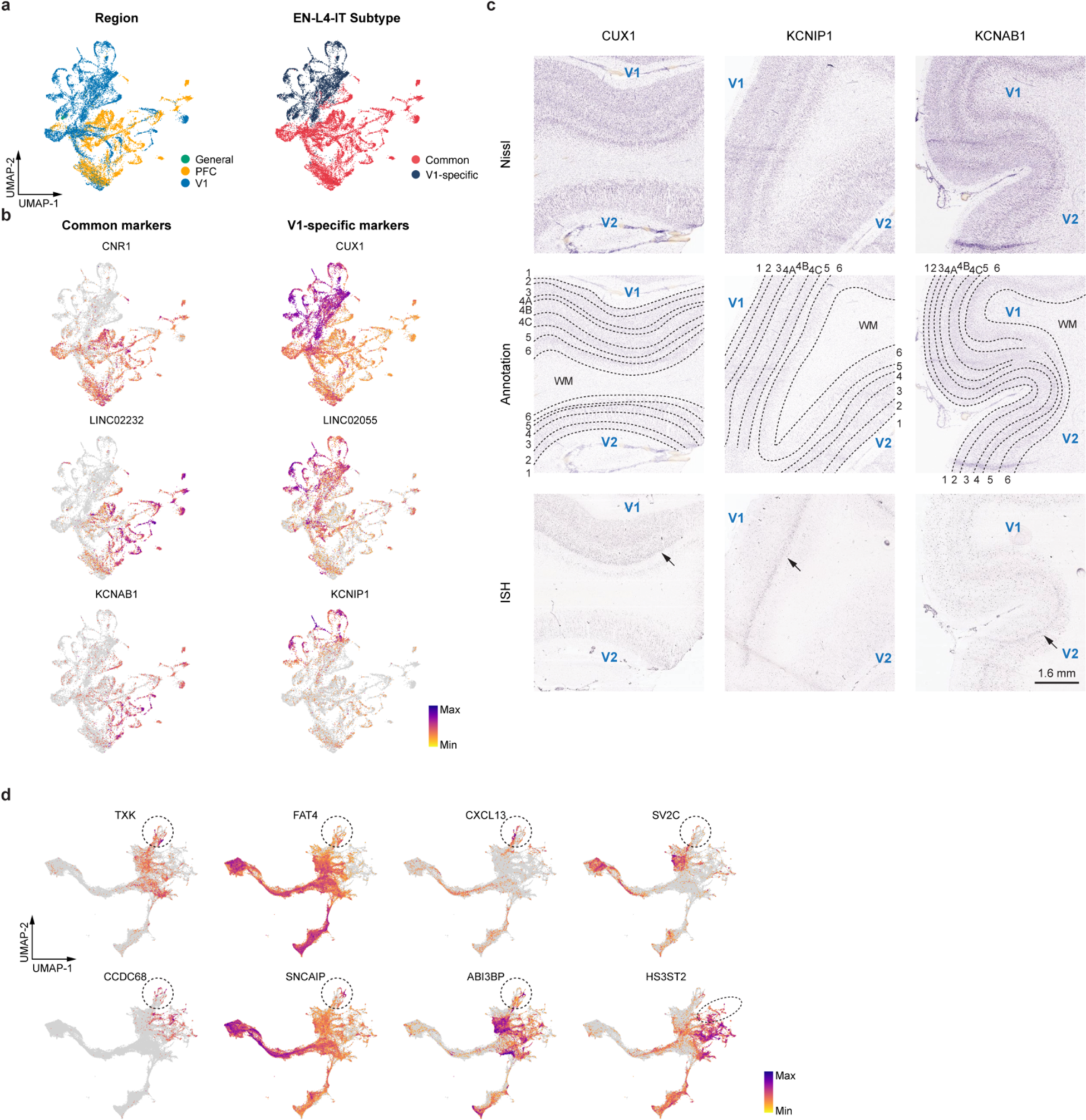
Markers of V1-specific EN-L4-IT subtype. **a**, UMAP plots of all EN-L4-IT color-coded by regions (left) and subtypes (right). **b**, UMAP plots showing the expression levels of representative differentially expressed genes between V1-specific and common EN-L4-IT neurons. **c**, In situ hybridization (ISH) of V1-biased (*CUX1* and *KCNIP1*), and common-biased genes in EN-L4-IT neurons in adult human V1 and V2 areas. **d**, UMAP plots of EN-L4-IT subtype marker genes found in adult human V1.

**Extended Data Fig. 13.**
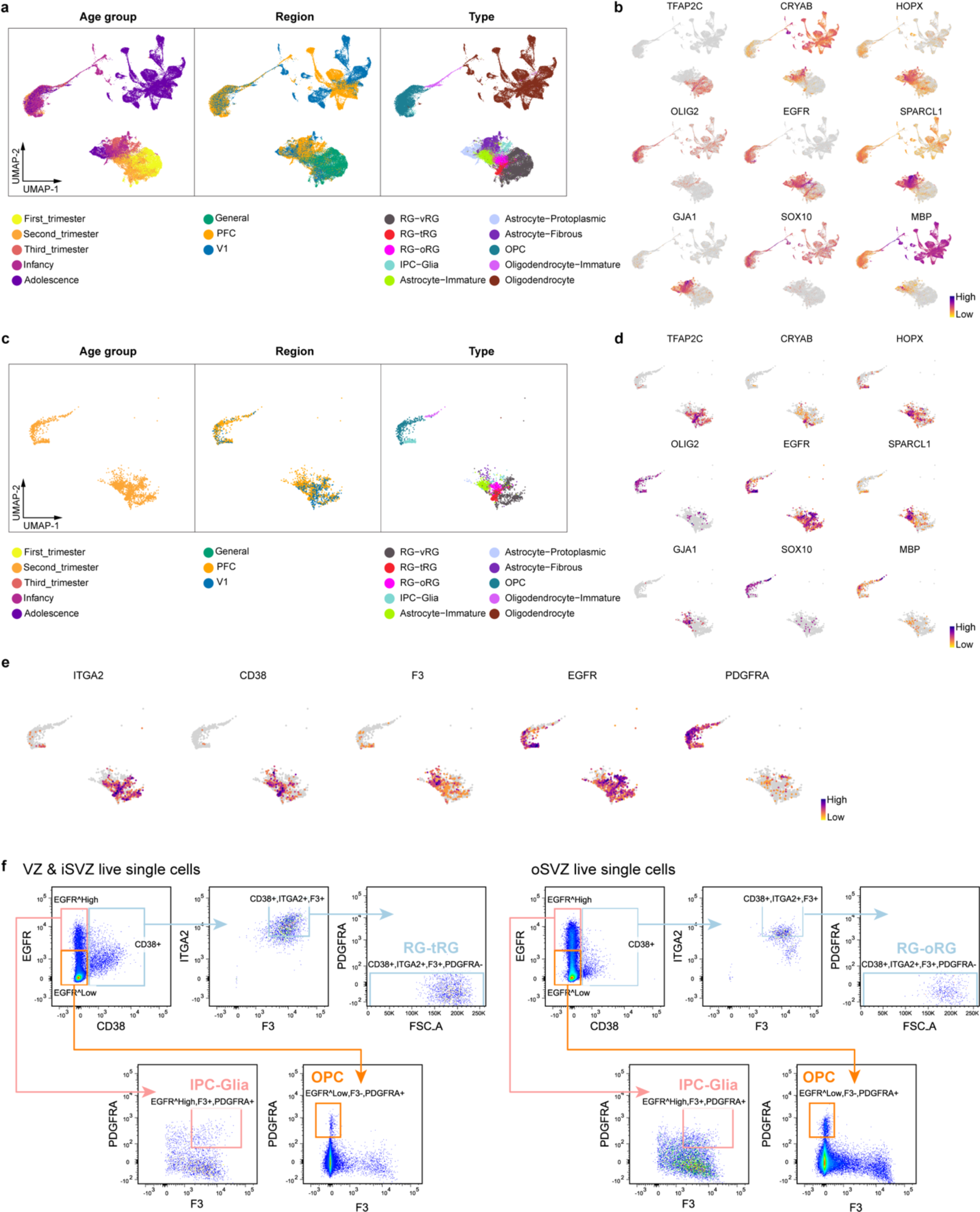
Markers of human glial cells and their isolation strategies. **a**, UMAP plots of cells belonging to glial lineages color-coded by age groups (left), regions (middle), and types (right). **b**, UMAP plots of cells belonging to glial lineages showing the expression levels of typical marker genes of individual cell types. **c**, UMAP plots of GW20 to GW23 cells belonging to glial lineages color-coded by age groups (left), regions (middle), and types (right). **d**, UMAP plots of GW20 to GW23 cells belonging to glial lineages showing the expression levels of typical marker genes of individual cell types. **e**, UMAP plots of GW20 to GW23 cells belonging to glial lineages showing the expression levels of surface markers used for glial progenitor isolation. **f**, Schematic of the sorting strategy for glial progenitors. VZ & iSVZ, ventricular zone and inner subventricular zone; oSVZ, outer subventricular zone.

**Extended Data Fig. 14.**
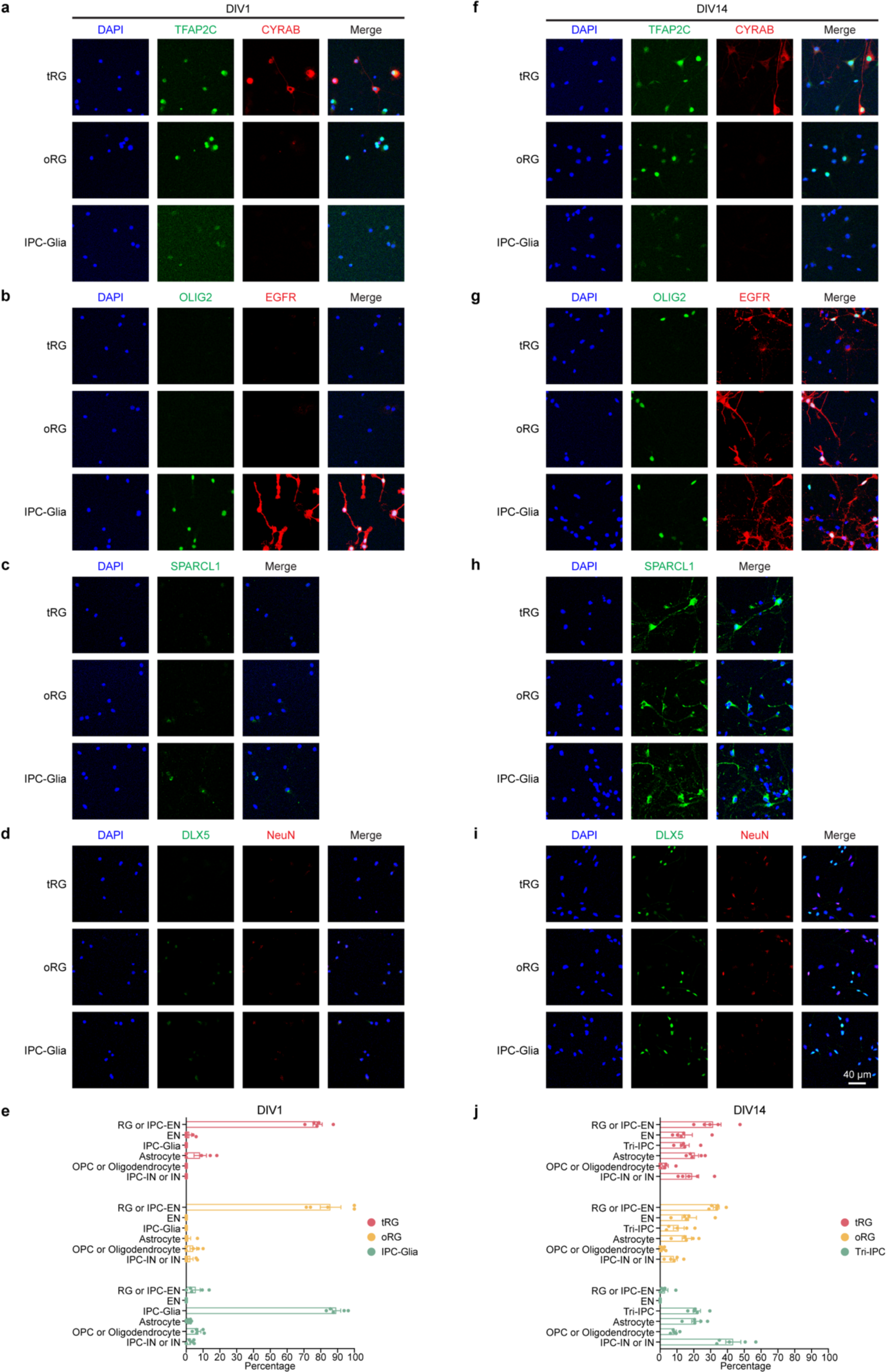
Immunostaining characterization of human glial progenitor differentiation. **a**–**d**, Immunostaining of isolated glial progenitors on days in vitro 1. **e**, Quantification of six cell types after sorting on days in vitro 1 (n = 5, 5, 5 samples), including RG or IPC-EN (TFAP2C^+^), IPC-Glia (OLIG2^+^EGFR^+^), OPC or oligodendrocyte (OLIG2^+^EGFR^−^), astrocyte (SPARCL1^+^), EN (NeuN^+^), and IPC-IN or IN (DLX5^+^). **f**–**i**, Immunostaining of progenies of glial progenitors on days in vitro 14. **j**, Quantification of six cell types after sorting on days in vitro 14 (n = 5, 5, 5 samples), including RG or IPC-EN (TFAP2C^+^), IPC-Glia (OLIG2^+^EGFR^+^), OPC or oligodendrocyte (OLIG2^+^EGFR^−^), astrocyte (SPARCL1^+^), EN (NeuN^+^), and IPC-IN or IN (DLX5^+^).

**Extended Data Fig. 15.**
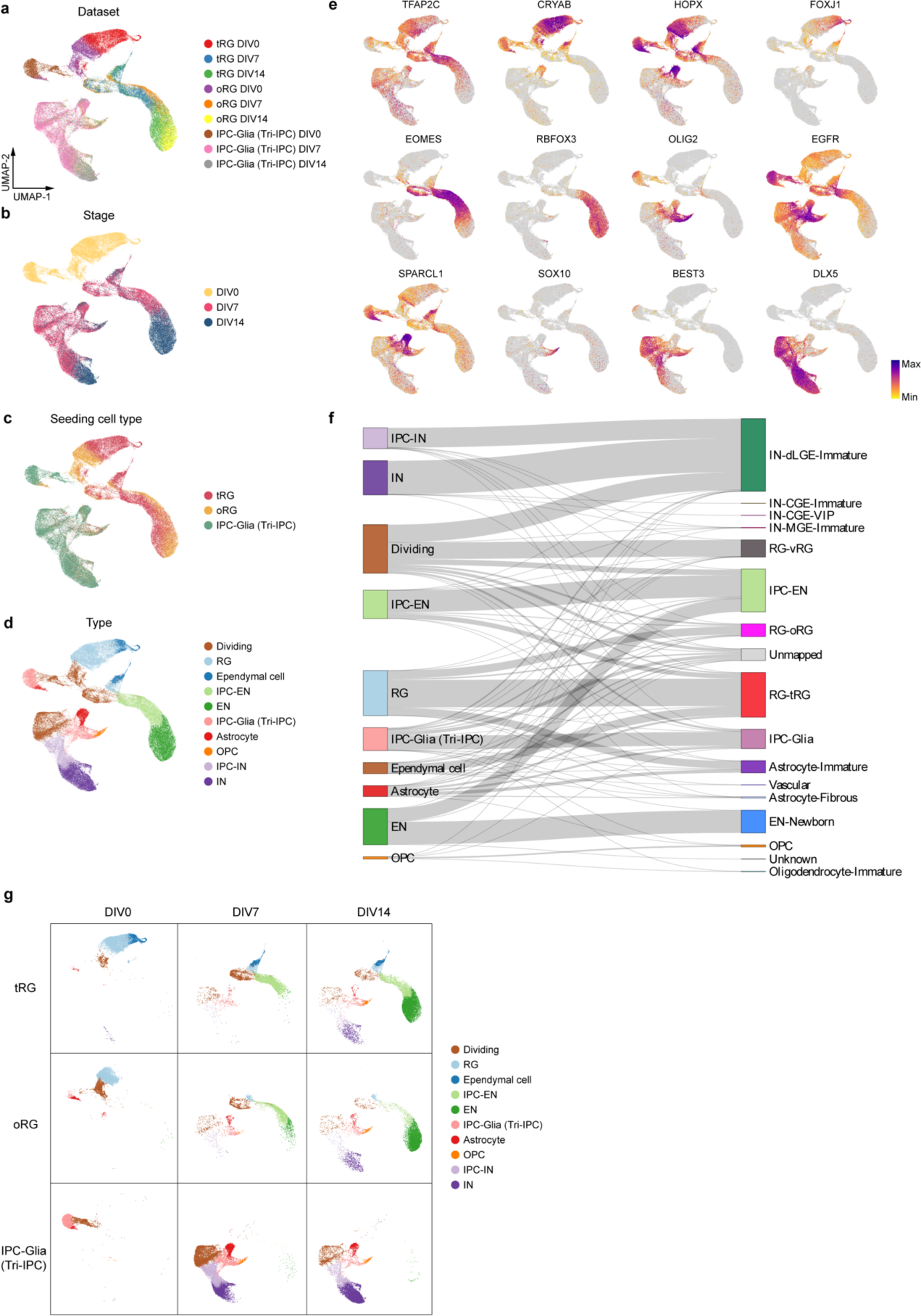
ScRNA-seq characterization of human glial progenitor differentiation. **a**–**d**, UMAP plots of isolated glial progenitors and their progenies during in vitro differentiation based on single-cell RNA sequencing data color-coded by datasets (**a**), stages (**b**), seeding cell types (**c**), and types (**d**). **e**, UMAP plots of isolated glial progenitors and their progenies showing the expression levels of typical marker genes of individual cell types. **f**, A Sankey plot showing the mapping of glial progenitors and their progenies to the snMultiome atlas by SingleCellNet. **g**, UMAP plots of isolated glial progenitors and their progenies separated by seeding cell types and stages.

**Extended Data Fig. 16.**
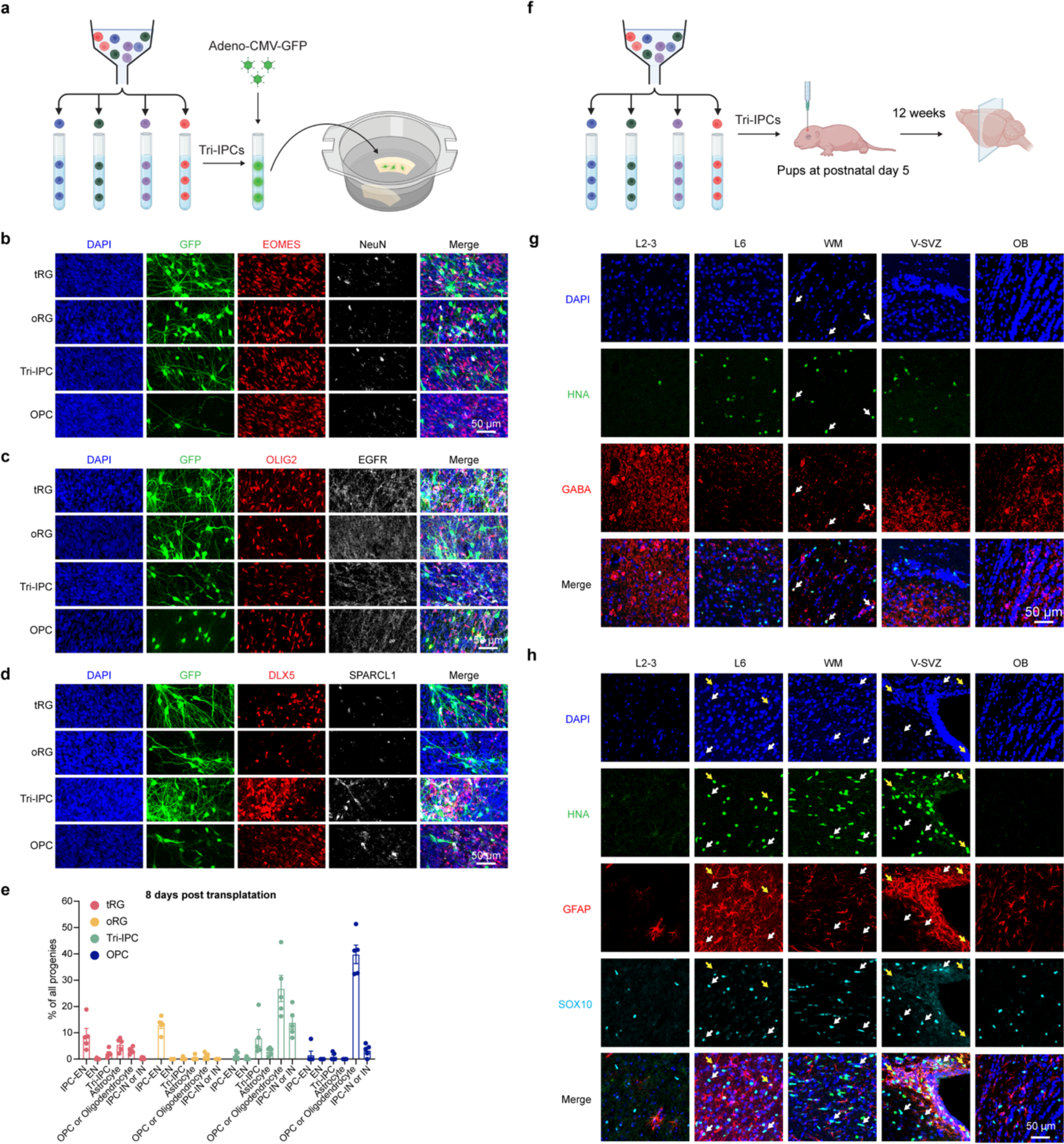
Lineage potential of human glial progenitors. **a**, Schematic of the slice transplantation assay for glial progenitors. **b**–**d**, Immunostaining of progenies after progenitor transplantation to acute cortical slices on days in vitro 8. **e**, Quantification of progeny types after progenitor transplantation to acute cortical slices (n = 5, 5, 5, 5 samples), including IPC-EN (EOMES^+^), EN (NeuN^+^), Tri-IPC (OLIG2^+^EGFR^+^), astrocyte (SPARCL1+), OPC or oligodendrocyte (OLIG2^+^EGFR^−^), and IPC-IN or IN (DLX5^+^). **f**, Schematic of the in vivo transplantation assay for glial progenitors. **g**, Immunostaining of progenies after progenitor in vivo transplantation into mouse cortex (n = 2 injections). White arrows indicate HNA^+^GABA^+^ inhibitory neurons. HNA, human nuclear antigen; L2-3, layer 2-3; L6, layer 6; WM, white matter; V-SVZ, ventricular-subventricular zone; OB, olfactory bulb. **h**, Immunostaining of progenies after progenitor in vivo transplantation into mouse cortex (n = 2 injections). White arrows indicate HNA^+^SOX10^+^ OPCs or oligodendrocytes. Yellow arrows indicate HNA^+^GFAP^+^ astrocytes. HNA, human nuclear antigen; L2-3, layer 2-3; L6, layer 6; WM, white matter; V-SVZ, ventricular-subventricular zone; OB, olfactory bulb.

**Extended Data Fig. 17.**
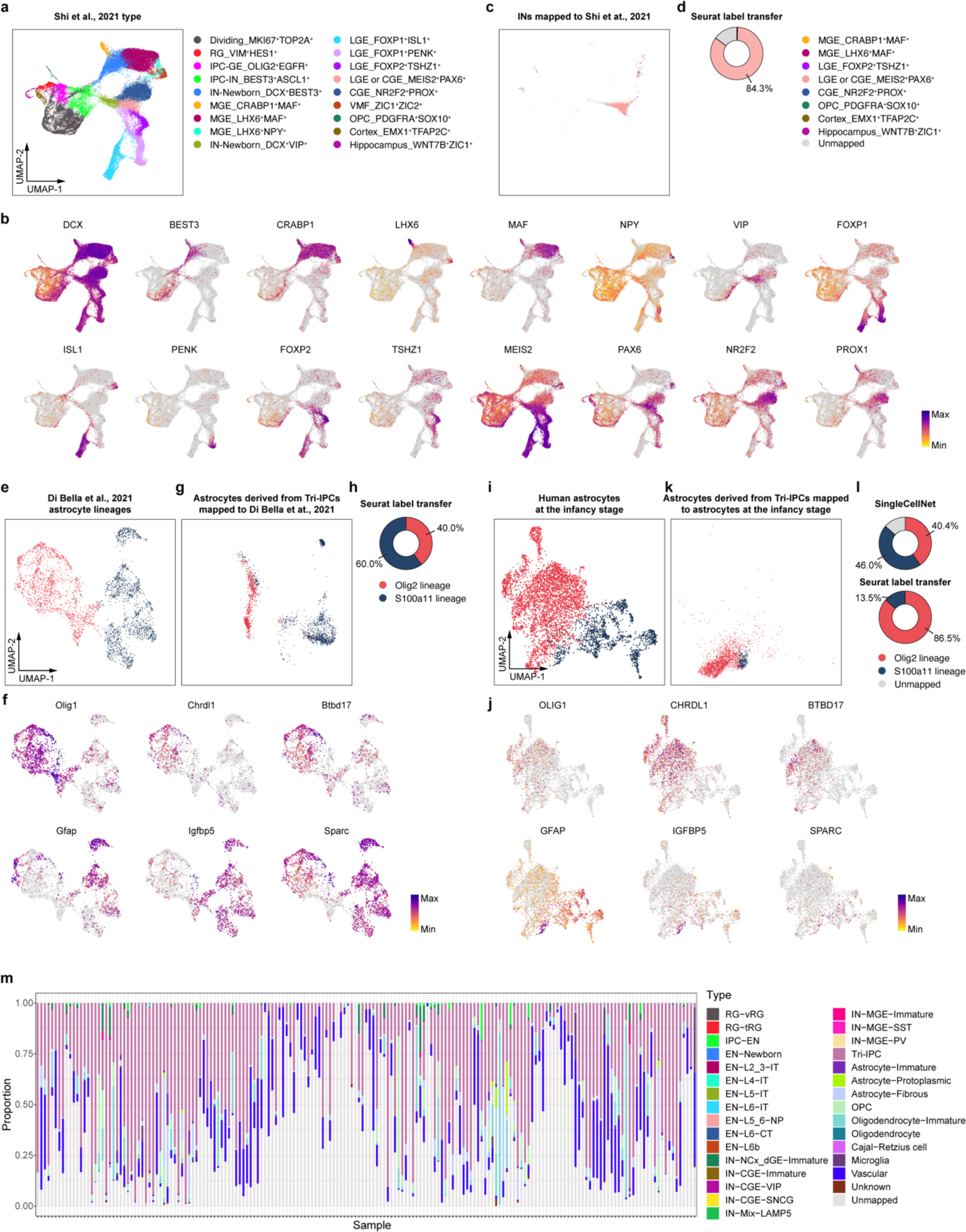
Mapping Tri-IPC progenies to reference data. **a**, UMAP plot of a reference human ganglionic eminence dataset^51^. Cells are color-coded by types. **b**, UMAP plots of human ganglionic eminence cells showing the expression levels of typical marker genes of individual cell types. **c**, UMAP plots of Tri-IPC-derived INs projected to the human ganglionic eminence dataset. Cells are color-coded by types and the legend can be found in panel d. **d**, Identities of Tri-IPC-derived INs mapped by Seurat label transfer. **e**, UMAP plot of mouse astrocytes from a reference developing mouse cortex dataset^55^. Cells are color-coded by lineages and the legend can be found in panel h. **f**, UMAP plots of the reference mouse astrocytes showing the expression levels of typical marker genes of individual astrocyte lineages. **g**, UMAP plots of Tri-IPC-derived astrocytes projected to the reference mouse astrocytes. Cells are color-coded by lineages and the legend can be found in panel h. **h**, Identities of Tri-IPC-derived astrocytes mapped by Seurat label transfer. **i**, UMAP plot of human astrocytes at the infancy stage. Cells are color-coded by lineages and the legend can be found in panel l. **j**, UMAP plots of human astrocytes showing the expression levels of typical marker genes of individual astrocyte lineages. **k**, UMAP plots of Tri-IPC-derived astrocytes projected to the reference human astrocytes. Cells are color-coded by lineages and the legend can be found in panel l. **l**, Identities of Tri-IPC-derived astrocytes predicted by SingleCellNet (top) or mapped by Seurat label transfer (bottom). **m**, Proportion of each SingleCellNet-predicted cell type across GBM samples.

**Extended Data Fig. 18.**
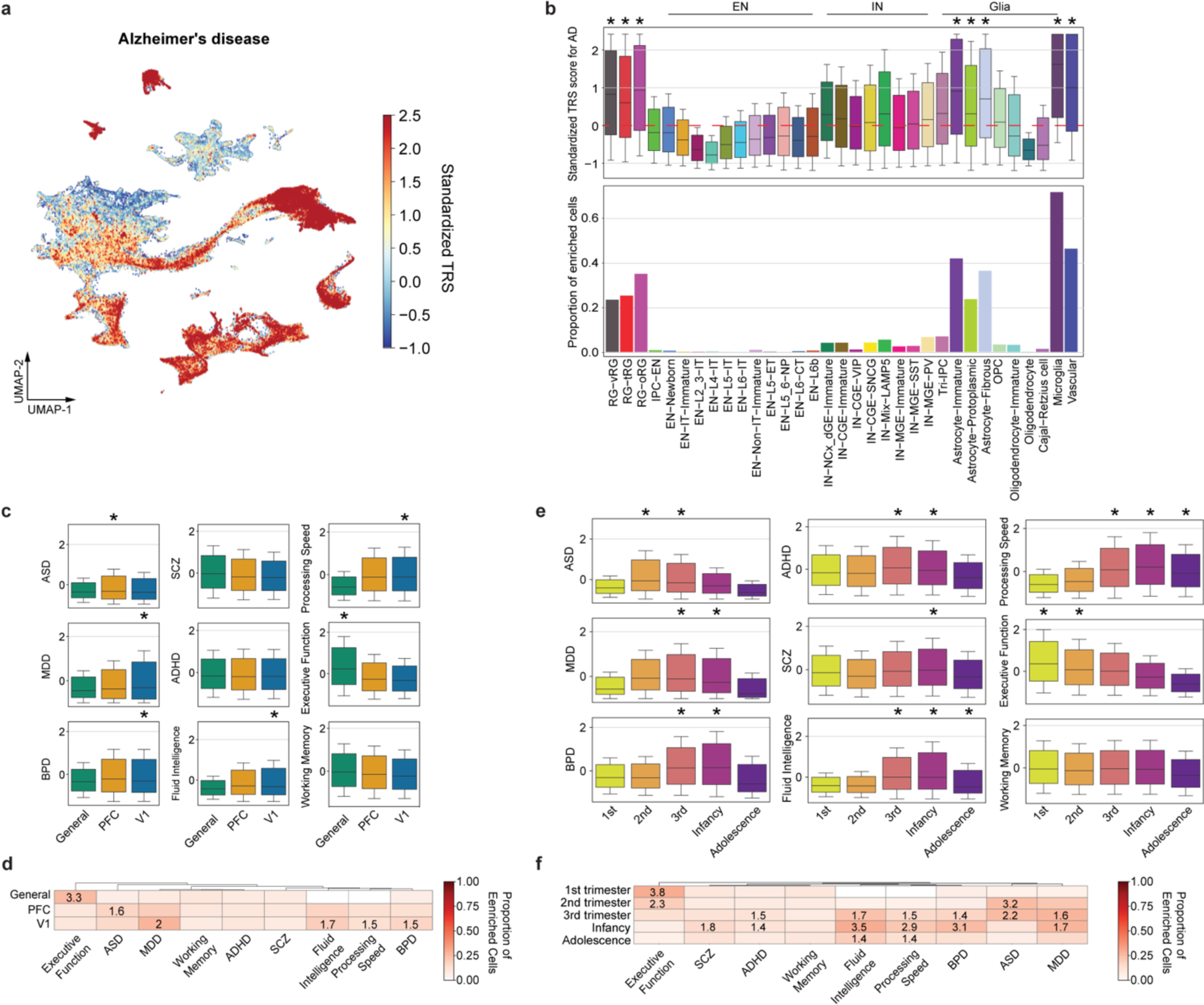
Neocortical cell association with human cognition and brain disorders. **a**, UMAP plot showing the standardized per-cell SCAVENGE trait relevance score (TRS) for Alzheimer’s disease. **b**, Top, boxplots showing the standardized SCAVENGE TRS for Alzheimer’s disease across cell types. Boxplot center: median; hinges: the 25th and 75th percentiles; whiskers: standard error. Bottom, bar plots showing the proportion of the cells with enriched trait relevance for Alzheimer’s disease across cell types. Two-sided hypergeometry test; *FDR < 0.01 & odds ratio > 1.4. **c**, Boxplots showing standardized SCAVENGE TRS for nine cognitive and disease traits across regions. Boxplot center: median; hinges: the 25th and 75th percentiles; whiskers: standard error. Two-sided hypergeometry test; *FDR < 0.01 & odds ratio > 1.4. **d**, Heatmap showing the proportion of the cells with enriched trait relevance across regions. Tiles with significant TRS enrichment (two-sided hypergeometric test, *FDR < 0.01 & odds ratio > 1.4) are annotated by their odd ratios. **e**, Boxplots showing standardized SCAVENGE TRS for nine cognitive and disease traits across developmental stages. Boxplot center: median; hinges: the 25th and 75th percentiles; whiskers: standard error. Two-sided hypergeometry test; *FDR < 0.01 & odds ratio > 1.4. **f**, Heatmap showing the proportion of the cells with enriched trait relevance across developmental stages. Tiles with significant TRS enrichment (two-sided hypergeometric test, *FDR < 0.01 & odds ratio > 1.4) are annotated by their odd ratios.

## Reference

1. Molnár, Z. et al. New insights into the development of the human cerebral cortex. J Anat 235, 432–451 (2019).

2. Long, H. K., Prescott, S. L. & Wysocka, J. Ever-Changing Landscapes: Transcriptional Enhancers in Development and Evolution. Cell 167, 1170–1187 (2016).

3. Nowakowski, T. J. et al. Spatiotemporal gene expression trajectories reveal developmental hierarchies of the human cortex. Science 358, 1318–1323 (2017).

4. Zhong, S. et al. A single-cell RNA-seq survey of the developmental landscape of the human prefrontal cortex. Nature 555, 524–528 (2018).

5. Li, M. et al. Integrative functional genomic analysis of human brain development and neuropsychiatric risks. Science 362, eaat7615 (2018).

6. Polioudakis, D. et al. A Single-Cell Transcriptomic Atlas of Human Neocortical Development during Mid-gestation. Neuron 103, 785–801.e8 (2019).

7. Fan, X. et al. Single-cell transcriptome analysis reveals cell lineage specification in temporal-spatial patterns in human cortical development. Sci Adv 6, eaaz2978 (2020).

8. Eze, U. C., Bhaduri, A., Haeussler, M., Nowakowski, T. J. & Kriegstein, A. R. Single-cell atlas of early human brain development highlights heterogeneity of human neuroepithelial cells and early radial glia. Nat Neurosci 24, 584–594 (2021).

9. Bhaduri, A. et al. An atlas of cortical arealization identifies dynamic molecular signatures. Nature 598, 200–204 (2021).

10. Ramos, S. I. et al. An atlas of late prenatal human neurodevelopment resolved by single-nucleus transcriptomics. Nat Commun 13, 7671 (2022).

11. Herring, C. A. et al. Human prefrontal cortex gene regulatory dynamics from gestation to adulthood at single-cell resolution. Cell 185, 4428–4447.e28 (2022).

12. Li, Y. et al. Spatiotemporal transcriptome atlas reveals the regional specification of the developing human brain. Cell 186, 5892–5909.e22 (2023).

13. Braun, E. et al. Comprehensive cell atlas of the first-trimester developing human brain. Science 382, eadf1226 (2023).

14. Velmeshev, D. et al. Single-cell analysis of prenatal and postnatal human cortical development. Science 382, eadf0834 (2023).

15. Ziffra, R. S., et al. Single-cell epigenomics reveals mechanisms of human cortical development. Nature 598, 205–213 (2021).

16. Trevino, A. E. et al. Chromatin and gene-regulatory dynamics of the developing human cerebral cortex at single-cell resolution. Cell 184, 5053–5069.e23 (2021).

17. van Bruggen, D. et al. Developmental landscape of human forebrain at a single-cell level identifies early waves of oligodendrogenesis. Dev Cell 57, 1421–1436.e5 (2022).

18. Zhu, K. et al. Multi-omic profiling of the developing human cerebral cortex at the single-cell level. Sci Adv 9, eadg3754 (2023).

19. Mannens, C. C. A. et al. Chromatin accessibility during human first-trimester neurodevelopment. Nature 2024 1–8 (2024) doi:10.1038/s41586-024-07234-1.

20. Hao, Y. et al. Integrated analysis of multimodal single-cell data. Cell 184, 3573–3587.e29 (2021).

21. Jorstad, N. L. et al. Transcriptomic cytoarchitecture reveals principles of human neocortex organization. Science 382, eadf6812 (2023).

22. Bishop, K. M., Goudreau, G. & O’Leary, D. D. M. Regulation of area identity in the mammalian neocortex by Emx2 and Pax6. Science 288, 344–349 (2000).

23. Chen, K. H., Boettiger, A. N., Moffitt, J. R., Wang, S. & Zhuang, X. Spatially resolved, highly multiplexed RNA profiling in single cells. Science 348, aaa6090 (2015).

24. Lim, L., Mi, D., Llorca, A. & Marín, O. Development and Functional Diversification of Cortical Interneurons. Neuron 100, 294–313 (2018).

25. Lim, L. et al. Optimization of interneuron function by direct coupling of cell migration and axonal targeting. Nature Neuroscience 2018 21:7 21, 920–931 (2018).

26. Stenman, J., Toresson, H. & Campbell, K. Identification of Two Distinct Progenitor Populations in the Lateral Ganglionic Eminence: Implications for Striatal and Olfactory Bulb Neurogenesis. Journal of Neuroscience 23, 167–174 (2003).

27. Akay, L. A., Effenberger, A. H. & Tsai, L. H. Cell of all trades: oligodendrocyte precursor cells in synaptic, vascular, and immune function. Genes Dev 35, 180–198 (2021).

28. Mittelbronn, M., Dietz, K., Schluesener, H. J. & Meyermann, R. Local distribution of microglia in the normal adult human central nervous system differs by up to one order of magnitude. Acta Neuropathol 101, 249–255 (2001).

29. Kriegstein, A. & Alvarez-Buylla, A. The Glial Nature of Embryonic and Adult Neural Stem Cells. Annu Rev Neurosci 32, 149–184 (2009).

30. Brill, M. S. et al. Adult generation of glutamatergic olfactory bulb interneurons. Nature Neuroscience 2009 12:12 12, 1524–1533 (2009).

31. Fischer, D. S., Schaar, A. C. & Theis, F. J. Modeling intercellular communication in tissues using spatial graphs of cells. Nature Biotechnology 2022 41:3 41, 332–336 (2022).

32. Jin, S. et al. Inference and analysis of cell-cell communication using CellChat. Nat Commun 12, 1–20 (2021).

33. Bravo González-Blas, C., et al. SCENIC+: single-cell multiomic inference of enhancers and gene regulatory networks. Nature Methods 2023 20:9 20, 1355–1367 (2023).

34. Loupe, J. M. et al. Multiomic profiling of transcription factor binding and function in human brain. Nat Neurosci 27, 1387–1399 (2024).

35. Song, M. et al. Cell-type-specific 3D epigenomes in the developing human cortex. Nature 587, 644–649 (2020).

36. Aibar, S. et al. SCENIC: Single-cell regulatory network inference and clustering. Nat Methods 14, 1083–1086 (2017).

37. Jolma, A. et al. DNA-dependent formation of transcription factor pairs alters their binding specificity. Nature 2015 527:7578 527, 384–388 (2015).

38. Wu, W. S. & Lai, F. J. Functional redundancy of transcription factors explains why most binding targets of a transcription factor are not affected when the transcription factor is knocked out. BMC Syst Biol 9, 1–9 (2015).

39. Street, K. et al. Slingshot: Cell lineage and pseudotime inference for single-cell transcriptomics. BMC Genomics 19, 1–16 (2018).

40. Van den Berge, K., et al. Trajectory-based differential expression analysis for single-cell sequencing data. Nat Commun 11, 1–13 (2020).

41. Li, Y. E. et al. A comparative atlas of single-cell chromatin accessibility in the human brain. Science 382, eadf7044 (2023).

42. Cadwell, C. R., Bhaduri, A., Mostajo-Radji, M. A., Keefe, M. G. & Nowakowski, T. J. Development and Arealization of the Cerebral Cortex. Neuron 103, 980–1004 (2019).

43. Huang, W. et al. Origins and Proliferative States of Human Oligodendrocyte Precursor Cells. Cell 182, 594–608.e11 (2020).

44. Fu, Y. et al. Heterogeneity of glial progenitor cells during the neurogenesis-to-gliogenesis switch in the developing human cerebral cortex. Cell Rep 34, 108788 (2021).

45. Yang, L., Li, Z., Liu, G., Li, X. & Yang, Z. Developmental Origins of Human Cortical Oligodendrocytes and Astrocytes. Neurosci Bull 38, 47–68 (2022).

46. Liu, D. D. et al. Purification and characterization of human neural stem and progenitor cells. Cell 186, 1179–1194.e15 (2023).

47. Weng, Q. et al. Single-Cell Transcriptomics Uncovers Glial Progenitor Diversity and Cell Fate Determinants during Development and Gliomagenesis. Cell Stem Cell 24, 707–723.e8 (2019).

48. Zhang, Y. et al. Cortical Neural Stem Cell Lineage Progression Is Regulated by Extrinsic Signaling Molecule Sonic Hedgehog. Cell Rep 30, 4490–4504.e4 (2020).

49. Li, X. et al. Decoding Cortical Glial Cell Development. Neurosci Bull 37, 440–460 (2021).

50. Andrews, M. G. et al. LIF signaling regulates outer radial glial to interneuron fate during human cortical development. Cell Stem Cell 30, 1382–1391.e5 (2023).

51. Shi, Y. et al. Mouse and human share conserved transcriptional programs for interneuron development. Science 374, eabj6641 (2021).

52. Schmitz, M. T. et al. The development and evolution of inhibitory neurons in primate cerebrum. Nature 603, 871–877 (2022).

53. Tan, Y. & Cahan, P. SingleCellNet: A Computational Tool to Classify Single Cell RNA-Seq Data Across Platforms and Across Species. Cell Syst 9, 207–213.e2 (2019).

54. Zhou, J. et al. Dual lineage origins of neocortical astrocytes. bioRxiv 2023.09.12.557313 (2023) doi:10.1101/2023.09.12.557313.

55. Di Bella, D. J. et al. Molecular logic of cellular diversification in the mouse cerebral cortex. Nature 595, 554–559 (2021).

56. Neftel, C. et al. An Integrative Model of Cellular States, Plasticity, and Genetics for Glioblastoma. Cell 178, 835–849.e21 (2019).

57. Couturier, C. P. et al. Single-cell RNA-seq reveals that glioblastoma recapitulates a normal neurodevelopmental hierarchy. Nature Communications 2020 11:1 11, 1–19 (2020).

58. Ruiz-Moreno, C. et al. Harmonized single-cell landscape, intercellular crosstalk and tumor architecture of glioblastoma. bioRxiv 2022.08.27.505439 (2022) doi:10.1101/2022.08.27.505439.

59. Albiach, A. M. et al. Glioblastoma is spatially organized by neurodevelopmental programs and a glial-like wound healing response. bioRxiv (2023) doi:10.1101/2023.09.01.555882.

60. Edwards, S. L., Beesley, J., French, J. D. & Dunning, M. Beyond GWASs: Illuminating the Dark Road from Association to Function. The American Journal of Human Genetics 93, 779–797 (2013).

61. Finucane, H. K. et al. Partitioning heritability by functional annotation using genome-wide association summary statistics. Nature Genetics 2015 47:11 47, 1228–1235 (2015).

62. Yu, F. et al. Variant to function mapping at single-cell resolution through network propagation. Nature Biotechnology 2022 40:11 40, 1644–1653 (2022).

63. Nott, A. et al. Brain cell type–specific enhancer–promoter interactome maps and disease-risk association. Science 366, 1134–1139 (2019).

64. Corces, M. R. et al. Single-cell epigenomic analyses implicate candidate causal variants at inherited risk loci for Alzheimer’s and Parkinson’s diseases. Nat Genet 52, 1158–1168 (2020).

65. Yang, A. C. et al. A human brain vascular atlas reveals diverse mediators of Alzheimer’s risk. Nature 603, 885–892 (2022).

66. Arranz, A. M. & De Strooper, B. The role of astroglia in Alzheimer’s disease: pathophysiology and clinical implications. Lancet Neurol 18, 406–414 (2019).

67. Abrahams, B. S. et al. SFARI Gene 2.0: A community-driven knowledgebase for the autism spectrum disorders (ASDs). Mol Autism 4, 1–3 (2013).

68. Harumi Yabuta, N. & Callaway, E. M. Functional Streams and Local Connections of Layer 4C Neurons in Primary Visual Cortex of the Macaque Monkey. Journal of Neuroscience 18, 9489–9499 (1998).

69. Krienen, F. M. et al. A marmoset brain cell census reveals regional specialization of cellular identities. Sci Adv 9, eadk3986 (2023).

70. Kohwi, M. et al. A Subpopulation of Olfactory Bulb GABAergic Interneurons Is Derived from Emx1- and Dlx5/6-Expressing Progenitors. Journal of Neuroscience 27, 6878–6891 (2007).

71. Young, K. M., Fogarty, M., Kessaris, N. & Richardson, W. D. Subventricular Zone Stem Cells Are Heterogeneous with Respect to Their Embryonic Origins and Neurogenic Fates in the Adult Olfactory Bulb. Journal of Neuroscience 27, 8286–8296 (2007).

72. Fuentealba, L. C. et al. Embryonic Origin of Postnatal Neural Stem Cells. Cell 161, 1644– 1655 (2015).

73. Marcy, G. et al. Single-cell analysis of the postnatal dorsal V-SVZ reveals a role for Bmpr1a signaling in silencing pallial germinal activity. Sci Adv 9, eabq7553 (2023).

74. Joyce, A. et al. Origin of GABAergic neurons in the human neocortex. Nature 417, 645– 649 (2002).

75. Zecevic, N., Hu, F. & Jakovcevski, I. Interneurons in the developing human neocortex. Dev Neurobiol 71, 18–33 (2011).

76. Hansen, D. V. et al. Non-epithelial stem cells and cortical interneuron production in the human ganglionic eminences. Nat Neurosci 16, 1576–1587 (2013).

77. Ma, T. et al. Subcortical origins of human and monkey neocortical interneurons. Nature Neuroscience 2013 16:11 16, 1588–1597 (2013).

78. Alzu’Bi, A. et al. The Transcription Factors COUP-TFI and COUP-TFII have Distinct Roles in Arealisation and GABAergic Interneuron Specification in the Early Human Fetal Telencephalon. Cerebral Cortex 27, 4971–4987 (2017).

79. Petanjek, Z., Berger, B. & Esclapez, M. Origins of Cortical GABAergic Neurons in the Cynomolgus Monkey. Cerebral Cortex 19, 249–262 (2009).

80. Wu, S. et al. Tangential migration and proliferation of intermediate progenitors of GABAergic neurons in the mouse telencephalon. Development 138, 2499–2509 (2011).

81. Delgado, R. N. et al. Individual human cortical progenitors can produce excitatory and inhibitory neurons. Nature 601, 397–403 (2021).

82. Dahmane, N. et al. The Sonic Hedgehog-Gli pathway regulates dorsal brain growth and tumorigenesis. Development 128, 5201–5212 (2001).

83. Ortega, J. A., Radonjić, N. V. & Zecevic, N. Sonic hedgehog promotes generation and maintenance of human forebrain Olig2 progenitors. Front Cell Neurosci 7, 62556 (2013).

84. Zhang, Y. et al. Cortical Neural Stem Cell Lineage Progression Is Regulated by Extrinsic Signaling Molecule Sonic Hedgehog. Cell Rep 30, 4490–4504.e4 (2020).

85. Tong, C. K. et al. A dorsal SHH-dependent domain in the V-SVZ produces large numbers of oligodendroglial lineage cells in the postnatal brain. Stem Cell Reports 5, 461–470 (2015).

86. Yu, Y. et al. Interneuron origin and molecular diversity in the human fetal brain. Nat Neurosci 24, 1745–1756 (2021).

87. Chung, C. et al. Cell-type-resolved somatic mosaicism reveals clonal dynamics of the human forebrain. bioRxiv 2023.10.24.563814 (2023) doi:10.1101/2023.10.24.563814.

88. Kim, S. N. et al. Cell lineage analysis with somatic mutations reveals late divergence of neuronal cell types and cortical areas in human cerebral cortex. bioRxiv 2023.11.06.565899 (2023) doi:10.1101/2023.11.06.565899.

89. Gaugler, T. et al. Most genetic risk for autism resides with common variation. Nature Genetics 2014 46:8 46, 881–885 (2014).

90. Grove, J. et al. Identification of common genetic risk variants for autism spectrum disorder. Nature Genetics 2019 51:3 51, 431–444 (2019).

91. Parikshak, N. N. et al. Integrative functional genomic analyses implicate specific molecular pathways and circuits in autism. Cell 155, 1008 (2013).

92. Willsey, A. J. et al. Coexpression Networks Implicate Human Midfetal Deep Cortical Projection Neurons in the Pathogenesis of Autism. Cell 155, 997–1007 (2013).

93. Velmeshev, D. et al. Single-cell genomics identifies cell type-specific molecular changes in autism. Science 364, 685–689 (2019).

94. Wang, L. & Kriegstein, A. Nuclei Isolation from Tissue for 10x Multiome by Iodixanol. protocol.io (2023) doi:10.17504/PROTOCOLS.IO.EQ2LYJ3NPLX9/V1.

95. Wolock, S. L., Lopez, R. & Klein, A. M. Scrublet: Computational Identification of Cell Doublets in Single-Cell Transcriptomic Data. Cell Syst 8, 281–291.e9 (2019).

96. Zhang, Y. et al. Model-based analysis of ChIP-Seq (MACS). Genome Biol 9, R137 (2008).

97. Amemiya, H. M., Kundaje, A. & Boyle, A. P. The ENCODE Blacklist: Identification of Problematic Regions of the Genome. Sci Rep 9, (2019).

98. Stuart, T., Srivastava, A., Madad, S., Lareau, C. A. & Satija, R. Single-cell chromatin state analysis with Signac. Nature Methods 2021 18:11 18, 1333–1341 (2021).

99. Choudhary, S. & Satija, R. Comparison and evaluation of statistical error models for scRNA-seq. Genome Biol 23, 27 (2022).

100. Butler, A., Hoffman, P., Smibert, P., Papalexi, E. & Satija, R. Integrating single-cell transcriptomic data across different conditions, technologies, and species. Nat Biotechnol 36, 411–420 (2018).

101. Waltman, L. & Van Eck, N. J. A smart local moving algorithm for large-scale modularity-based community detection. Eur Phys J B 86, 471 (2013).

102. Phipson, B. et al. propeller: testing for differences in cell type proportions in single cell data. Bioinformatics 38, 4720–4726 (2022).

103. Ritchie, M. E. et al. limma powers differential expression analyses for RNA-sequencing and microarray studies. Nucleic Acids Res 43, e47–e47 (2015).

104. Schep, A. N., Wu, B., Buenrostro, J. D. & Greenleaf, W. J. chromVAR: inferring transcription-factor-associated accessibility from single-cell epigenomic data. Nat Methods 14, 975–978 (2017).

105. Fornes, O. et al. JASPAR 2020: update of the open-access database of transcription factor binding profiles. Nucleic Acids Res 48, D87–D92 (2020).

106. Hao, Y. et al. Dictionary learning for integrative, multimodal and scalable single-cell analysis. Nat Biotechnol (2023) doi:10.1038/s41587-023-01767-y.

107. Nguyen, L. Van et al. Fast unfolding of communities in large networks. Journal of Statistical Mechanics: Theory and Experiment 2008, P10008 (2008).

108. Schindelin, J., et al. Fiji: An open-source platform for biological-image analysis. Nat Methods 9, 676–682 (2012).

109. Palla, G. et al. Squidpy: a scalable framework for spatial omics analysis. Nature Methods 2022 19:2 19, 171–178 (2022).

110. Germain, P. L., Robinson, M. D., Lun, A., Garcia Meixide, C. & Macnair, W. Doublet identification in single-cell sequencing data using scDblFinder. F1000Research 2022 10:979 10, 979 (2022).

111. Yu, G., Wang, L. G., Han, Y. & He, Q. Y. ClusterProfiler: An R package for comparing biological themes among gene clusters. OMICS 16, 284–287 (2012).

112. Hie, B., Cho, H., DeMeo, B., Bryson, B. & Berger, B. Geometric Sketching Compactly Summarizes the Single-Cell Transcriptomic Landscape. Cell Syst 8, 483–493.e7 (2019).

113. Bravo González-Blas, C., et al. cisTopic: cis-regulatory topic modeling on single-cell ATAC-seq data. Nat Methods 16, 397–400 (2019).

114. Janky, R. et al. iRegulon: From a Gene List to a Gene Regulatory Network Using Large Motif and Track Collections. PLoS Comput Biol 10, e1003731 (2014).

115. Van de Sande, B. et al. A scalable SCENIC workflow for single-cell gene regulatory network analysis. Nat Protoc 15, 2247–2276 (2020).

116. Scrucca, L., Fraley, C., Murphy, T. B. & Raftery, A. E. Model-Based Clustering, Classification, and Density Estimation Using Mclust in R. Model-Based Clustering, Classification, and Density Estimation Using mclust in R (CRC Press, 2023). doi:10.1201/9781003277965.

117. Rivellese, F., et al. Rituximab versus tocilizumab in rheumatoid arthritis: synovial biopsy-based biomarker analysis of the phase 4 R4RA randomized trial. Nat Med 28, 1256–1268 (2022).

118. Robinson, M. D., McCarthy, D. J. & Smyth, G. K. edgeR: A Bioconductor package for differential expression analysis of digital gene expression data. Bioinformatics 26, 139–140 (2009).

119. Merkle, F. T., Mirzadeh, Z. & Alvarez-Buylla, A. Mosaic organization of neural stem cells in the adult brain. Science (1979) 317, 381–384 (2007).

120. Ulirsch, J. C., et al. Interrogation of human hematopoiesis at single-cell and single-variant resolution. Nat Genet 51, 683–693 (2019).

