## Supplementary material for "Molecular and cellular dynamics of the developing human neocortex at single-cell resolution": SI Guide

**Supplementary Table 1 | snMultiome sample meta data and QC metrics.**

**Supplementary Table 2 | snMultiome cell-level metadata and QC metrics.**

**Supplementary Table 3 | Marker genes and cell type proportions in the snMultiome data.**

**Supplementary Table 4 | Transcription factor motif enrichment across cell types by chromVar.**

**Supplementary Table 5 | MERFISH sample meta data and panel information.**

**Supplementary Table 6 | MERFISH cell-level metadata.**

**Supplementary Table 7 | Node-centric expression model (NCEM) analysis results.**

**Supplementary Table 8 | CellChat analysis results.**

**Supplementary Table 9 | Fixed single-cell RNA-seq meta data and QC metrics.**

**Supplementary Table 10 | Somatostatin receptor agonist treatment scRNA-seq cell-level metadata and QC metrics.**

**Supplementary Table 11 | Differential gene expression analysis following somatostatin receptor agonist treatment.**

**Supplementary Table 12 | Gene set enrichment analysis of sample treated with somatostatin receptor agonists.**

**Supplementary Table 13 | Target regions and genes of SCENIC+ eRegulons.**

**Supplementary Table 14 | Validation of eRegulons by ChIP-seq and H3K4me3 PLAC-seq data.**

**Supplementary Table 15 | Transcription factor expression, region-based AUC, and gene-based AUC of eRegulons.**

**Supplementary Table 16 | Slingshot trajectory and pseudotime.**

**Supplementary Table 17 | eRegulon modules and gene ontology enrichment.**

**Supplementary Table 18 | Differentially expressed genes between V1-specific and common EN-L4-IT cells.**

**Supplementary Table 19 | TradeSeq analysis results at bifurcation points.**

**Supplementary Table 20 | Glial progenitor culture scRNA-seq sample meta data and QC metrics.**

**Supplementary Table 21 | Glial progenitor culture scRNA-seq cell-level metadata and QC metrics.**

**Supplementary Table 22 | Clonal analysis of isolated glial progenitors.**

**Supplementary Table 23 | SCAVENGE analysis results summarized across cell types.**

**Supplementary Table 24 | SCAVENGE analysis results summarized across regions and age groups.**

**Supplementary Table 25 | Spearman Correlation between eRegulon (activators) gene-based AUC with SCAVENGE TRS.**
